# Identification of Siderophores with Unexpected Antibacterial Properties from *Actinoplanes teichomyceticus*

**DOI:** 10.64898/2026.03.19.712980

**Authors:** Abiodun S. Oyedele, Somnath Jana, KyuOk Jeon, Navin Vazrala, Donald F. Stec, Kwangho Kim, Gary A. Sulikowski, Allison S. Walker

## Abstract

*Actinoplanes teichomyceticus* is a well-established producer of bioactive secondary metabolites, including the glycopeptide antibiotic teicoplanin. Although its antibiotic biosynthetic capacity has been extensively investigated, its siderophore diversity and any additional biological functions of these iron-chelating metabolites remain comparatively underexplored. We identified a reproducibly bioactive, teicoplanin-independent fraction that inhibited *Bacillus spizizenii*. Molecular networking applied to this fraction identified hydroxamate ferrioxamine and desferrioxamine-type siderophores as the dominant metabolites, including acylated analogs detected as Al^3+^- and Fe^3+^-chelated species. Robust siderophore secretion was confirmed by the CAS assay. Notably, siderophore-enriched fractions exhibited selective antibacterial activity against Gram-positive bacteria, with minimum inhibitory concentrations of approximately 16 µg/mL against *B. spizizenii* and partial inhibition of *Staphylococcus aureus*, while no activity was observed against *Escherichia coli*. Synthetic C7 and C9 acyl-desferrioxamine analogs showed enhanced antibacterial activity upon Al³ chelation, indicating a metal-dependent bioactivity. These findings reveal an unexpected antibacterial role for ferrioxamine-type siderophores produced by *A. teichomyceticus*, extending their function beyond iron acquisition, possibly through a “Trojan horse” (or “Trojan metal”) mechanism.

## Introduction

Microorganisms such as bacteria and fungi rely on siderophores, high-affinity ferric iron (Fe^3+^) chelators, to scavenge iron in limiting environments, shaping microbial ecology and interspecies competition. The general mechanism involves the extracellular sequestration of ferric ions by the siderophore, followed by the active transport of the resultant iron-siderophore complex into the cell through specific outer membrane receptors, where the iron is then released for metabolic use.^1–3^ Siderophores exhibit remarkable chemical diversity and are broadly classified based on the functional groups that coordinate the ferric ion: primarily catecholates (or phenolates), hydroxamates, and carboxylates, with many “mixed-type” structures also known, and their biosynthesis involves either modular NRPS enzymes or NRPS-independent siderophore (NIS) synthetases of the IucA/IucC superfamily.^4,5^ Siderophores have exceptional affinity for Fe³, with formation constants greater than 10^30^, as is seen for the catecholate siderophore enterobactin.^1^ Beyond their primary role in iron homeostasis, additional biological functions of siderophores have recently been discovered. They are now recognized as key players in microbial ecology, acting as virulence factors for pathogenic bacteria, mediators of biofilm formation, agents of interspecies signaling, and potent biocontrol agents against phytopathogens.^6–8^ Siderophores can have antibacterial activity by depriving competing microbes of essential iron in low-iron environments.^2^ They can also be engineered to have direct toxic effects by conjugating with antibiotics in a “Trojan horse” strategy.^9^ Recent studies have explored siderophores as tools for targeted antimicrobial delivery, enhancing the efficacy of existing drugs against resistant pathogens.^7,10^

*Actinoplanes teichomyceticus* is a rare Gram-positive, spore-forming actinomycete originally isolated from a soil sample in India.^11^ This bacterium is well understood as the producer of teicoplanin, a complex glycopeptide antibiotic that is a “last resort” drug for treating severe infections caused by multidrug-resistant Gram-positive pathogens, such as *methicillin-resistant Staphylococcus aureus* (MRSA).^12,13^ Despite decades of intensive research aimed at optimizing teicoplanin fermentation and genetically engineering the producing strain for enhanced yields of derivatives,^14,15^ teicoplanin and teichomycin A1^16^ remain the major characterized natural products produced by *A. teichomyceticus*.^16^ However, genome analyses of *A. teichomyceticus* ATCC 31121 using antiSMASH 5.0 reveals a broader biosynthetic potential encompassing additional natural products and unexplored secondary metabolites^17^ (Fig. 1), containing multiple BGCs predicted by our machine learning genome mining pipeline^18^ to produce antibacterial compounds and multiple putative siderophore-producing BGCs (Fig. 2). We hypothesize that the predicted antibacterial and siderophore-producing BGCs are either “cryptic” or that their products were previously overlooked by researchers focused only on teicoplanin.

**Fig. 1.**
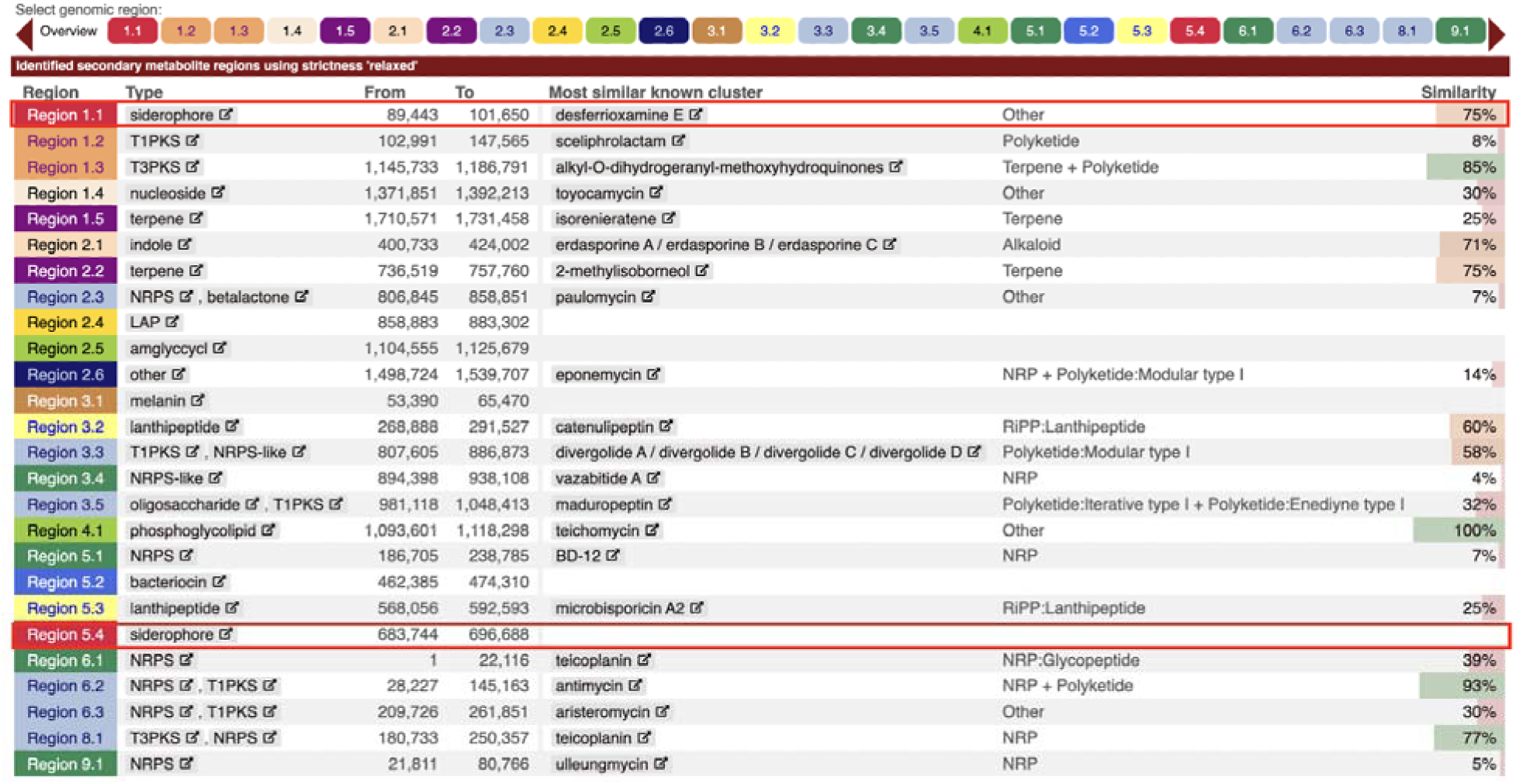
antiSMASH 5.1.1 analysis (strictness: relaxed) of *Actinoplanes teichomyceticus* DSM 43866 showing predicted secondary metabolite biosynthetic gene clusters. Regions encoding siderophore biosynthesis are highlighted in red. Teicoplanin-associated BGCs were identified in regions 6.1 and 8.1, while the teichomycin BGC was detected in region 4.1.

**Fig. 2.**
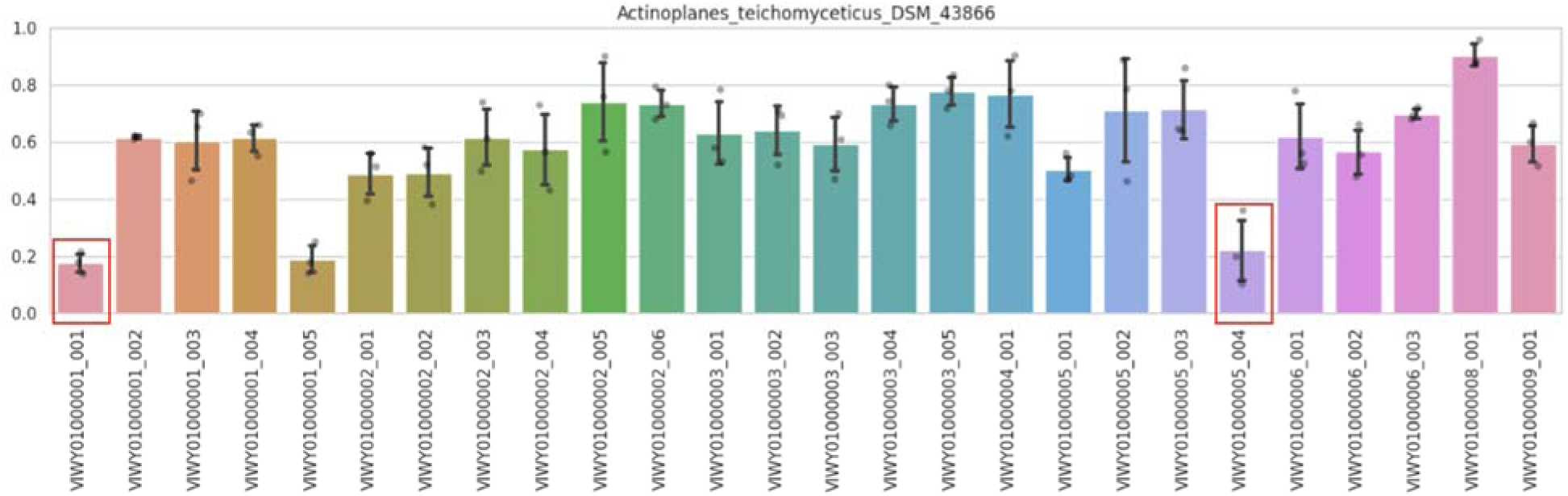
Machine learning–based prediction of antibiotic activity for each biosynthetic gene cluster (BGC) in *Actinoplanes teichomyceticus* DSM 43866. A probability score above 50% indicates a higher likelihood of antibiotic activity. BGCs associated with siderophore biosynthesis are highlighted in red and were predicted by the model to exhibit no antibiotic activity. BGCs associated with teicoplanin (region 8.1) and teichomycin (region 4.1) were predicted with very high antibacterial activities (Table S1).

Due to its predicted propensity to produce antibacterial natural products, we prioritized *A. teichomyceticus* DSM 43866 for natural product isolation. Through bioactivity-guided fractionation, we determined that *A. techomyceticus* DSM 43866 produced an active compound present in a fraction lacking teicoplanin. Further investigation by molecular networking with GNPS^19^ and LC-MS/MS revealed that the most abundant ions in the active fraction corresponded to acyl-desferrioxamine-type siderophores chelated with aluminum (Al^3+^) or iron (Fe^3+^). The activity of this fraction was surprising because acyl-desferrioxamines have been previously reported, but have not been described to have antibacterial activity, and were thought to only serve as iron chelators.^20,21^ To confirm acyl-desferrioxamines can exhibit antibacterial activity, we synthesized two members of this family, confirmed the LC-MS/MS matched the natural compounds, and determined these compounds are active in the presence of Al^3+^ but not in the absence of metal or in the presence of Fe^3+^. We propose that the Al³ -chelated siderophore enhances the uptake of this toxic metal ion into bacterial cells, contributing to its antimicrobial activity.^22,23^

## Results and Discussion

### Genome mining reveals biosynthetic potential of *A. teichomyceticus*

We previously developed a machine learning method capable of identifying if a BGC is likely to produce a natural product with antibacterial activity^18^ and successfully applied it towards the identification of the BGC that produces the dipyrimicins.^24^ We have continued to apply our methodology to identify bacterial strains predicted to produce multiple antibacterial natural products.

*Actinoplanes teichomyceticus* DSM 43866 emerged as a promising candidate. AntiSMASH^25^ analysis revealed that its genome encodes 26 BGCs, with the majority exceeding a 50% probability of producing an antibacterial compound, while a few were predicted to produce compounds without antibacterial activity across the three machine learning prediction algorithms (Table S1). Of these few, two BGCs were identified as the siderophore class by antiSMASH, one of which showed 75% KnownClusterBlast similarity to desferrioxamine with very low predicted scores of antibacterial activities (Fig. 1; Table S1). This result aligns with expectations, as siderophores are not typically known for antibiotic activity. Although the antibiotic-producing capacity of *A. teichomyceticus* has been extensively studied for teicoplanin derivatives, no siderophores have been isolated or characterized from *A. teichomyceticus* DSM 43866, despite antiSMASH 5.1.1 predicting two putative siderophore BGCs.^25^

### Bioactivity-directed isolation of bioactive compounds from *A. teichomyceticus*

Due to the ML-predicted antibiotic-producing capacity of *A. teichomyceticus* DSM 43866 beyond the previously described teicoplanin and teichomycin A1,^12,16,26^ we decided to culture *A. teichomyceticus* DSM 43866 under diverse media conditions to induce secondary metabolite production and investigate if previous studies missed additional antibiotics due to silent BGCs or due to focus on these previously discovered compounds. Specifically, we cultured *A. teichomyceticus* DSM 43866 in liquid media 65, agar media 65 and media ISP7. Crude extracts were subsequently obtained via liquid–liquid and solid-phase extractions (combined synthetic polymeric adsorbent resins) and screened for antibacterial activity against *Bacillus spizizenii* ATCC 6633 using agar diffusion assays. Extracts derived from cultures grown on M65 agar consistently produced larger zones of inhibition compared with its liquid counterparts (Fig. S4).

Guided by the bioassay results, the active crude extracts were subjected to iterative bioactivity-guided fractionation. Among the fractions displaying reproducible antibacterial activity, one chromatographic region eluting between 9.3 and 10.5 min was particularly notable (Fig. S5). This region consisted of three closely eluting peaks that could not be resolved under the HPLC conditions employed. The most abundant peak within this cluster corresponded to an ion at m/z 711.4244 [M+H], accompanied by two lower-intensity features.

High-resolution Orbitrap LC–MS analysis of this bioactive fraction revealed three dominant ions detected in positive mode at m/z 697.4444, 711.4244, and 740.3776 (Fig. 3, Table S2). These ions were consistently observed across replicate injections and accounted for the most abundant ions observed in the fraction. Structural characterization, metal chelation assignments, and comparative database analyses of these features are described in the following section.

**Fig. 3.**
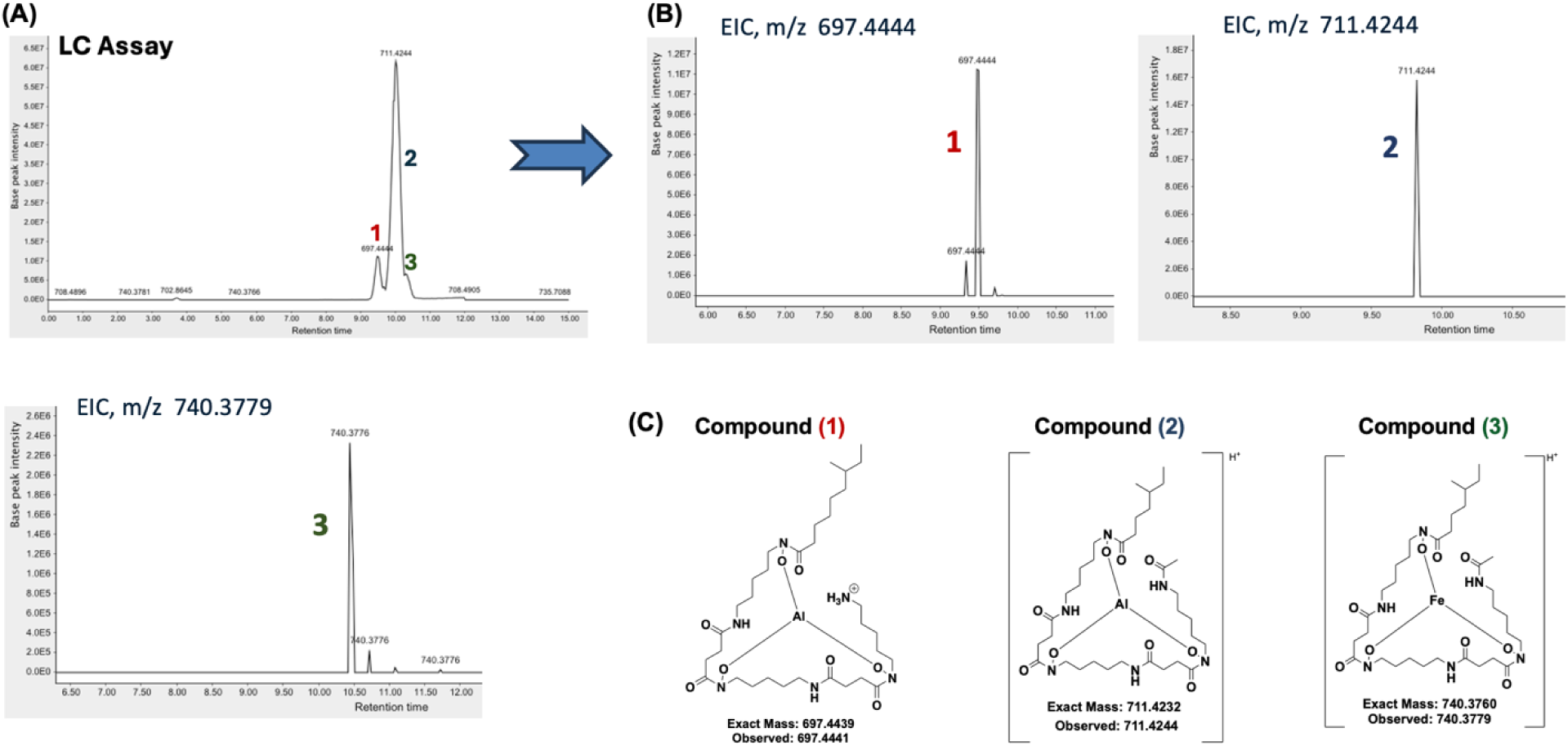
**(A)** Fractionation of the crude extract analyzed by LC assay revealed three co-eluting peaks within the same retention time window. **(B)** Corresponding extracted ion chromatograms (EIC) confirmed three distinct m/z features at 697.4444, 711.4244, and 740.3779. The fraction containing these co-eluting compounds produced a distinct inhibition zone against *Bacillus spizizenii* ATCC 6633 cultured on ISP2/M65 agar. **(C)** High-resolution mass analysis and structural elucidation identified the three closely associated peaks as hydroxamate-type siderophores, and their proposed chemical structures.

### Structure elucidation of naturally isolated siderophores using HRMS and Molecular Networking

To investigate the putative identities of these ions, MS/MS data were analyzed using molecular networking through the Global Natural Products Social Molecular Networking (GNPS) platform.^19^ The most abundant feature at m/z 711.4244 clustered within a molecular family enriched in known ferrioxamine and desferrioxamine siderophores, including nodes with matches to C7 acyl desferrioxamine B + Al (m/z 655.398), desferrioxamine B + Al (m/z 669.414), ferrioxamine D1 + Al (m/z 726.363), and related analogs at m/z 754.407, 768.409, and 780.409 (Fig. 4 and 5). This clustering strongly suggested that the detected ions belong to the ferrioxamine class of siderophores.

**Fig. 4.**
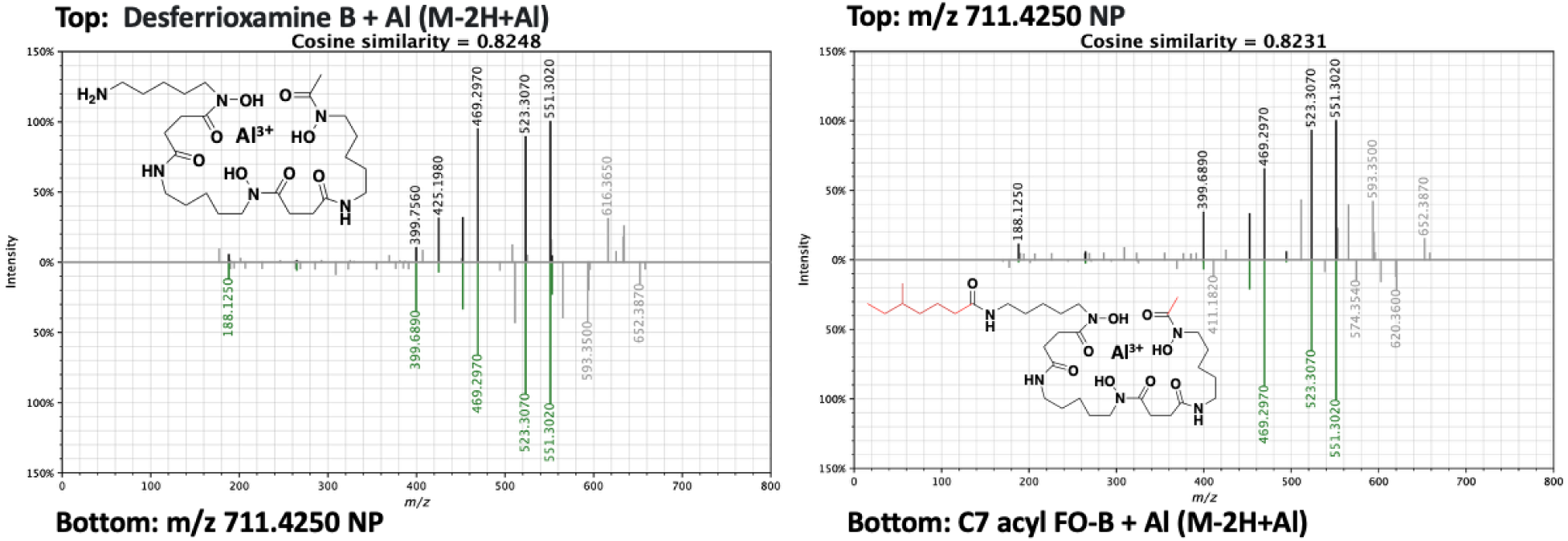
Mirror plot of the experimental and reference MS/MS spectra generated using the Metabolomics Spectrum Resolver. GNPS molecular networking showed that the isolated natural product (m/z 711.4244) clustered with deferoxamine B + Al (M-2H+Al) and C7 acyl FO-B + Al (M-2H+Al), exhibiting high cosine similarity scores of 0.8248 and 0.8231, respectively. Differences in instrument or instrument settings could explain the small deviations in MS/MS spectra compared to the GNPS database.

**Fig. 5.**
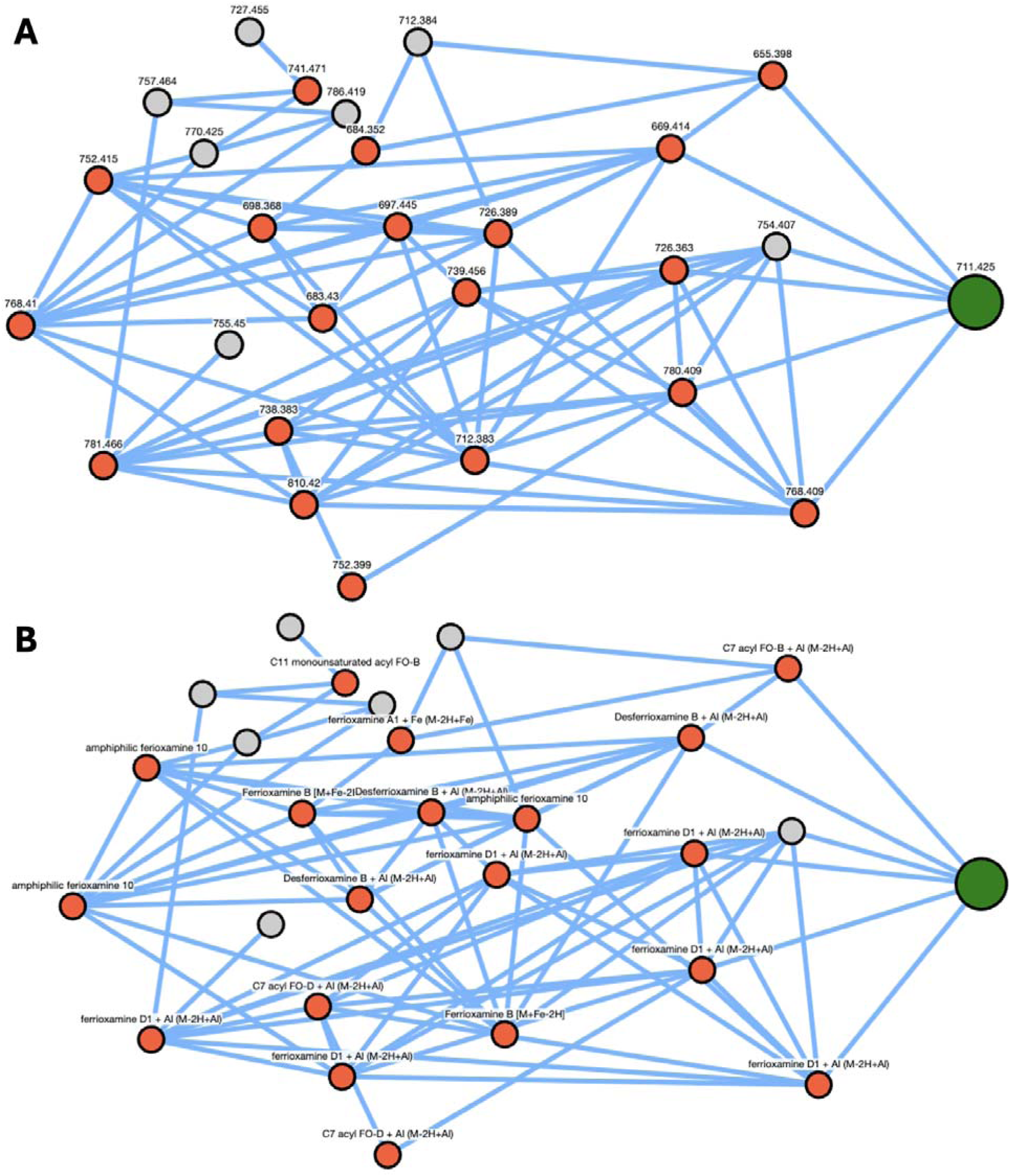
Molecular networking analysis on the Global Natural Products Social (GNPS)^32^ platform for dereplication and visualization of chemical space. The most abundant unknown ion at m/ 711.4244 (highlighted in green circle) clusters within a molecular family containing nodes with fragmentation patterns that match known ferrioxamines, including m/z 655.398 (C7-acyl FO-B + Al), 669.414 (desferrioxamine B + Al), 726.363 (ferrioxamine D1 + Al), 754.407, 768.409, and 780.409. The close spatial clustering of m/z 711.4244 with these known siderophores provides strong evidence supporting its identification as a novel acyl-ferrioxamine-related metabolite. A) Shows a network labeled by m/z value, while B) is labeled by the names of matches in the GNPS database, nodes with no label had no match.

Based on molecular networking relationships, accurate mass measurements, and comparison to reported siderophore analogs,^20^ the ions at m/z 697.4444 and 711.4244 were assigned as Al³ - chelated C9 acyl ferrioxamine B and C7 acyl ferrioxamine D, respectively, while the ion at m/ 740.3776 was annotated as a previously unreported Fe³ -chelated ferrioxamine analog. The experimentally observed masses closely matched the calculated exact masses for the corresponding molecular formulas (C_33_H_61_AlN_6_O_8_, C_33_H_59_AlN_6_O_9_, and C_33_H_59_FeN_6_O_9_), supporting these assignments.

Further confidence in these annotations was provided by MS^2^ spectral comparisons. The m/ 711.4244 feature exhibited high cosine similarity scores (>0.8) to reference spectra for desferrioxamine B + Al and C7 acyl ferrioxamine B + Al in the GNPS database. Mirrored MS/MS spectral comparisons revealed closely aligned fragmentation patterns between the isolated compounds and database references, further substantiating the structural assignments (Fig. 4).

To further characterize the bioactive siderophores produced by *A. teichomyceticus* DSM 43866, in addition to comparing to spectra in the GNPS database, we also analyzed MS/MS fragmentation patterns for m/z of 697.4444, 711.4244, 740.3776 and were able to assign many of the observed peaks to expected fragments of the assigned structures, further supporting the identity of these compounds (Fig. 6–8). This mass spectrum analysis does not enable us to determine the branching pattern of the C7 and C9 acyl chains. We have chosen to draw them with a methyl group on the fourth carbon of the C7 chain and the sixth carbon of the C9 chain, because these branching patterns have been reported several times for acyl-desferrioxamines,^20,21,27–29^ although alternative regioisomers for the acyl chain are possible and have also been reported.^27,30,31^

**Fig. 6.**
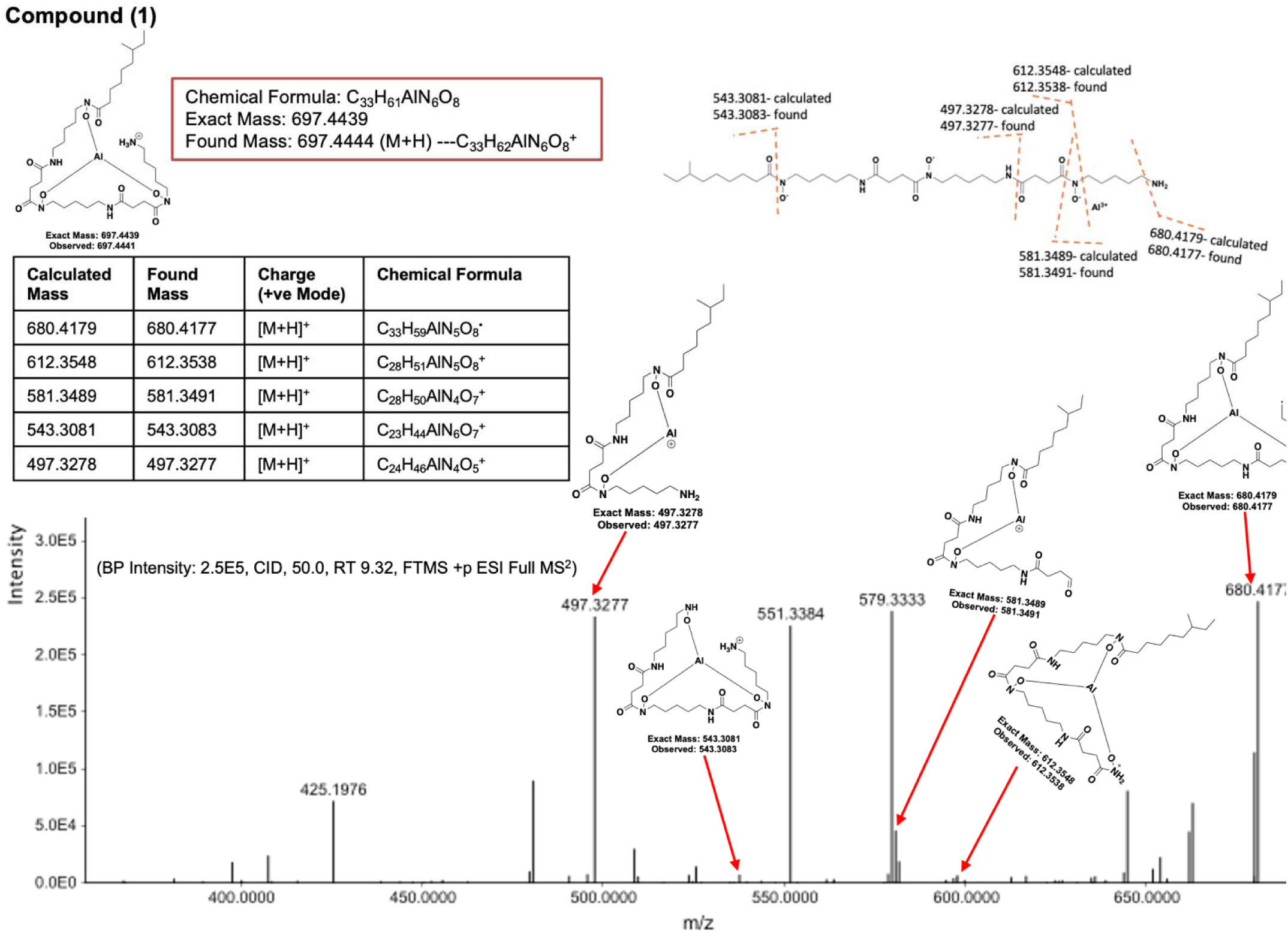
Structural elucidation of Peak 1 (m/z 697.4444 M + H^+^) was performed using high-resolution mass spectrometry (HRMS) on an LC/MS Orbitrap 3 XL (Penn). The LC chromatogram, full MS spectra, and MS² fragmentation patterns were acquired, and th structural interpretation was conducted using MZmine 4.7.^33^ High-resolution Orbitrap LC–MS/MS spectrum acquired in positive-mode ESI showing CID fragmentation of the precursor ion at RT 9.32 min using 50% normalized collision energy, yielding a base peak intensity of 2.5 × 10 in a full MS² (FTMS) scan. (BP Intensity: 2.5E5, CID, 50.0, RT 9.32, FTMS +p ESI Full MS^2^). Likely fragments that match observed masses are shown above the spectra, with arrow between the structure and the associated peak. We also show the full structure of the molecule with bonds that would need to break to make the predicted fragments shown above the spectra.

**Fig. 7.**
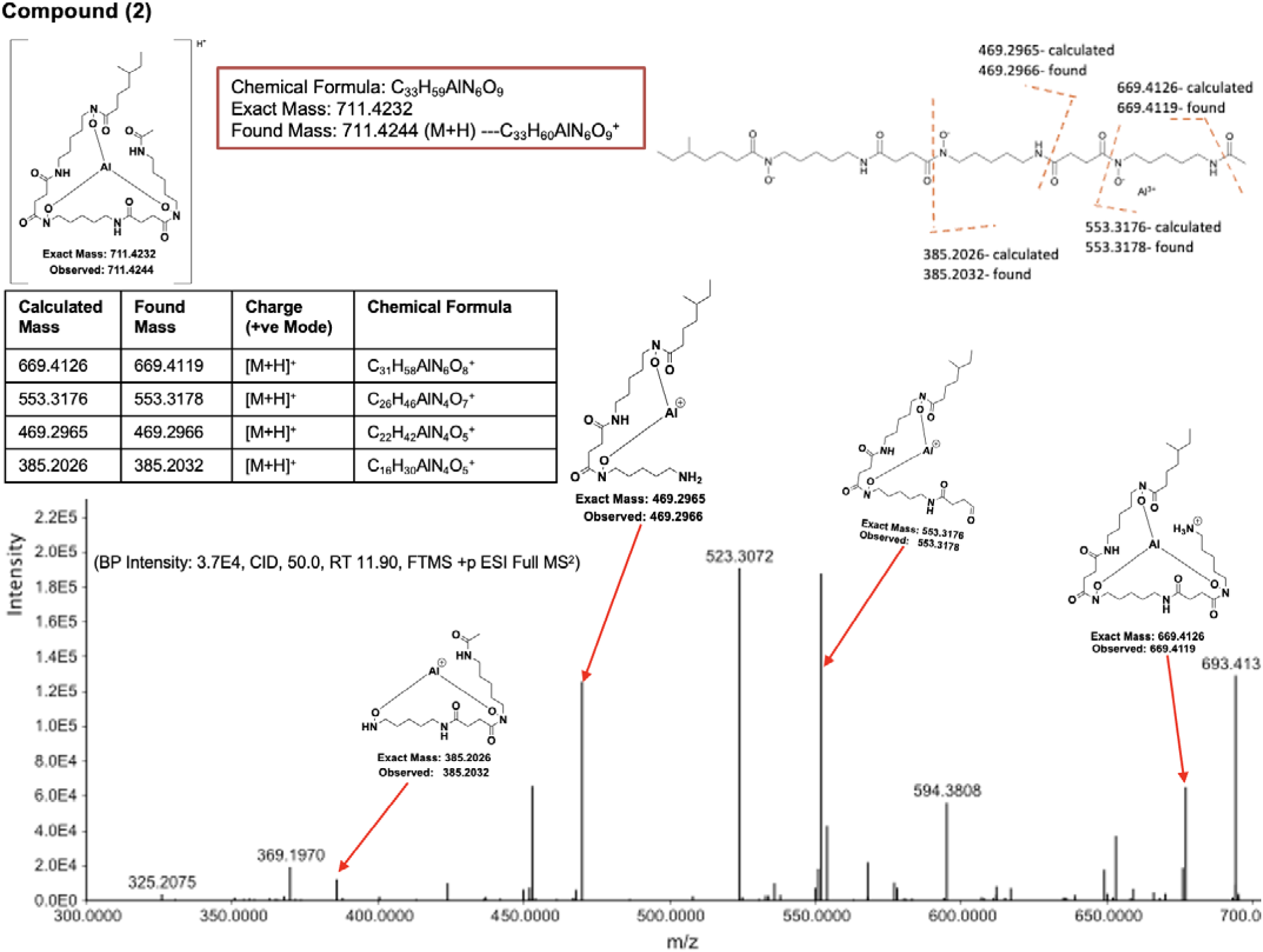
Structural elucidation of the most abundant of the 3 peaks, Peak 2 (m/z 711.4244 [M+H]^+^ was performed using high-resolution mass spectrometry (HRMS) on an LC/MS Orbitrap 3 XL (Penn). The LC chromatogram, full MS spectra, and MS² fragmentation patterns were acquired, and the structural interpretation was conducted using MZmine 4.7. High-resolution Orbitrap LC–MS/MS spectrum acquired in positive-mode ESI showing CID fragmentation of the precursor ion at RT 11.90 min using 50% normalized collision energy, yielding a base peak intensity of 3.7 × 10 in a full MS² (FTMS). (BP Intensity: 3.7E4, CID, 50.0, RT 11.90, FTMS +p ESI Full MS^2^). Likely fragments that match observed masses are shown above the spectra, with arrows between the structure and the associated peak. We also show the full structure of th molecule with bonds that would need to break to make the predicted fragments shown above the spectra.

**Fig. 8.**
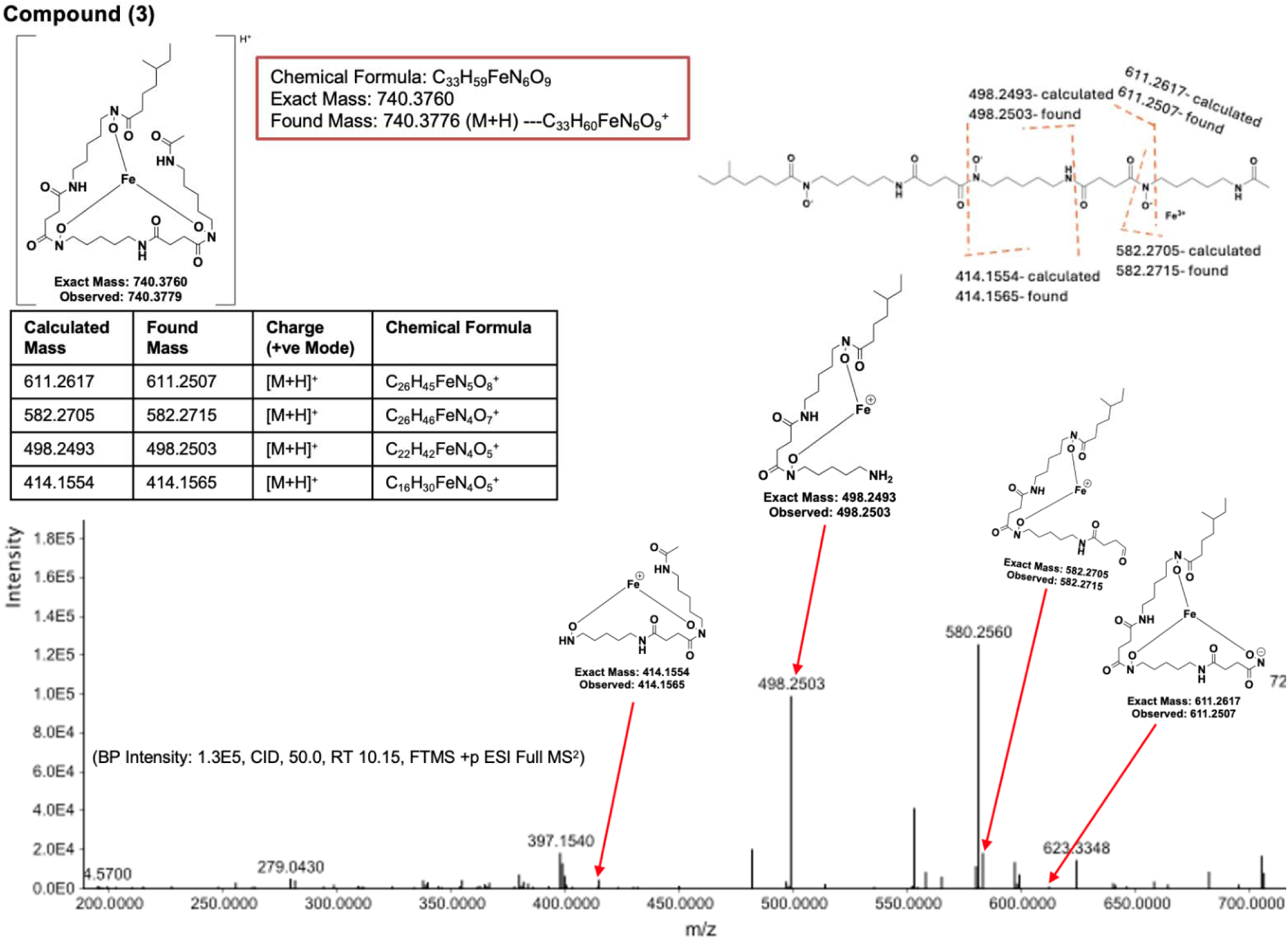
Structural elucidation of Peak 3 m/z 740.3776 [M+H]^+^ was performed using high-resolution mass spectrometry (HRMS) on an LC/MS Orbitrap 3 XL (Penn). The LC chromatogram, full MS spectra, and MS² fragmentation patterns were acquired, and th structural interpretation was conducted using MZmine 4.7. High-resolution Orbitrap LC–MS/MS spectrum acquired in positive-mode ESI showing CID fragmentation of the precursor ion at RT 10.15 min using 50% normalized collision energy, yielding a base peak intensity of 1.3 × 10 in a full MS² (FTMS) scan. (BP Intensity: 1.3E5, CID, 50.0, RT 10.15, FTMS +p ESI Full MS^2^). Likely fragments that match observed masses are shown above the spectra, with arrows between the structure and the associated peak. We also show the full structure of the molecule with bonds that would need to break to make the predicted fragments shown above the spectra.

Taken together, the HRMS/MS fragmentation data and molecular networking analysis provide robust evidence that the active fraction isolated from *A. teichomyceticus* DSM 43866 consists of hydroxamate-type ferrioxamine siderophores. This conclusion is consistent with genome mining predictions.

### CAS Assay Confirmation of Siderophore Production in *Actinoplanes teichomyceticus* DSM 43866

Genome mining of *A. teichomyceticus* DSM 43866 using antiSMASH identified two putative BGCs associated with siderophore biosynthesis, one showing up to 75% similarity to the desferrioxamine E cluster (Fig. 1). This is consistent with our bioactivity-directed isolation of hydroxamate-type siderophores with antibacterial activity from the same strain. To empirically confirm siderophore biosynthesis by *A. teichomyceticus* DSM 43866, we employed the Chrome azurol S (CAS) blue agar assay, a universal method for siderophore detection.^34^ In this assay, *A. teichomyceticus* DSM 43866 was cultured on half M65 agar positioned adjacent to CAS agar within the same Petri dish. After four days, bacterial growth was observed on the M65 agar, and a distinct color change from blue to orange occurred in the neighboring CAS agar, indicating chelation and removal of Fe^3+^ from the CAS dye complex. By 10 days, all of the blue dye in th CAS medium adjacent to the bacterial colony was stripped/sequestered, while control plates lacking bacterial growth retained the original blue color (Fig. 9).

**Fig. 9.**
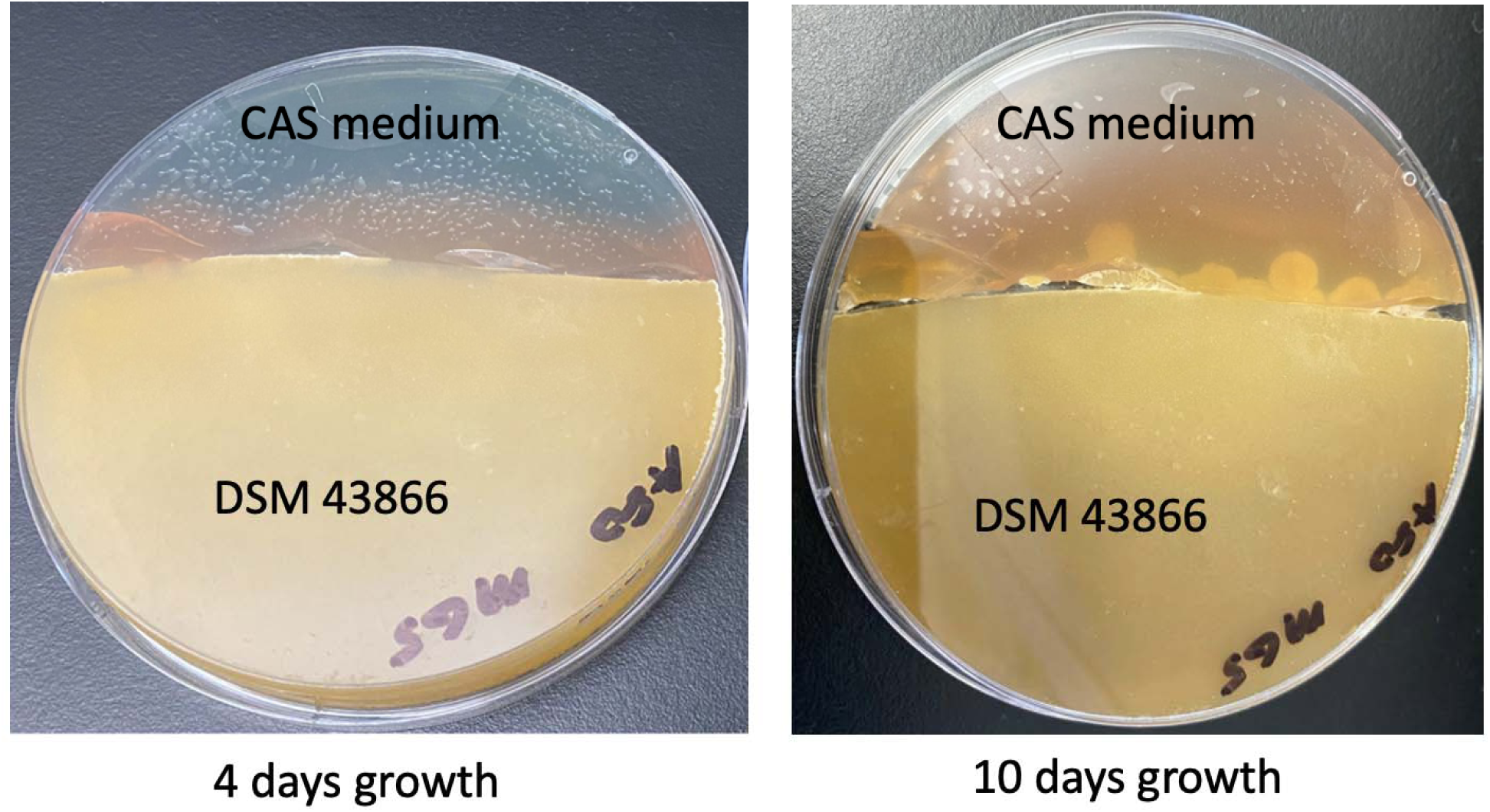
Chrome Azurol S (CAS) agar assay for siderophore detection in *A. teichomyceticus* DSM 43866. The CAS agar plate was incubated alongside an M65 agar plate inoculated with *A. teichomyceticus* DSM 43866. After four days, the CAS dye exhibited a visible color shift from blue to orange, indicating initial siderophore production. By day 10, the medium had completely changed to orange, confirming that *A. teichomyceticus* DSM 43866 actively secretes siderophores capable of chelating and removing iron from the CAS complex.

This characteristic shift in color arises from the equilibrium reaction:

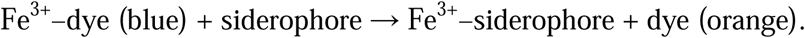

These observations confirm that *A. teichomyceticus* DSM 43866 actively produces siderophores, consistent with genome mining predictions and subsequent metabolite isolation.

### Synthesis of acyl-desferrioxamine D

Isolation of sufficient quantities of the individual siderophores for full NMR characterization proved challenging due to their low abundance, close retention times, and hydrophobicity. Therefore, to confirm that acyl-desferrioxamines are responsible for the observed activity, we set out to synthesize C7 and C9 acyl-desferrioxamine D. C7 acyl-desferrioxamine D was chosen as the synthetic target because it is the ion with the strongest intensity in the active fraction. The m/z corresponding to C9 acyl-desferrioxamine D was at significantly lower intensity, but still observable (Fig. 10). Despite its low intensity we decided to synthesize it because previously isolated acyl-desferrioxamines vary significantly in the identity of the acyl chain,^20,21^ and synthesis of this analog enables preliminary Structure Activity Relationship (SAR) at the acyl chain. Additionally, the synthesis of the C9 acyl-desferrioxamine D analog demonstrates the flexibility of our synthetic method in appending different acyl chains to desferrioxamine D, which may be useful for future studies.

**Fig. 10.**
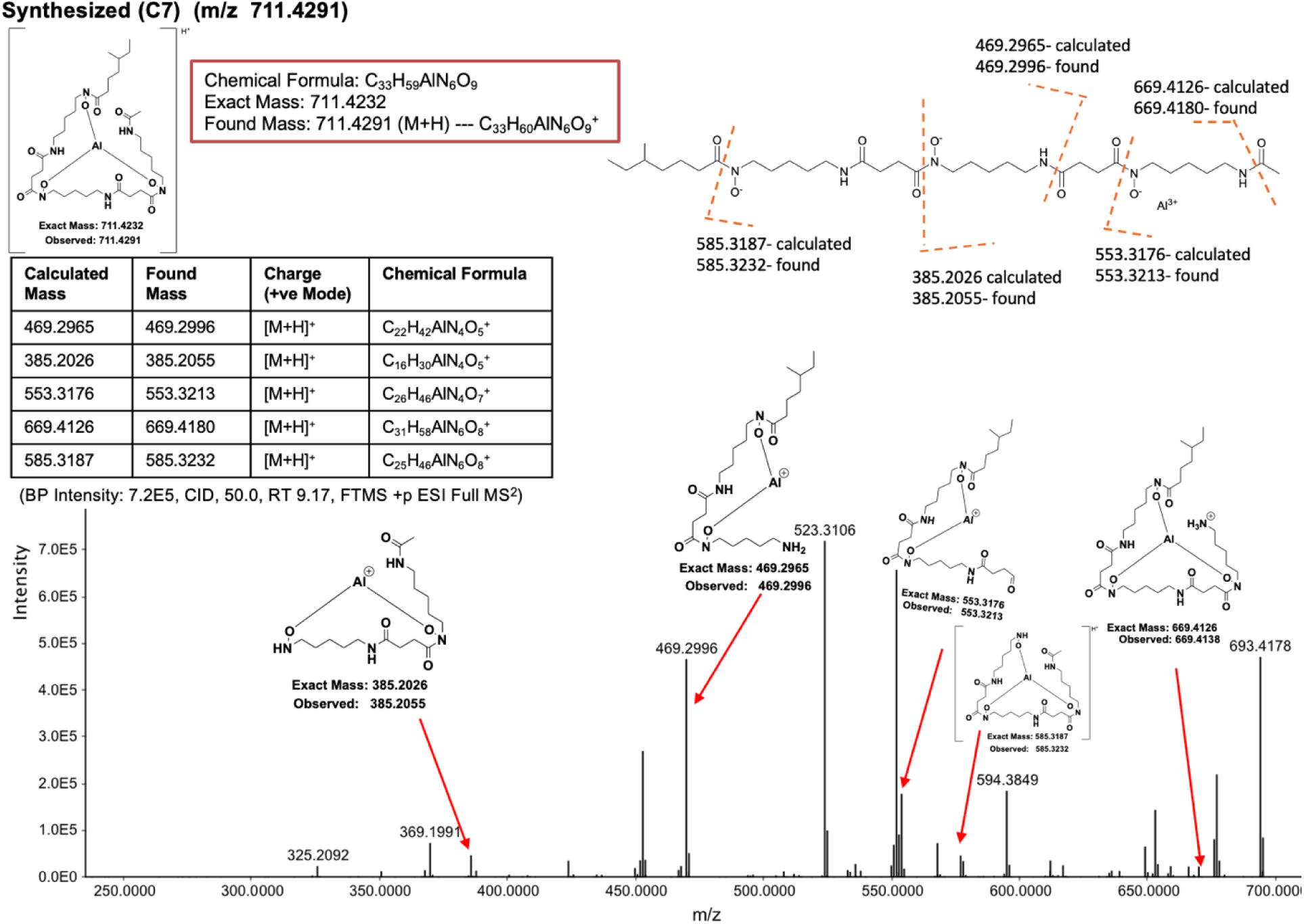
High-resolution LC–MS/MS characterization of the synthetic C7 siderophore (m/z 711.4291, [M+H]□) acquired on an LC–MS Orbitrap 3 XL (Penn). The figure includes the LC chromatogram, full-scan HRMS, and MS² product-ion spectra. Feature detection, alignment, and spectral interpretation were performed in MZmine 4.7. High-resolution Orbitrap LC–MS/MS spectrum acquired in positive-mode ESI showing CID fragmentation of the precursor ion at RT 9.17 min using 50% normalized collision energy, yielding a base peak intensity of 7.2 × 10^5^ in a full MS² (FTMS). (BP Intensity: 7.2E5, CID, 50.0, RT 9.17, FTMS +p ESI Full MS^2^)

The target siderophore analogs were prepared through a modular, solution-phase synthetic route involving amide bond formation, azide reductions, acylation, and global hydrogenolysis (Schemes 1–2). All reactions were conducted under standard inert conditions using commercially available reagents, and intermediates were purified by automated flash chromatography or preparative reverse-phase HPLC. Full experimental details and spectral characterization are provided in the Methods and Supporting Information.

Briefly, tert-butyl (5-azidopentyl)(benzyloxy)carbamate was first reduced via a Staudinger reaction with triphenylphosphine to afford the corresponding amine in 95% yield^35^ (Scheme 1). This intermediate was coupled to the carboxylic acid fragment (compound 7) using EDC/DMAP-mediated amide bond formation to give the protected linear scaffold in 79% yield. Subsequent azide reduction afforded the corresponding amine (36% yield), which was further functionalized through acylation reactions to introduce alkyl side chains (up to 93% yield, depending on the derivative).

Final deprotection steps were achieved using either trifluoroacetic acid for Boc removal (quantitative yield) or catalytic hydrogenolysis (Pd/C, H_2_) to remove benzyl protecting groups, furnishing the final hydroxamate-containing siderophore analogs VU0980806 (2) and VU0980805 in 60–61% yield. All final compounds were confirmed to be ≥95% pure by LC–MS and were fully characterized by ^1^H and ^13^C NMR spectroscopy, with data consistent with the proposed structures.^35–37^

**Scheme 1.**
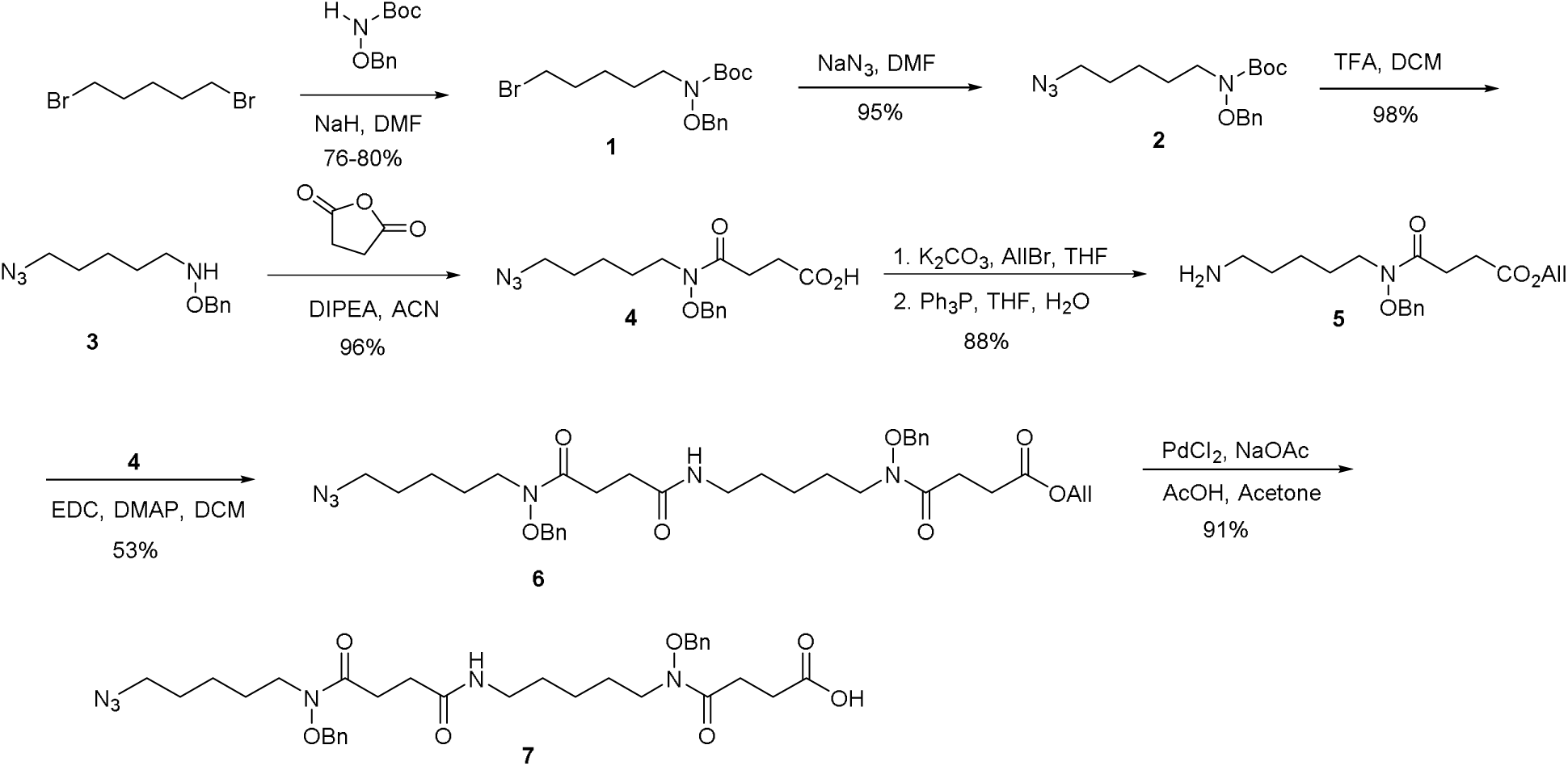
Synthesis of 3-(5-azidopentyl)-14-(benzyloxy)-4,7,15-trioxo-1-phenyl-2-oxa-3,8,14-triazaoctadecan-18-oic acid (**7**).

**Scheme 2.**
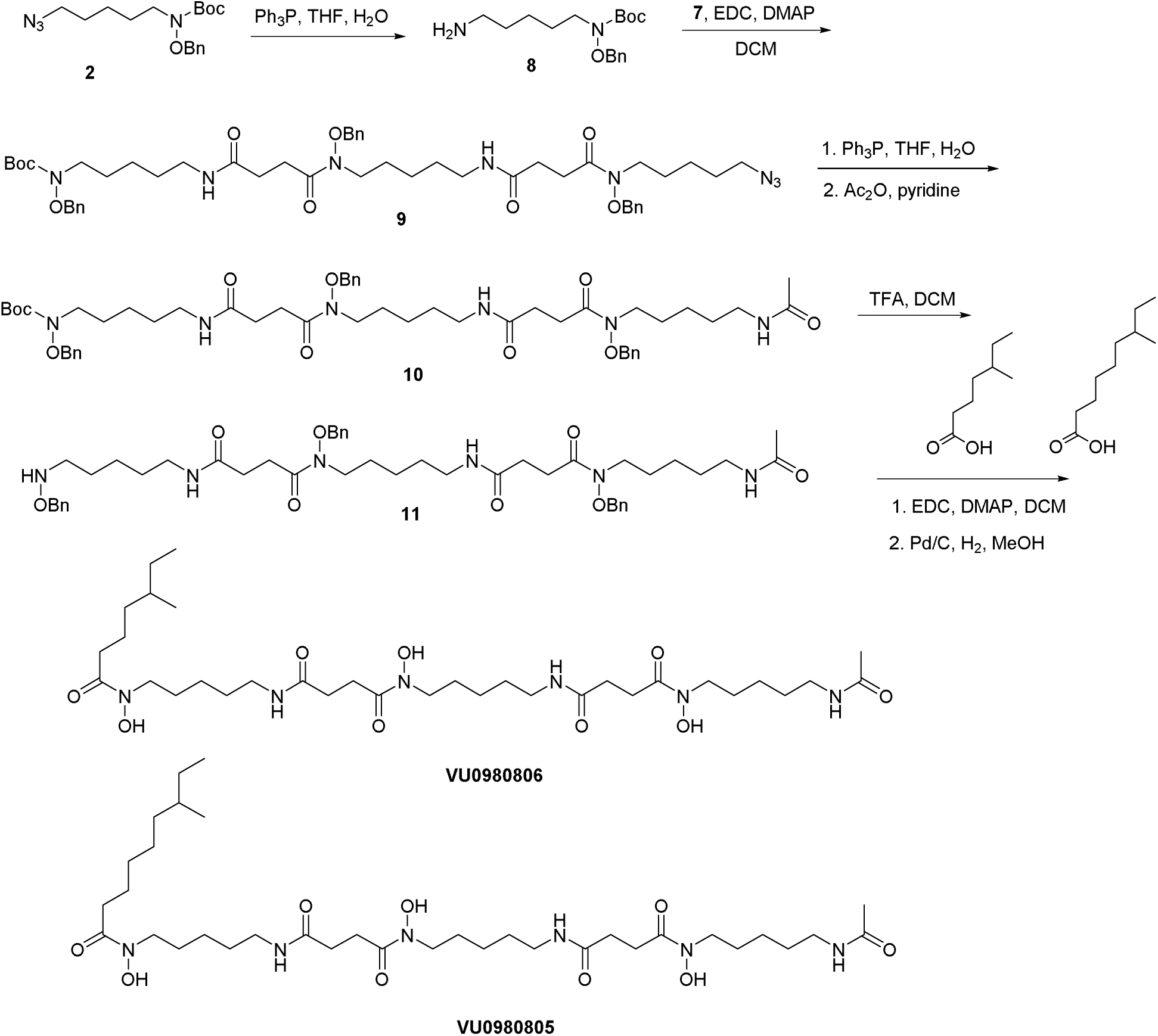
Synthesis of **VU0980806** and **VU0980805.**

### NMR Elucidation of the Synthetic Siderophores C7 and C9 acyl-desferrioxamine D and LC-MS/MS Comparison to Natural Isolates

Synthesis of the target compounds yielded sufficient material for comprehensive NMR characterization of siderophores C7 and C9 (VU0980806 and VU0980805, respectively) which demonstrate successful synthesis of the target compounds. Complete ^1^H and ^13^C NMR datasets are presented in (SI) and summarized in the methods and materials section. Structural identity was evaluated by side-by-side LC–MS/MS comparison of the natural siderophore and the synthetic standard. The synthetic species displayed the expected fragmentation pattern for the precursor ion at m/z 711.4291 (C_33_H_59_AlN_6_O_9_, [M+H]^+^), diagnostic product ions were observed at m/z 469.2996 (C_22_H_42_AlN O ^+^), 385.2055 (C H AlN O ^+^), and 553.3213 (C H AlN O ^+^) (Fig. 10). In addition, the synthetic C7 analog co-eluted within a similar retention time with the natural product and exhibited an indistinguishable fragmentation pattern (Fig. 11). Taken together, the concordant retention time and MS/MS features support structural assignment of to the Al-complexed C7 acyl FO-D isolated from *A. teichomyceticus* DSM 43866 (m/z 711.4244), although it is possible the natural compound has different branching on the acyl chain as this would not likely significantly change retention time or in the MS/MS spectrum.

**Fig. 11.**
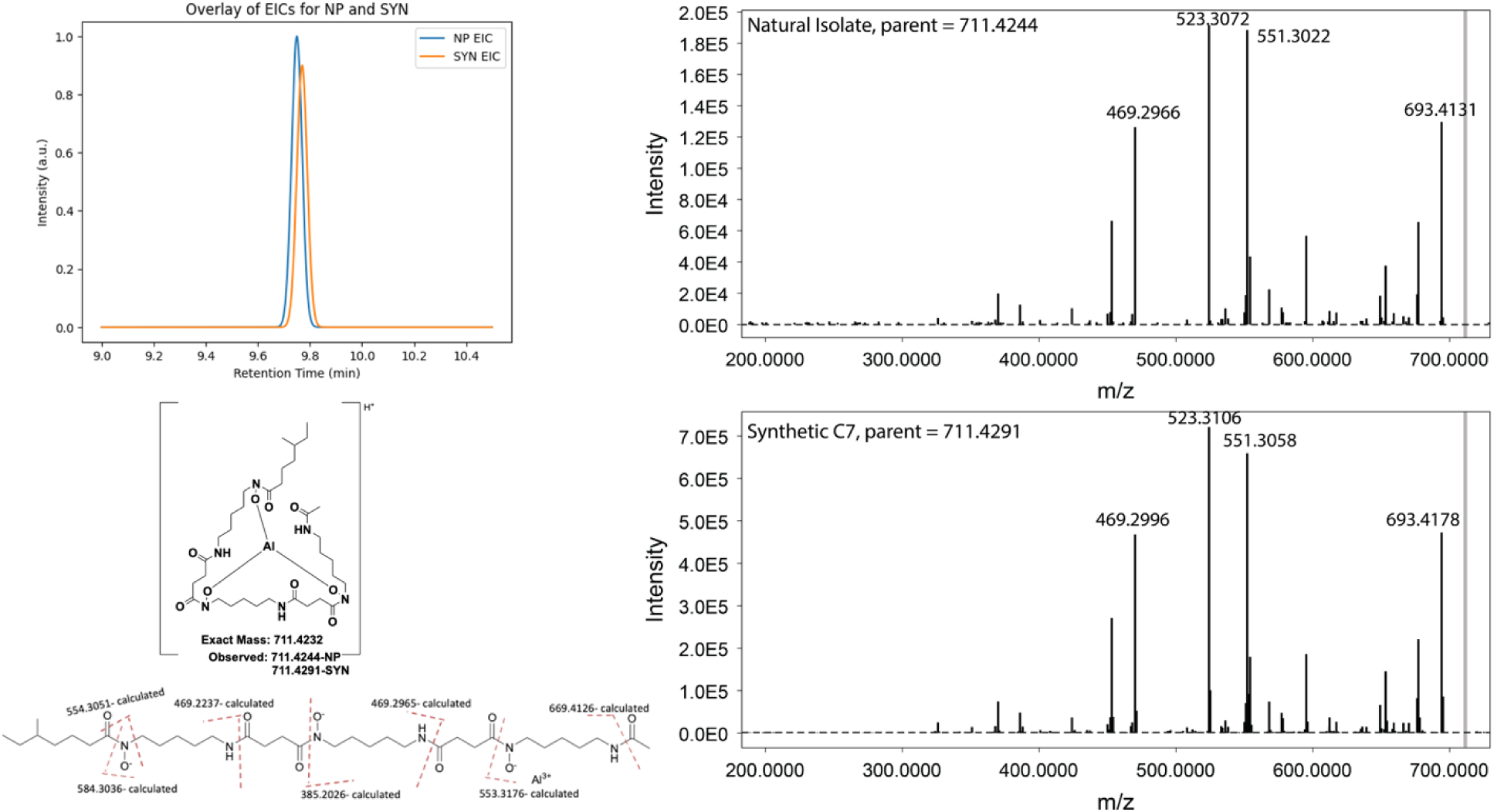
Comparative LC–MS/MS Analysis of Natural Product (NP) and Synthetic (SYN) Counterparts Demonstrating Closely Matched Retention Time and Fragmentation Patterns

### Antibacterial Minimum Inhibitor Concentration (MIC) determination

Prior to MIC determination, antimicrobial susceptibility was evaluated using agar diffusion assays. Crude extracts from *A. teichomyceticus* DSM 43866 demonstrated strong antibacterial activity against *Bacillus spizizenii* ATCC 6633. At the same time, the fractionated hydroxamate siderophore fractions 6 and 7 retained strong activity, producing zones of growth inhibition at doses of 2–8 µl (Fig. S7). In contrast, no significant inhibition was observed against the Gram-negative strain *Escherichia coli* MG1655 (also known as ATCC 700926).

We evaluated the antibacterial activities of the synthetic siderophores C7, C9, and Des (commercially available desferrioxamine mesylate salt) in comparison with their aluminum-chelated counterparts (C7–Al, C9–Al, and Def-Al) using an agar-diffusion inhibition assay against *Bacillus subtilis* (*Bacillus spizizenii*) ATCC 6633. The aluminum-chelated derivatives exhibited noticeably enhanced inhibitory activity relative to their unchelated forms in this assay (Table 1; Fig. S8, S9)

**Table 1.** Minimum Inhibitory Concentrations (MIC, µg/mL) of the isolated siderophore, its synthetic analog, and commercially available desferrioxamine mesylate salt chelated to aluminum ion against *Bacillus subtilis (Bacillus spizizenii)* ATCC 6633 and *Escherichia coli* ATCC 25922.

| Bacterial Strain | MIC ($\mu\text{g/mL}$) – Isolated | MIC ($\mu\text{g/mL}$) – Synthetic Siderophore (C7-Al) | MIC ($\mu\text{g/mL}$) – Synthetic Siderophore (C9-Al) | MIC ($\mu\text{g/mL}$) – desferrioxamine |
| --- | --- | --- | --- | --- |

|  | *Siderophore fraction (2)* |  |  | *mesylate salt (Def-Al)* |
| --- | --- | --- | --- | --- |
| *Bacillus subtilis* (*Bacillus spizizenii*) ATCC 6633 | 16 | 64 | 64 | 64 |
| *Escherichia coli* ATCC 25922 | > 256 | ND | ND | ND |
| *Staphylococcus aureus* ATCC 25923 | ND* | 64 | 64 | 128 |

\*ND exhibited partial inhibition at 16 µg/mL, ND: not determined

Subsequent minimum inhibitory concentration (MIC) assays demonstrated that the siderophore-enriched fraction exhibited selective antibacterial activity against the Gram-positive strain *Bacillus spizizenii* ATCC 6633, with an MIC value of 16 µg/mL. Partial growth inhibition was also observed for *Staphylococcus aureus* ATCC 25923 at approximately 16 µg/mL; however, a definitive MIC could not be established due to limited material availability. No measurable antibacterial activity was detected against the Gram-negative strain *Escherichia coli* MG1655. In contrast, the synthetic analog corresponding to the most abundant aluminum-chelated siderophore (compound 2) displayed substantially weaker activity as did its C9 analog, even in the presence of excess Al^3+^, both with MIC values of approximately 64 µg/mL against both *B. spizizenii* and *S. aureus* (Table 1; Fig 12, Fig. S9). Therefore, a difference of two carbons in the acyl chain does not result in a detectable difference in activity, indicating that a specific acyl chain length is not needed for activity. For the synthetic compounds, inhibition due to Al^3+^ alone was ruled out through controls performed at the same concentration of AlCl_3_ (2.5 mg/mL) that was used to prepare the 64 μg/mL Al-chelated samples that showed inhibition (Fig. S9, S10), as partial growth was observed at this concentration of AlCl_3_. Collectively, these results indicate that the naturally produced siderophores from *Actinoplanes teichomyceticus* exhibit moderate yet selective antibacterial activity toward Gram-positive bacteria. They also indicate that the activity of the natural isolate is not fully explained by the dominant C7 acyl-desferrioxame D.

**Fig. 12.**
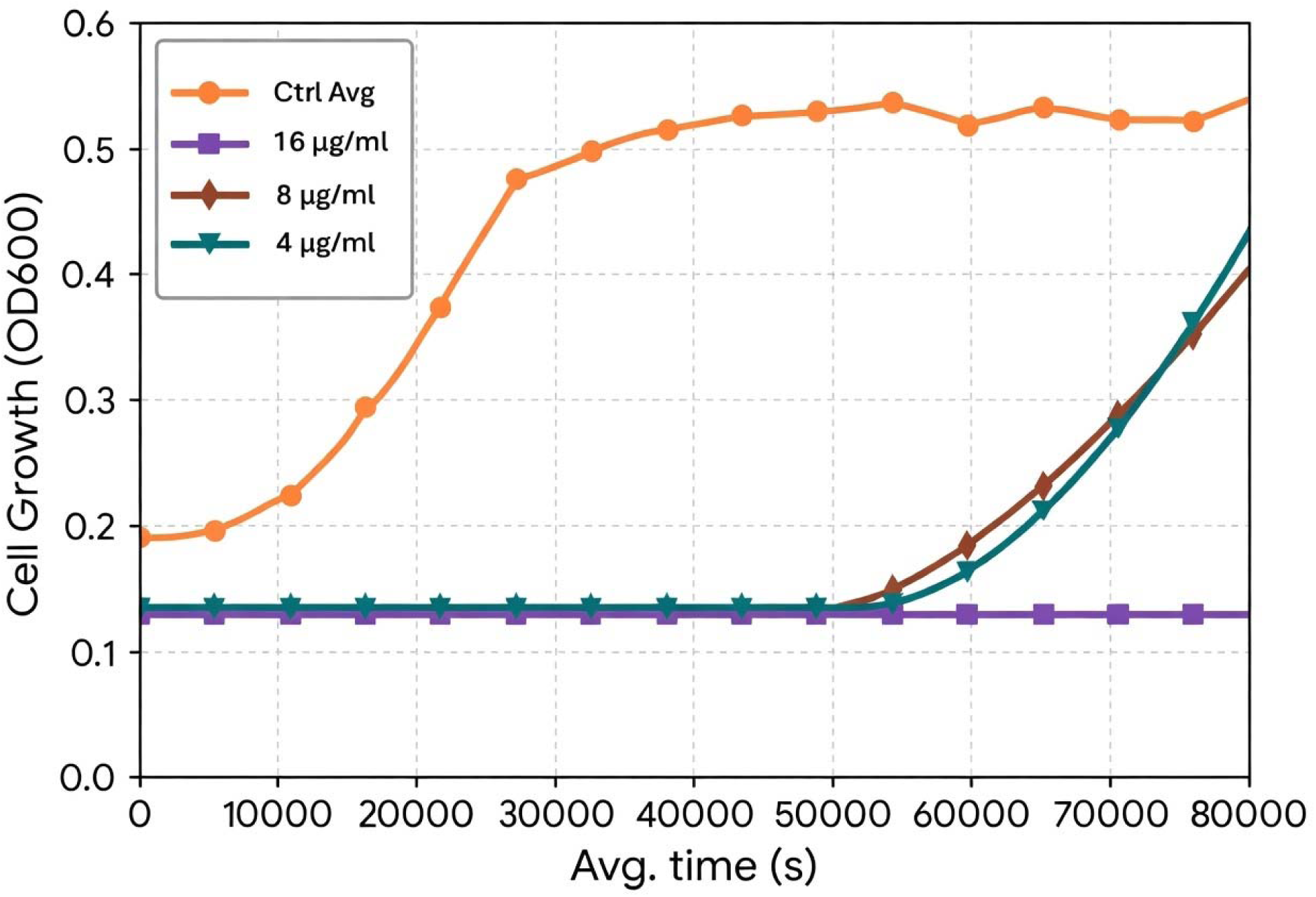
Minimum Inhibitory Concentration (MIC) growth curve of *Bacillus subtilis (Bacillus spizizenii)* ATCC 6633 showing optical density (OD_600_) over time measured using a plate reader. The growth curves represent the control (yellow), which contains only the MHB medium and the bacterial cell, 16 µg/mL (purple), 8 µg/mL (brown), and 4 µg/mL (green) isolated siderophore concentrations. The reduction in OD_600_ values with increasing concentration indicates dose-dependent inhibition of bacterial growth.

**Fig. 13.**
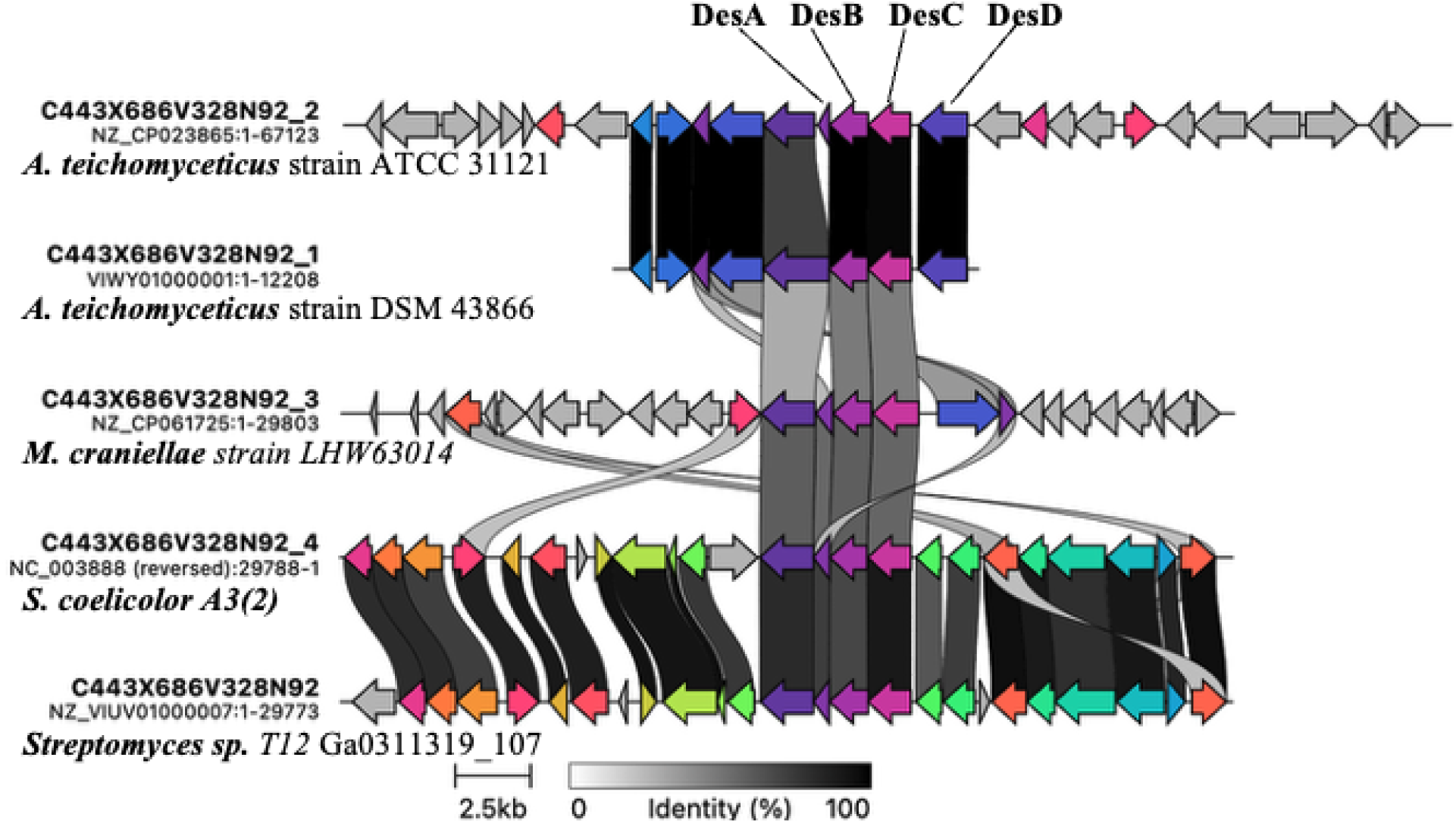
Clinker alignment (CAGECAT) of the putative desferrioxamine-like NI-siderophore locus from *Actinoplanes teichomyceticus* ATCC 31121 (NZ_CP023865) and *A. teichomyceticu* DSM 43866 (VIWY01000001). The conserved core region includes DesA, DesB, DesC, and DesD, and shows homology and synteny with NI-siderophore loci in other actinomycete genomes included for comparison, including *Streptomyces coelicolor* A3(2) (NC_003888), *Micromonospora craniellae* LHW63014 (NZ_CP061725), and *Streptomyces* sp. T12 Ga0311319_107 (NZ_VIUV01000007).

We further evaluated the minimum inhibitory concentration (MIC) of the isolated siderophore fraction under iron- and aluminum-enriched conditions (Figure S12). The fraction was initially tested at 8 µg/mL and subsequently subjected to serial twofold dilutions to a final concentration of 1 µg/mL against *Bacillus subtilis* (*Bacillus spizizenii)* ATCC 6633 over a 24 h incubation period. To assess the influence of metal availability on antibacterial activity, assay media were supplemented with either FeCl_3_ or AlCl_3_ to a final concentration of 200 µM. Across all treatment conditions, only modest differences in antibacterial activity were observed. Notably, at 8 µg/mL, the siderophore fraction consistently maintained complete growth inhibition of *B. spizizenii* ATCC 6633 under both Fe^3+^- and Al^3+^-enriched conditions (Fig S10). These results argue against iron sequestration as the primary mechanism of antibacterial activity. If the dominant species detected by LC-MS, Al^3+^-bound C7 acyl-desferrioxamine D, were present entirely at 8 µg/mL, this would correspond to an estimated molar concentration of approximately 11 µM. Under these conditions, free iron would remain in substantial excess relative to the siderophore, given the presence of 200 µM FeCl_3_ in the assay medium. Consequently, iron deprivation alone cannot account for the observed growth inhibition.

Instead, the data are consistent with a mechanism in which metal-loaded siderophore complexes are actively transported into the bacterial cell. In particular, uptake of the Al^3+^-siderophore complex may lead to intracellular aluminum accumulation, resulting in metal-mediated toxicity and growth inhibition. Such a mechanism aligns with established models of siderophore-mediated metal trafficking and highlights a potential role for non-iron metal transport in the antibacterial activity of this siderophore fraction.

Traditionally, siderophores, including desferrioxamine and acyl-desferrioxamines, have not been associated with direct antimicrobial activity. Instead, they are well known for their role in iron acquisition and have been clinically exploited as delivery vehicles for antibiotics in “Trojan horse” strategies, exemplified by cefiderocol (Fetroja), a siderophore–cephalosporin conjugate.^9^ Siderophores conjugated to toxic platinum(IV)-containing drugs with antibacterial activity have been reported, but these are also synthetic constructs, not naturally-occurring and the platinum is not coordinated by the siderophore itself.^38^ Our findings therefore challenge the conventional understanding of siderophores in nature and suggest additional bioactive functions.

### Homologous Siderophore-Associated Genes Identified Across Actinomycete Genomes by CAGECAT Analysis

We next used CAGECAT (clinker)^39^ to compare the putative desferrioxamine-associated biosynthetic region in *A. teichomyceticus* strains against characterized NI-siderophore BGCs, with the goal of determining whether these *Actinoplanes* loci contain the core genes required for desferrioxamine-family biosynthesis and whether their synteny supports production of related hydroxamate siderophores. In the well-studied actinomycete *Streptomyces coelicolor* A3(2),^20^ desferrioxamine biosynthesis is encoded by the *des* gene cluster (including desABCD),^40^ and curated reference BGCs in MIBiG link defined loci to named siderophore products, including desferrioxamine B/desferrioxamine E (e.g., BGC0000940) and coelichelin (e.g., BGC0000325). *A. teichomyceticus* DSM 43866 and ATCC 31121^12^ contain BGCs consistent with a desferrioxamine-like pathway, with an alignment revealing a highly conserved central block that includes DesA, DesB, DesC, and DesD, with strong homology to corresponding regions in other actinomycete genomes, while flanking regions show greater variability in gene content and organization. To demonstrate that this pathway is widespread and conserved, we also included comparison to other Actinomycetes including an additional non-*Streptomyces* strain, *Micromonospora craniellae* (NZ_CP061725),^31^ and found that the core biosynthetic genes are conserved. Given that DesC is a key determinant of acyl-desferrioxamine diversification,^27^ a logical next step for future work would be to perform a comparative analysis of DesC (e.g., phylogeny and active-site residue conservation) across *Actinoplanes* strains and test whether DesC sequence features correlate with strains reported to produce acylated versus non-acylated desferrioxamines.

### Mechanistic Considerations and Ecological Implications

The unexpected antibacterial activity observed for ferrioxamine-type siderophores is consistent with a “Trojan horse” model in which siderophore–metal complexes promote uptake of disruptive metals and perturb cellular metal homeostasis. This mechanism has been proposed and experimentally supported for desferrioxamine (dfo) scaffolds complexed with Al^3+^ and Ga^3+^, where metal–siderophore complexes inhibited microbial growth more effectively than free metal ions across multiple organisms.^41^ Future work could investigate if Ga^3+^-chelated acyl-desferrioxiames also show antibacterial activity.

Consistent with that precedent, our MIC data show strain-dependent potency. Using the molecular weight of the desferrioxamine mesylate salt used in our experiments (656 g/mol), the observed MIC of 64 µg/mL against *B. spizizenii* corresponds to ∼97 µM. This is a stronger activity than Huayhuaz reported against Gram-positive *Staphylococcus aureus* (IC > 500 µM), suggesting stronger inhibition under our assay conditions.^41^

Ecologically, aluminum is abundant in soils but becomes increasingly mobilized as pH declines; reactive aqueous aluminum species, including Al^3+^, are therefore most available in acidic microenvironments.^42^ Dissolved aluminum in acidic forest soil waters also increases with decreasing pH, supporting the presence of environmentally accessible aluminum pools where siderophore–metal complexation could plausibly occur.^43^ Together, these observations support a model in which ferrioxamine-type siderophores may contribute to microbial competition not only through iron acquisition but also via metal-mediated growth suppression under conditions where soluble aluminum is available.^42,43^

### Machine learning overlooks the potential activity of siderophores

The BGC we identified as the likely acyl-desfferioxamine-producing cluster was assigned a low probability of antibacterial activity of approximately 0.2 (20%) by machine learning classifiers, well below the 0.5 threshold used to indicate likely activity (Fig. 2; Table S1). We interpret this as an incorrect negative prediction that arises from limitations in the training labels rather than a definitive lack of antibacterial relevance. Specifically, the Walker and Clardy model is trained on literature-derived activity annotations and therefore inherits biases created by incomplete testing and reporting.^18^ When a compound class is consistently characterized and curated under a non-antibacterial functional label (for example, “siderophore activity”), antibacterial activity is not captured unless it is explicitly reported and encoded in the source dataset.^18^ In our curated database, desferrioxamine and other hydroxamate siderophores are not annotated as antibacterial, creating a systematic blind spot in which the model has limited or no positive examples to learn antibacterial-associated patterns for this chemical class. Consequently, hydroxamate siderophore BGCs are expected to be assigned low antibacterial probabilities even when their biological effects could plausibly contribute to growth suppression or competitive outcomes under relevant conditions. This illustrates a key limitation of activity-prediction models: when activities have been overlooked or unrecorded for an entire compound class, predictions for that class will likely be inaccurate.

### Conclusion

In this study, we combined genome mining, siderophore detection, bioactivity-guided fractionation, HRMS/MS analysis, molecular networking, and synthetic chemistry to identify ferrioxamine-type siderophores with unexpected antibacterial activity. Our results show that these compounds are active in the presence of Al^3+^ but not in the presence of Fe^3+^ or in the absence of metal, suggesting that the activity is Al^3+^-dependent and might be due to a “Trojan metal” effect, where the siderophore transports the toxic Al^3+^ into the cell. Moreover, they highlight siderophores as an underexplored class of bioactive metabolites with potential therapeutic significance, particularly against Gram-positive pathogens. The increased activity of the natural acyl-desferrioxamine fraction compared to the synthetic C7- and C9-acyldesferrioxamines is an area of particular interest for future investigation. It is possible that there is a less abundant compound that is even more active than C7- and C9-acyldesferrioxamine, or that there are synergistic pairs of compounds present in the fraction.

## Materials and Methods

### General experimental procedures and materials

The *Actinoplanes teichomyceticus* DSM 43866 strain used in this study was obtained from the German Collection of Microorganisms and Cell Cultures (DSMZ). The *Bacillus spizizenii* ATCC 6633 indicator strain was purchased from Fisher Scientific. Additional indicator strains employed in the bioactivity assays, including *Escherichia coli* MG1655 (ATCC 700926), *Pseudomonas aeruginosa* (ATCC 27853), and *Staphylococcus aureus* (ATCC 25923), were obtained from the American Type Culture Collection (ATCC). BD Difco ISP2 medium and M65 (GYM *Streptomyces* medium) were procured from Fisher Scientific and used for bacterial cultivation. HyperSep C18 cartridges (Fisher Scientific) were employed for solid-phase extraction. Mueller–Hinton broth (MHB; Thermo Scientific) was used in the minimum inhibitory concentration (MIC) assays. Analytical separations were performed using HyperSIL GOLD C18 columns (Thermo Scientific). All solvents used for organic extraction, chromatographic separation, and mass spectrometric analyses were HPLC- and LC-MS-grade and obtained from Fisher Scientific or Sigma Aldrich. Deuterated NMR solvents were purchased from Sigma Aldrich. Aluminum chloride (anhydrous 99%) and ferric chloride (anhydrous) were obtained from Thermo Fisher Scientific

### Genome mining and BGC activity prediction

The genome sequence of *Actinoplanes teichomyceticus* DSM 43866 (NCBI accession: VIWY01000001; whole-genome shotgun sequence VIWY01000000, version VIW01000001.1) was retrieved from the NCBI GenBank database. To maintain consistency with the machine learning framework of Walker and Clardy, which was trained using antiSMASH version 5 and Resistance Gene Identifier (RGI) version 5, the same versions were applied for biosynthetic gene cluster (BGC) annotation and activity prediction. Comparative analysis indicated no major discrepancies between annotations generated by antiSMASH versions 5 and 7; thus, antiSMASH 5 was used for the majority of downstream bioinformatic analyses.^17,44^ (Fig.1, 2, Table S1).

To further explore siderophore biosynthetic potential, BGCs from *Micromonospora craniellae* LHW63014 (GenBank: NZ_CP061725), *Streptomyces sp.* T12 Ga0311319_107 (GenBank: NZ_VIUV01000007), *and Streptomyces coelicolor* A3(2) (GenBank: NC_003888) were aligned with the siderophore-associated BGCs of *A. teichomyceticus*. Alignments and visualizations were performed using clinker through the CAGECAT webserver (https://cagecat.bioinformatics.nl/), enabling the identification of conserved siderophore biosynthetic gene architectures across strains.

### Culture Conditions and Fermentation

*Actinoplanes teichomyceticus* DSM 43866 was obtained from the DSMZ culture collection. Seed cultures were initiated in liquid Medium 65 (M65) and incubated at 28 °C with shaking at 200 rpm for 3–4 days. Production cultures were prepared by inoculating 2% (v/v) of the seed culture into fresh M65 medium and incubating under identical conditions for 7–14 days. Both liquid and agar-based cultivation formats were employed to evaluate media-dependent metabolite expression. Notably, agar-based cultures yielded the production of bioactive siderophores. The composition of M65 (GYM Streptomyces medium) was as follows: glucose, 4.0 g; yeast extract, 4.0 g; malt extract, 10.0 g; CaCO_3_, 2.0 g (omitted for liquid medium); agar, 20.0 g; distilled water, 1000 mL. The pH was adjusted to 7.2 prior to autoclaving and the addition of agar.

### *Actinoplanes teichomyceticus* 16S rRNA Gene Amplification and Sequencing

Genomic DNA from *Actinoplanes teichomyceticus* DSM 43866 was extracted following a thermal lysis protocol. Briefly, 50 µL of bacterial culture was centrifuged at 4200 rpm for 30 min at 4 °C to pellet the cells. The supernatant was discarded, and the pellet was resuspended in 100 µL of PrepMan reagent (Thermo Scientific). Samples were boiled at 100 °C for 15 min in a Thermo Scientific block heater, cooled on ice for 5 min, and centrifuged at maximum speed (12,000 rpm) for 2–5 min to clarify the lysate.

The resulting supernatant served as the DNA template for polymerase chain reaction (PCR) amplification of the 16S rRNA gene using universal bacterial primers Eub27F (5′-AGAGTTTGATCMTGGCTCAG-3′) and Eub1492R (5′-GGTTACCTTGTTACGACTT-3′). Each 50 µL PCR reaction was prepared using 25 µL of Q5 High-Fidelity 2X Master Mix (New England Biolabs), 2.5 µL each of 10 µM forward and reverse primers (final concentration 0.5 µM each), a variable volume of DNA template containing less than 1,000 ng of total DNA, and nuclease-free water to a final volume of 50 µL.

Reactions were assembled on ice, briefly centrifuged, and amplified under the following thermocycling conditions: initial denaturation at 98 °C for 30 s; 30 cycles of denaturation at 98 °C for 10 s, annealing at 55 °C for 20 s, and extension at 72 °C for 30 s; followed by a final extension at 72 °C for 2 min.

PCR amplicons were analyzed by agarose gel electrophoresis in 0.5× TAE buffer using gels stained with SYBR Safe DNA gel stain. Five microliters of each PCR product mixed with loading dye, along with a DNA ladder, were electrophoresed at 100 V for 20–30 min. Bands of approximately 1,500 bp were visualized under blue-light illumination using a Gel-Doc imaging system (Bio-Rad), confirming successful amplification of the 16S rRNA gene.

Amplified products were purified by combining 5 µL of PCR product with 2 µL of ExoSAP-IT (Thermo Fisher Scientific), centrifuged briefly, and then the PCR products were treated with ExoSAP-IT and incubated in a thermocycler at 37 °C for 15 minutes, followed by enzyme inactivation at 80 °C for 15 minutes. The purified amplicons were diluted to a concentration of 40 ng/µL, verified using a NanoDrop spectrophotometer, and submitted for Sanger sequencing through GenHunter. The resulting sequences were analyzed using NCBI BLASTn, confirming the isolate identity as *Actinoplanes teichomyceticus* DSM 43866 (Fig. S1–S3).

### Extraction of Secondary Metabolites

Following 14 days of fermentation, agar cultures were frozen at –20 °C overnight, then thawed at room temperature for >5 h prior to extraction. The agar plates were soaked in equal volumes of ethyl acetate and deionized water (1 L per 20 plates) and subjected to sonication for approximately 1 h. The resulting liquid phase was separated from the agar matrix via vacuum filtration, and the organic (ethyl acetate) layer was collected and concentrated under reduced pressure.

The residual aqueous phase from the ethyl acetate extraction was further extracted by adsorption onto synthetic resin beads. The aqueous extract was transferred to a 2 L flask and amended with Amberlite XAD-4, Amberlite XAD-7HP, and Dianion HP20 resins (Thermo Scientific) to a combined concentration of approximately 10% (w/v). The suspension was incubated at 30 °C with shaking at 150 rpm for 1 h to facilitate the adsorption of secondary metabolites onto the resins. The beads were pelleted by centrifugation (4000 rpm, 30 min), and the supernatant was discarded.

Sequential elutions were performed by agitating the pelleted resins with 800 mL of methanol followed by 800 mL of acetone, each at 30 °C and 150 rpm for 1 h. After each step, the mixtures were centrifuged and vacuum-filtered to remove residual solids. The ethyl acetate, methanol, and acetone extracts were combined and concentrated under reduced pressure using a rotary evaporator. The resulting residue was reconstituted in 25% acetonitrile (ACN), yielding the crude extract for subsequent bioactivity assays and chromatographic purification.

### Bioactivity-guided isolation

The crude extract was subjected to multiple rounds of HPLC purification. Fractions were collected across retention times corresponding to peaks eluting between 2–15 min of the method. Hydroxamate-type siderophores were identified as the bioactive metabolites of interest through bioactivity-guided fractionation. Each fraction was concentrated in vacuo and tested against *Bacillus spizizenii* ATCC 6633. Fractions displaying activity, defined as an inhibition zone greater than that observed for the 25% acetonitrile (ACN) negative control, were selected for further purification.

Bioactivity was consistently observed in fractions eluting at ∼9.3–10.5 min. These fractions were subsequently subjected to additional rounds of HPLC purification on a Thermo Fisher Vanquish HPLC system equipped with a photodiode array detector and automated fraction collector.

Separations were carried out using a Hypersil GOLD C18 column (3 µm, 4.6 × 100 mm) under a linear gradient of 0–100% ACN/H O (95:5) containing 0.1% (v/v) formic acid over 10 min. Three closely eluting peaks within the 9.5–10.5 min window were collected, corresponding to the active siderophore metabolites (Fig S4–S5).

### LC-HRMS Data Acquisition

#### Sample Preparation

Crude extracts were centrifuged at 4,000 rpm for 10 min, and the resulting supernatant was collected and reconstituted in 25% acetonitrile (ACN/H_2_O, v/v) prior to analysis.

### Chromatographic Conditions

Chromatographic separation was performed on a Waters ACQUITY UPLC system (Waters Corporation, Milford, MA, USA) using a Hypersil GOLD C18 reverse-phase column (3 µm, 4.6 × 100 mm). The mobile phase consisted of solvent A [0.5% formic acid in H□O/ACN (95:5, v/v)] and solvent B [0.5% formic acid in ACN/H O (95:5, v/v)]. Elution was carried out at a flow rate of 0.300 mL/min using the following linear gradient: 0–1 min, 100% A; 1–10 min, 0–100% B; 10–12 min, 100% B; 12.1–14 min, 100% A (re-equilibration). The total run time was 15 min. The sample manager was operated in partial loop injection mode using weak wash solvent (water, 500 µL) and strong wash solvent (acetonitrile, 200 µL) between injections.

### High-Resolution Mass Spectrometry

Mass spectrometric detection was performed on an LTQ Orbitrap XL hybrid mass spectrometer (Thermo Fisher Scientific, Bremen, Germany) equipped with an electrospray ionization (ESI) source operated in positive ion mode. Full-scan MS1 spectra were acquired using the Fourier transform mass spectrometry (FTMS) analyzer over a mass range of m/z 150–2000, at a resolving power of 30,000 FWHM, in profile mode with normal mass range setting.

Data-dependent acquisition (DDA) was employed for MS^2^ fragmentation. Precursor ion isolation was performed with an isolation width of 1.0 m/z. Fragmentation was achieved by collision-induced dissociation (CID) with a normalized collision energy (NCE) of 35.0%, an activation Q value of 0.250, and an activation time of 30.0 ms. Dynamic exclusion was enabled to minimize repeated fragmentation of the same precursor ions across the chromatographic run. Source fragmentation was disabled to avoid in-source adduct formation and ensure spectral integrity for unknown compound identification.

Instrument calibration was performed using the standard Thermo calibration solution prior to analysis, achieving a mass accuracy of <5 ppm across the acquisition range. Data acquisition was controlled using Xcalibur software (Thermo Fisher Scientific). Raw data files were processed for feature detection, dereplication, and putative structural annotation of natural product candidates against relevant spectral and chemical databases [MZmine, GNPS, Cystoscope].^32,33^

### Spectroscopic and Analytical Characterization

UV–Vis spectra were recorded at 215 nm using a Thermo Fisher Vanquish HPLC system equipped with a photodiode array detector. Analytical separations were performed on a Hypersil GOLD C18 column (3 μm, 4.6 × 100 mm) at a flow rate of 2.0 mL/min, employing a linear gradient of 0–100% acetonitrile (ACN)/H O (95:5) containing 0.1% (v/v) formic acid over 10 min.

High-resolution electrospray ionization mass spectra (ESI-HRMS) were acquired on an LTQ-Orbitrap 3 XL mass spectrometer (Thermo Scientific) at the Vanderbilt University Mass Spectrometry Core Facility. Data were collected in positive ion mode across the m/z range 100–2000. MS/MS data were collected using data-dependent acquisition with stepped collision energies (30–50 eV). Accurate mass measurements were compared against calculated formulas, and structural assignments were made from fragmentation patterns and processed using Xcalibur software and MZmine 4.0.

NMR experiments, including ¹H, ¹H–¹H COSY, ¹H–¹³C HSQC, and ¹H–¹³C HMBC, were recorded in methanol-d□(CD□OD) for the synthesized siderophore C7 and C9 on 600 and 800 MHz Bruker spectrometers at the Vanderbilt University Small Molecule NMR Facility. Spectra were processed and analyzed using TopSpin (Bruker) and MestReNova (MNOVA, Mestrelab Research) software (Fig. S11–S12).

### Antibacterial Assays and Minimum Inhibitory Concentration (MIC) Determination

Initial antimicrobial activity was assessed using agar diffusion assays (Kirby–Bauer method) on M65 or ISP2 agar plates.^45^ Aliquots of 8–10 µL of crude or fractionated extracts were spotted onto the agar surface, and zones of inhibition were recorded after incubation.

The minimum inhibitory concentration (MIC) was defined as the lowest concentration of hydroxamate-type siderophores that completely inhibited visible growth. MIC values were determined against *Bacillus spizizenii* ATCC 6633, *Escherichia coli* MG1655 (ATCC 700926), and *Staphylococcus aureus* ATCC 25923 following Clinical and Laboratory Standards Institute (CLSI) guidelines with minor modifications.

Naturally isolated hydroxamate-type siderophores were serially diluted sixfold in MHB from 16 to 0.5 μg/mL in a 96-well plate format (50 μL per well). An equal volume (50 μL) of bacterial inoculum prepared according to Kadeřábková et al.^46^ was added to each well, resulting in final assay concentrations of 8–0.25 μg/mL. For the synthetic counterparts, AlCl_3_ (initial concentration 10 mg/mL) was first mixed with each compound, and the resulting mixture was then serially diluted from 128 to 16 μg/mL of siderophore, yielding final concentrations of 64–8 μg/mL of siderophore after inoculum addition. Plates were incubated at 30 °C for 24 h, and bacterial growth was monitored by measuring OD_600_ using a Varioskan LUX Multimode Microplate Reader (Thermo Scientific). Tetracycline was used as a positive control. All assays were performed in triplicate.

### Molecular Networking Analysis

Raw LC–MS/MS data were converted to .mzML format and uploaded to the Global Natural Products Social (GNPS) platform. Spectral data were processed using the molecular networking workflow with a cosine score cutoff of 0.7 and a minimum of 6 matched fragment peaks. Networks were visualized using GNPS embedded Cytoscape and Cytoscape 3.9. Known siderophore nodes were annotated based on GNPS library matches and literature-reported fragmentation patterns (Fig. 5).

### Chrome Azurol S (CAS) Siderophore Detection Assay

Siderophore production by *A. teichomyceticus* DSM 43866 was evaluated using the Chrome Azurol S (CAS) blue agar assay.^34,47^ CAS agar plates were prepared according to the stepwise protocol of Louden et al,^34^ based on Schwyn and Neilands (1987).^3^ The CAS dye complex was prepared by combining CAS, FeCl_3_·6H_2_O in 10 mM HCl, and hexadecyltrimethylammonium bromide (HDTMA) to generate the characteristic blue solution. This was incorporated into a buffered MM9/PIPES agar base (pH 6.8) supplemented with casamino acids and glucose, before aseptic pouring of plates.

For the assay, *A. teichomyceticus* DSM 43866 was inoculated onto half-strength M65 agar positioned adjacent to CAS agar within the same Petri dish. Plates were incubated at 28 °C and observed for color changes over 4–10 days. By day 4, orange halos appeared in the CAS agar adjacent to bacterial colonies, indicating removal of Fe^3+^ from the dye complex by secreted siderophores. By day 10, the CAS agar surrounding the colony was completely decolorized, while uninoculated control plates retained their original blue color. This confirmed genome-predicted siderophore biosynthesis consistent with hydroxamate-type metabolites identified in the bioactivity-guided isolation (Fig. 9, Fig. S6, S7).

### Chemical Synthesis of Novel Siderophore Analog

#### General Experimental Information

All reagents and solvents were obtained from commercial suppliers and used as received unless otherwise noted. Anhydrous solvents were obtained from an MBraun MB-SPS solvent purification system. Reactions were performed under an inert atmosphere of Argon unless otherwise specified.

Analytical thin-layer chromatography (TLC) was carried out on silica gel plates (60F254), and visualization was achieved under UV light or by staining with KMnO_4_. Flash column chromatography was performed using a Teledyne ISCO CombiFlash system.

^1^H NMR spectra were recorded at room temperature on Bruker 400 MHz spectrometers, and ^13^C NMR spectra were obtained at 100-150 MHz. Chemical shifts (δ) are reported in ppm relative to the residual peaks of the deuterated solvent, with multiplicity (s = singlet, d = doublet, t = triplet, q = quartet, m = multiplet, br = broad), coupling constants (Hz), and integration values indicated. LC/MS analyses were performed on an Agilent 1260/G6125B Quadrupole system with electrospray ionization (ESI). Analytical HPLC was carried out using a Phenomenex Kinetex C18 column (50 x 2.1 mm, 2 min gradient from 5% MeCN / 0.1% TFA in H_2_O to 100% MeCN / 0.1% TFA) with UV detection at 214 and 254 nm. Preparative reverse-phase HPLC was conducted on a Gilson system with a Phenomenex Luna C18 column (100 Å, 50 x 21.2 mm, 5 μm) using a 10 min gradient from 5 → 95% MeCN / 0.1% TFA.

All final compounds were determined to be ≥95% pure by LC/MS analysis.

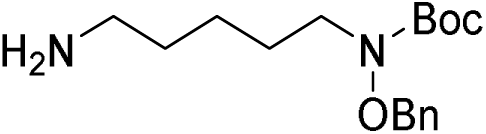

#### *tert*-Butyl (5-aminopentyl)(benzyloxy)carbamate (8)

Triphenylphosphine (1.33 g, 5.08 mmol) was added to a solution of tert-butyl (5-azidopentyl)(benzyloxy)carbamate (850 mg, 2.54 mmol) in THF (15 mL). The reaction mixture was stirred at 80 °C for 40 min (no more than an hour). Water (0.7 mL, 38.1 mmol) was added at room temperature and stirred at 80 °C for 45 min. The reaction mixture was concentrated. The residue was purified by silica gel chromatography with a Teledyne ISCO Combi-Flash (12 g column) eluting with 0 to 80 % MeOH in CH_2_Cl_2_ to provide the desired product (744 mg, 95% yield) as colorless liquid. LCMS ESI-MS(m/z) calc’d for C_17_H_28_N_2_O_3_ [M+H]^+^: 308.21 measured 309.1; ^1^H NMR (400 MHz, CDCl_3_) δ 7.44 – 7.31 (m, 5H), 4.82 (s, 2H), 3.40 (t, *J* = 7.2 Hz, 2H), 2.68 (t, *J* = 7.0 Hz, 2H), 1.55 (s, 6H), 1.50 (s, 12H), 1.37 – 1.27 (m, 2H).

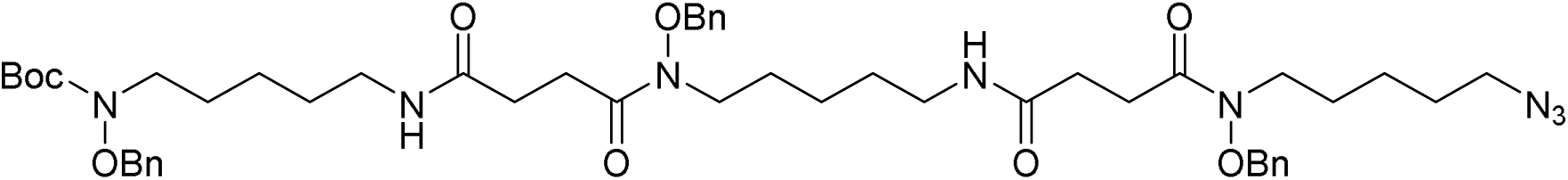

#### *tert*-Butyl (3-(5-azidopentyl)-14-(benzyloxy)-4,7,15,18-tetraoxo-1-phenyl-2-oxa-3,8,14,19-tetraazatetracosan-24-yl)(benzyloxy)carbamate (9)

*tert*-Butyl (5-aminopentyl)(benzyloxy)carbamate (262 mg, 848 μmol), EDC (244 mg, 1.27 mmol) and DMAP (51.8 mg, 424 μmol) were added to a solution of 3-(5-azidopentyl)-14-(benzyloxy)-4,7,15-trioxo-1-phenyl-2-oxa-3,8,14-triazaoctadecan-18-oic acid (**7**, 530 mg, 848 μmol) in CH_2_Cl_2_ (2.5 mL). The reaction mixture was stirred at room temperature for 20 h. The reaction mixture was diluted with CH_2_Cl_2_ and washed with sat. aq. NH_4_Cl solution. The organic layer was concentrated without drying (the solution was cloudy). The residue was purified by silica gel chromatography with a Teledyne ISCO Combi-Flash (24 g column) eluting with 0 to 100 % EtOAc in CH_2_Cl_2_ and then 0 to 20 % MeOH in CH_2_Cl_2_ to provide the desired product (610 mg, 79% yield) as yellow liquid. LCMS ESI-MS(m/z) calc’d for C_49_H_70_N_8_O_9_ [M+H]^+^: 914.53 measured 915.1; ^1^H NMR (400 MHz, CDCl_3_) δ 7.44 – 7.30 (m, 15H), 4.85 (s, 4H), 4.81 (s, 2H), 3.68 – 3.58 (m, 4H), 3.40 (q, *J* = 7.6 Hz, 3H), 3.27 – 3.13 (m, 6H), 2.83 - 2.75 (m, 4H), 2.48 (q, *J* = 7.2 Hz, 4H), 1.66 – 1.55 (m, 12H), 1.52.-1.45 (m, 11H), 1.37 – 1.24 (m, 6H).

**Compound 7** was prepared according to the procedure reported in Chiu et al, 2020^35^ (Scheme 1). All spectral data were consistent with those reported.

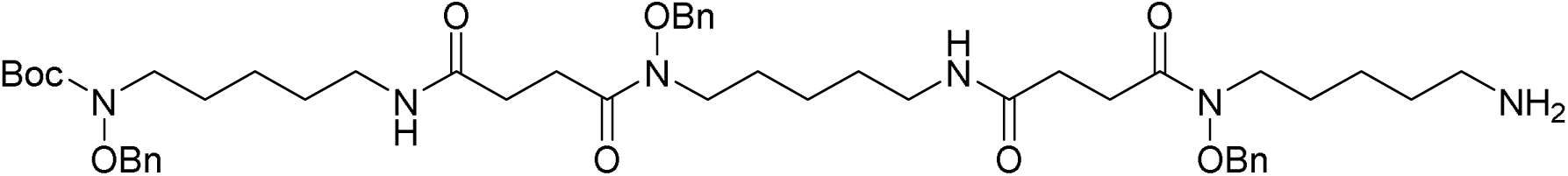

#### *tert*-Butyl (3-(5-aminopentyl)-14-(benzyloxy)-4,7,15,18-tetraoxo-1-phenyl-2-oxa-3,8,14,19-tetraazatetracosan-24-yl)(benzyloxy)carbamate

Triphenylphosphine (350 mg, 2 Eq, 1.33 mmol) was added to a solution of *tert*-butyl (3-(5-azidopentyl)-14-(benzyloxy)-4,7,15,18-tetraoxo-1-phenyl-2-oxa-3,8,14,19-tetraazatetracosan-24-yl) (benzyloxy)carbamate (610 mg, 667 μmol) in THF (6 mL). The reaction mixture was stirred at 80 °C for 50 min (no more than an hour). Water (0.2 mL, 10.0 mmol) was added at room temperature and stirred at 80 °C for 1 h. The reaction mixture was concentrated. The residue was purified by silica gel chromatography with a Teledyne ISCO Combi-Flash (12 g column) eluting with 0 to 100 % MeOH in CH_2_Cl_2_ to provide the desired product (215 mg, 36% yield) as pale yellow liquid. LCMS ESI-MS(m/z) calc’d for C_49_H_72_N_6_O_9_ [M+H]^+^: 888.54 measured 889.2; ^1^H NMR (400 MHz, CDCl_3_) δ 7.41 – 7.29 (m, 15H), 4.83 (d, *J* = 21.4 Hz, 6H), 3.73 – 3.54 (m, 4H), 3.44 – 3.33 (m, 4H), 3.26 – 3.04 (m, 4H), 2.73 (d, *J* = 43.8 Hz, 6H), 2.59 – 2.38 (m, 4H), 1.77 – 1.42 (m, 22H), 1.38 – 1.20 (m, 7H).

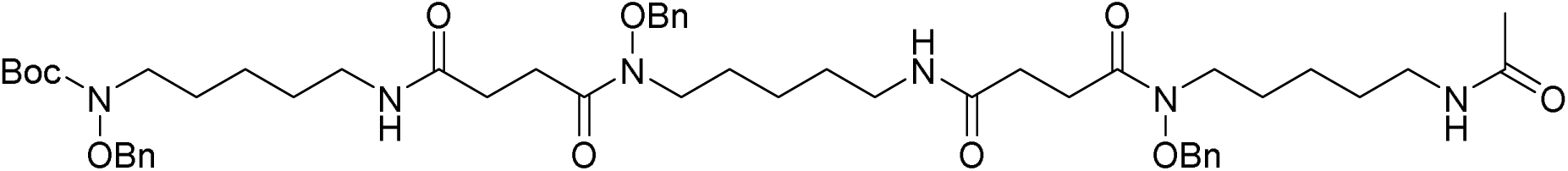

### *tert*-Butyl (benzyloxy)(9,20-bis(benzyloxy)-2,10,13,21,24-pentaoxo-3,9,14,20,25-pentaazatriacontan-30-yl)carbamate (10)

DMAP (5.91 mg, 48.4 μmol), acetic anhydride (34.3 μL, 363 μmol) and Et_3_N (101 μL, 725 μmol) were added to a solution of *tert*-butyl (3-(5-aminopentyl)-14-(benzyloxy)-4,7,15,18-tetraoxo-1-phenyl-2-oxa-3,8,14,19-tetraazatetracosan-24-yl) (benzyloxy)carbamate (215 mg, 242 μmol) in CH_2_Cl_2_ (5 mL) at 0 °C. The reaction mixture was stirred at room temperature for 15 min. Additional acetic anhydride (17 μL) and Et_3_N (50 μL) were added at 0 °C. The reaction mixture was stirred at room temperature for 4 h. Additional acetic anhydride (34.3 μL, 363 μmol) and Et_3_N (101 μL, 725 μmol) were added and stirred at room temperature for 16 h. The reaction mixture was concentrated. The residue was purified by silica gel chromatography with a Teledyne ISCO Combi-Flash (12g column) eluting with 0 to 20 % MeOH in CH_2_Cl_2_ to provide the desired product (210 mg, 93% yield) as yellow liquid. LCMS ESI-MS(m/z) calc’d for C_51_H_74_N_6_O_10_ [M+H]^+^: 930.55 measured 931.1; ^1^H NMR (400 MHz, CDCl_3_) δ 7.45 – 7.29 (m, 15H), 6.24 (d, *J* = 40.7 Hz, 3H), 4.88 – 4.75 (m, 6H), 3.63 (s, 4H), 3.39 (t, *J* = 7.3 Hz, 4H), 3.19 (td, *J* = 6.3, 3.0 Hz, 6H), 2.78 (s, 4H), 2.48 (q, *J* = 6.4 Hz, 4H), 2.09 (s, 6H), 1.95 (d, *J* = 2.8 Hz, 3H), 1.69 – 1.54 (m, 7H), 1.49 (s, 15H), 1.35 – 1.20 (m, 8H).

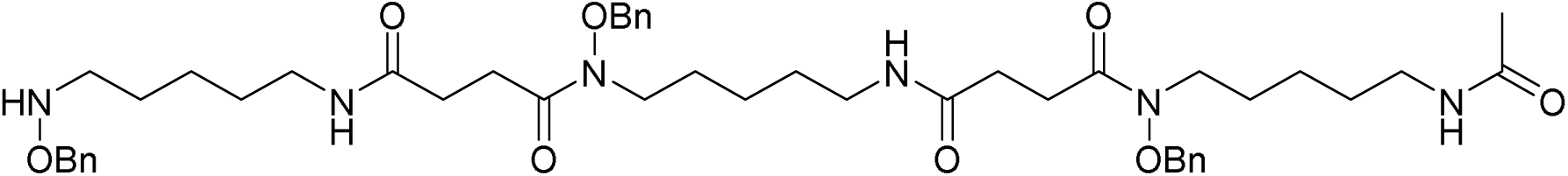

### *N*^1^-(5-Acetamidopentyl)-*N*^1^-(benzyloxy)-*N*^4^-(14-(benzyloxy)-10,13-dioxo-1-phenyl-2-oxa-3,9,14-triazanonadecan-19-yl)succinimide (11)

TFA (500 μL, 6.49 mmol) were added to a solution of *tert*-butyl (benzyloxy)(9,20-bis(benzyloxy)-2,10,13,21,24-pentaoxo-3,9,14,20,25-pentaazatriacontan-30-yl)carbamate (210 mg, 226 μmol) in CH_2_Cl_2_ (1.5 mL) at 0 °C. The reaction mixture was stirred at room temperature for 1 h. The reaction was diluted with CH_2_Cl_2_ and sat. aq. NaHCO_3_ solution. Extracted with CH_2_Cl_2_ (3 X 10 mL). The combined organic layers were dried (MgSO_4_), filtered and concentrated. The crude mixture (185 mg, 99% yield) was used for next step without further purification. LCMS ESI-MS(m/z) calc’d for C_46_H_66_N_6_O_8_ [M+H]^+^: 830.49 measured 831.1; ^1^H NMR (400 MHz, CDCl_3_) δ 7.39 – 7.28 (m, 15H), 6.30 – 6.10 (m, 3H), 4.84 (s, 2H), 4.83 (s, 2H), 4.69 (s, 2H), 3.73 – 3.54 (m, 4H), 3.29 – 3.11 (m, 6H), 2.91 (dd, *J* = 9.2, 5.0 Hz, 2H), 2.83 – 2.71 (m, 4H), 2.49 – 2.42 (m, 4H), 1.93 (s, 3H), 1.66 – 1.57 (m, 5H), 1.55 – 1.42 (m, 7H), 1.39 – 1.20 (m, 7H).

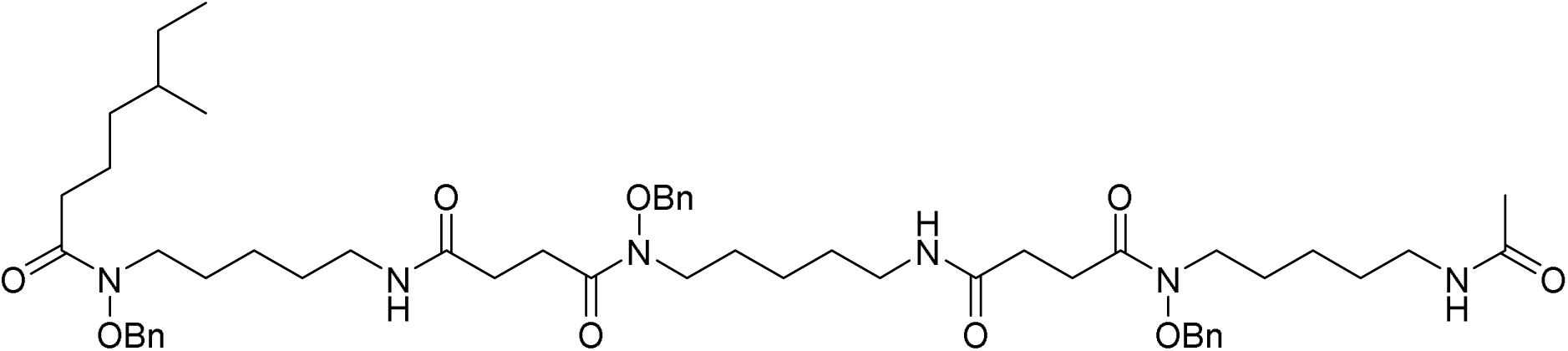

### *N*^1^-(5-acetamidopentyl)-*N*^1^-(benzyloxy)-*N*^4^-(14-(benzyloxy)-3-(5-methylheptanoyl)-10,13-dioxo-1-phenyl-2-oxa-3,9,14-triazanonadecan-19-yl)succinimide

5-Methylheptanoic acid (8.85 mg, 61.4 μmol), EDC (17.6 mg, 92.1 μmol) and DMAP (3.75 mg, 30.7 μmol) were added to a solution of *N*^1^-(5-acetamidopentyl)-*N*^1^-(benzyloxy)-*N*^4^-(14-(benzyloxy)-10,13-dioxo-1-phenyl-2-oxa-3,9,14-triazanonadecan-19-yl)succinamide (51.0 mg, 61.4 μmol) in CH_2_Cl_2_ (2 mL). The reaction mixture was stirred at room temperature for 21 h. The reaction mixture was concentrated. The residue was purified by silica gel chromatography with a Teledyne ISCO Combi-Flash (4 g column) eluting with 0 to 30% MeOH (10% MeOH in CH_2_Cl_2_) in CH_2_Cl_2_ and then 0 to 20% MeOH in CH_2_Cl_2_ to provide the desired product(44 mg, 76% yield) as colorless liquid. LCMS ESI-MS(m/z) calc’d for C_54_H_80_N_6_O_9_ [M+H]^+^: 956.60 measured 957.1; ^1^H NMR (400 MHz, CDCl_3_) δ 7.37 (d, *J* = 4.0 Hz, 15H), 6.35 – 6.19 (m, 3H), 4.85 (s, 2H), 4.83 (s, 2H), 4.80 (s, 2H), 3.69 – 3.57 (s, 6H), 3.21 – 3.15 (m, 6H), 2.81 – 2.75 (m, 4H), 2.50 – 2.45 (m, 4H), 2.36 (t, *J* = 7.7 Hz, 2H), 1.94 (s, 3H), 1.69 – 1.54 (m, 6H), 1.52 – 1.43 (m, 6H), 1.37 – 1.22 (m, 10H), 1.15 – 1.05 (m, 2H), 0.88 – 0.79 (m, 7H).

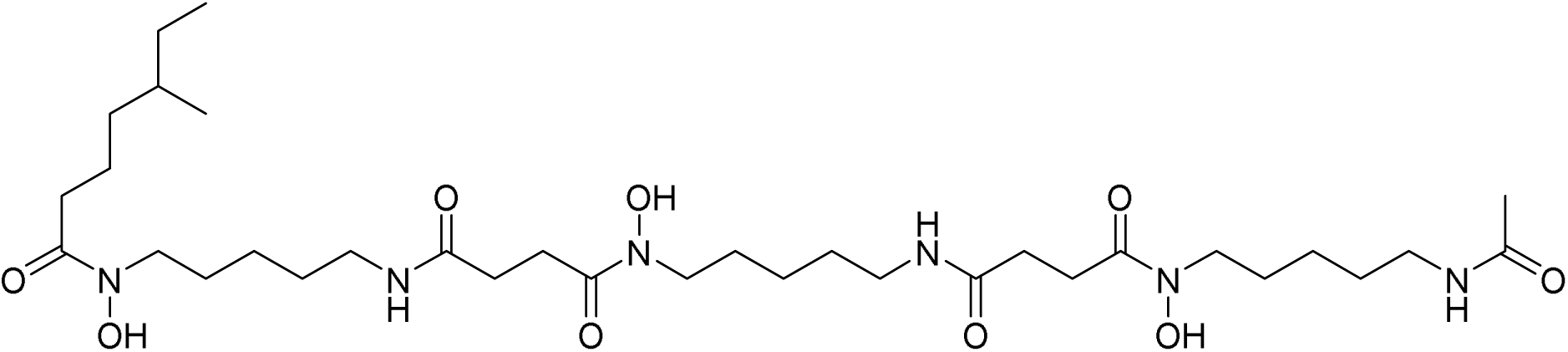

### *N*^1^-(5-Acetamidopentyl)-*N*^1^-hydroxy-*N*^4^-(5-(*N*-hydroxy-4-((5-(*N*-hydroxy-5-methylheptanamido)pentyl)amino)-4-oxobutanamido)pentyl)succinimide (VU0980806)

1-2 Drops of acetic acid was added to a solution of *N*^1^-(5-acetamidopentyl)-*N*^1^-(benzyloxy)-*N*^4^-(14-(benzyloxy)-3-(5-methylheptanoyl)-10,13-dioxo-1-phenyl-2-oxa-3,9,14-triazanonadecan-19-yl)succinamide (44 mg, 46 μmol) in MeOH (5 mL). The reaction vial was degassed with Ar (g). palladium on carbon (73 mg, 10% Wt, 69 μmol) was added. The reaction mixture was stirred at 45 °C for 3.5 h with H_2_ (g) balloon. The reaction solution was filtered through a syringe filter (washed with MeOH and CH_2_Cl_2_, more than 5 times). The solution was concentrated to provide the desired product (19 mg, 60% yield). LCMS ESI-MS(m/z) calc’d for C_33_H_62_N_6_O_9_ [M+H]^+^: 686.46 measured 392.1; ^1^H NMR (400 MHz, MeOH-d_4_) δ 3.60 – 3.57 (m, 4H), 3.17 – 3.13 (m, 6H), 2.76 (t, *J* = 7.2 Hz, 2H), 2.47 – 2.41 (m, 4H), 1.92 (d, *J* = 1.2 Hz, 3H), 1.72 – 1.45 (m, 14H), 1.39 – 1.29 (m, 12H), 1.21 – 1.05 (m, 6H), 0.94 – 0.76 (m, 11H); ^13^C NMR (101 MHz, MeOH-d_4_) δ 174.7, 173.4, 173.1, 173.0, 171.8, 171.5, 39.0, 38.9, 36.4, 35.8, 34.3, 34.1, 34.0, 32.3, 31.6, 30.2, 29.3, 29.2, 29.1, 28.9, 28.5, 28.4, 27.5, 25.9, 23.5, 23.4, 22.3, 22.2, 21.4, 18.3, 18.2, 10.5, 10.4.

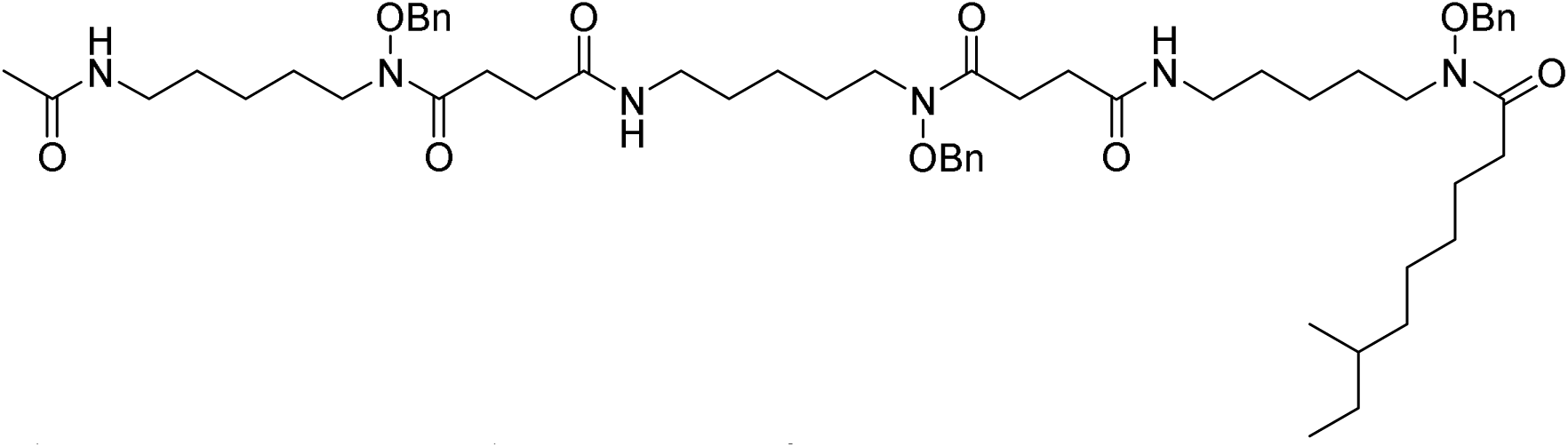

### *N*^1^-(5-acetamidopentyl)-*N*^1^-(benzyloxy)-*N*^4^-(14-(benzyloxy)-3-(7-methylnonanoyl)-10,13-dioxo-1-phenyl-2-oxa-3,9,14-triazanonadecan-19-yl)succinamide

To a solution of *N*^1^-(5-acetamidopentyl)-*N*^1^-(benzyloxy)-*N*^4^-(14-(benzyloxy)-10,13-dioxo-1-phenyl-2-oxa-3,9,14-triazanonadecan-19-yl)succinamide (43.0 mg, 51.7 μmol) in CH_2_Cl_2_ (2 mL) were added 7-Methyl-nonanoic acid, (8.91 mg, 1 Eq, 51.7 μmol), EDC (14.9 mg, 1.5 Eq, 77.6 μmol), and DMAP (3.16 mg, 0.5 Eq, 25.9 μmol). The reaction mixture was stirred at room temperature for 18 h. The reaction mixture was concentrated. The residue was purified by silica gel chromatography with a Teledyne ISCO Combi-Flash (4 g column) eluting with 0-7% MeOH/CH_2_Cl_2_ to provide *N*^1^-(5-acetamidopentyl)-*N*^1^-(benzyloxy)-*N*^4^-(14-(benzyloxy)-3-(7-methylnonanoyl)-10,13-dioxo-1-phenyl-2-oxa-3,9,14-triazanonadecan-19-yl)succinimide (44 mg, 70% yield) as colorless liquid. LCMS ESI-MS(m/z) calc’d for C_54_H_80_N_6_O_9_ [M+H]^+^: 985.32 measured 985.20; ^1^H NMR (400 MHz, CDCl_3_) δ 7.42-7.30 (m, 15H), 4.86 – 4.77 (m, 6H), 3.67-3.52 (m, 6H), 3.24-3.12 (m, 6H), 2.84-2.71 (m, 4H), 2.52-2.42 (m, 4H), 2.41-2.26 (m, 2H), 1.94 (s, 3H), 1.67-1.53 (m, 8H), 1.52-1.39 (m, 8H), 1.36 – 1.18 (m, 16H), 1.17 – 0.96 (m, 4H), 0.88-0.77 (m, 8H).

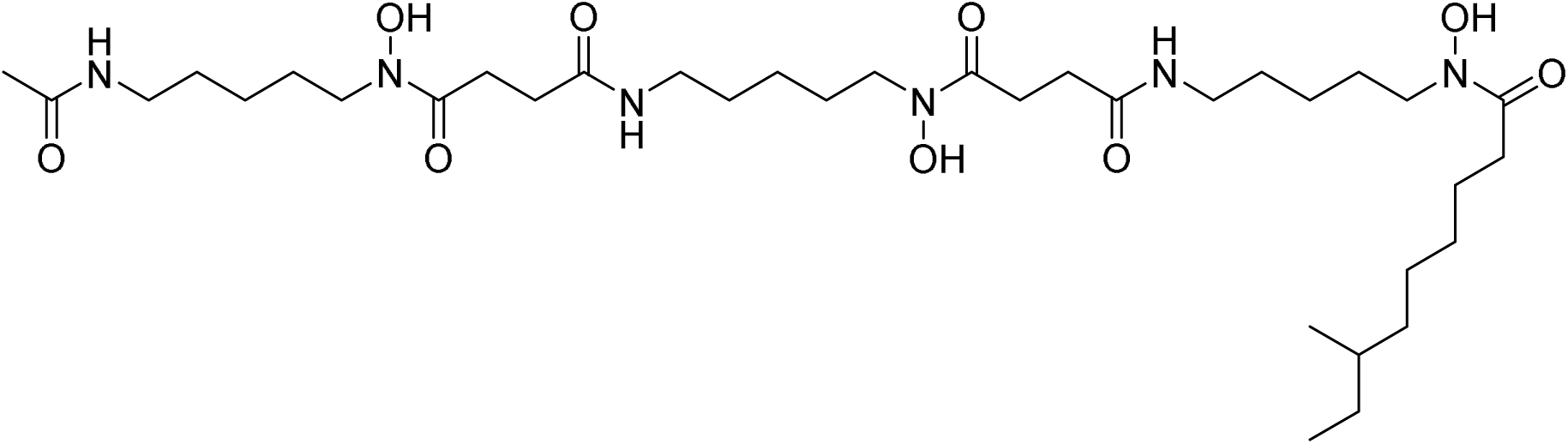

### *N*^1^-(5-acetamidopentyl)-*N*^1^-hydroxy-*N*^4^-(5-(N-hydroxy-4-((5-(*N*-hydroxy-7-methylnonanamido)pentyl)amino)-4-oxobutanamido)pentyl)succinimide (VU0980805)

To a solution of *N*^1^-(5-acetamidopentyl)-*N*^1^-(benzyloxy)-*N*^4^-(14-(benzyloxy)-3-(7-methylnonanoyl)-10,13-dioxo-1-phenyl-2-oxa-3,9,14-triazanonadecan-19-yl)succinamide (36.0 mg, 36.5 μmol) in MeOH (5.0 mL) was added 3 drops of acetic acid and palladium on carbon (58.3 mg, 10% Wt, 54.8 μmol). The reaction vial was degassed with Ar (g) for 15 min. The reaction mixture was stirred at 45 °C for 3.5 h with H_2_ (g) balloon. The reaction solution was filtered through a celite pad and washed with MeOH and CH_2_Cl_2_. The solution was concentrated to provide the *N*^1^-(5-acetamidopentyl)-*N*^1^-hydroxy-*N*^4^-(5-(N-hydroxy-4-((5-(*N*-hydroxy-7-methylnonanamido)pentyl)amino)-4-oxobutanamido)pentyl)succinimide (16.0 mg, 61% yield). ^1^H NMR (400 MHz, MeOH-d_4_) δ 3.67 – 3.56 (m, 4H), 3.23-3.11 (m, 4H), 2.83-2.71 (m, 2H), 2.53 – 2.42 (m, 2H), 2.229-2.14 (m, 2H), 1.95 (s, 3H), 1.73-1.47 (m, 8H), 1.43 – 1.25 (m, 10H), 1.22 – 1.06 (m, 2H), 0.96-0.82 (m, 4H).; ^13^C NMR (150 MHz, MeOH-d_4_) δ 180.1, 174.8, 174.6, 172.9, 49.3, 48.7, 39.9, 36.9, 33.0, 30.6, 30.5, 30.3, 30.0, 29.6, 29.0, 28.7, 28.5, 27.7, 26.8, 19.5, 11.5.

## Data Availability

All raw LC–MS/MS data and processed molecular networking results have been deposited in the Global Natural Products Social (GNPS) platform under the MassIVE accession number [to be assigned] and are publicly accessible (https://gnps2.org/dashboards/networkviewer/?usi=mzdata%3AGNPS2%3ATASK-111e11ec50814383affd7f06232c90d8-nf_output%2Fnetworking%2Fnetwork.graphml&usi-mgf=mzdata%3AGNPS2%3ATASK-111e11ec50814383affd7f06232c90d8-nf_output%2Fclustering%2Fspecs_ms.mgf#{}) or at (https://gnps.ucsd.edu/ProteoSAFe/result.jsp?view=network_displayer&componentindex=10&highlight_node=867&task=ac1ef22f5a3846998f3aaff091c2955f#%7B%7D) NMR spectra of the natural and synthetic siderophore analogs, as well as annotated spectra, are available in the Supporting Information. Additional experimental details, MIC assay raw values, and synthetic protocols, are also provided in the Supporting Information.

## Acknowledgements

Research reported in this publication was supported by the National Institute of General Medical Sciences under Award Number R35GM146987. The content is solely the responsibility of the authors and does not necessarily represent the official views of the National Institutes of Health. We gratefully acknowledge the contributions of Vanderbilt’s Advanced Computing Center for Research and Education (ACCRE), Mass Spectrometry Core Lab, Vanderbilt Small Molecule NMR Facility, Vanderbilt Biomolecular NMR Facility, and the Vanderbilt Molecular Design and Synthesis Center.

