## Supplemental for "Identification of Siderophores with Unexpected Antibacterial Properties from *Actinoplanes teichomyceticus*"

**Table of Contents**

**Figures**

**Fig. S1.** PCR analysis using agarose gel electrophoresis.

**Fig. S2.** Gene sequencing analysis of Pt-sol (nBLAST alignment results).

**Fig. S3.** Gene sequencing analysis of Pt-lq (nBLAST alignment results).

**Fig. S4.** Agar diffusion bioassay comparing crude extract (Pt ext) and aqueous fraction (Pt aq).

**Fig. S5.** HPLC chromatogram of crude extract from *Actinoplanes teichomyceticus* DSM 43866.

**Fig. S6.** Chrome Azurol S (CAS) agar assay for siderophore production.

**Fig. S7.** Comparative CAS activity assay of known siderophores and isolated compounds.

**Fig. S8.** Minimum inhibitory concentration (MIC) assay results in Mueller–Hinton Broth (Preliminary and Final Data).

**Fig. S9.** Comparative bioactivity of synthetic and ion-chelated siderophores.

**Fig. S10.** MIC evaluation under Fe- and Al-enriched conditions.

**Fig. S11.** Chemical structure and NMR characterization of synthetic hydroxamate siderophore analog VU0980805 (C9).

**Fig. S12.** ¹H and HSQC NMR spectra of synthesized C7.

**Tables**

**Table S1.** Machine learning models used to predict antibacterial activity probabilities for annotated antiSMASH 5.0 biosynthetic gene clusters in *A. teichomyceticus* DSM 43866.

**Table S2.** Diagnostic MS² fragments observed for aluminum- and iron-chelate siderophore analogs.

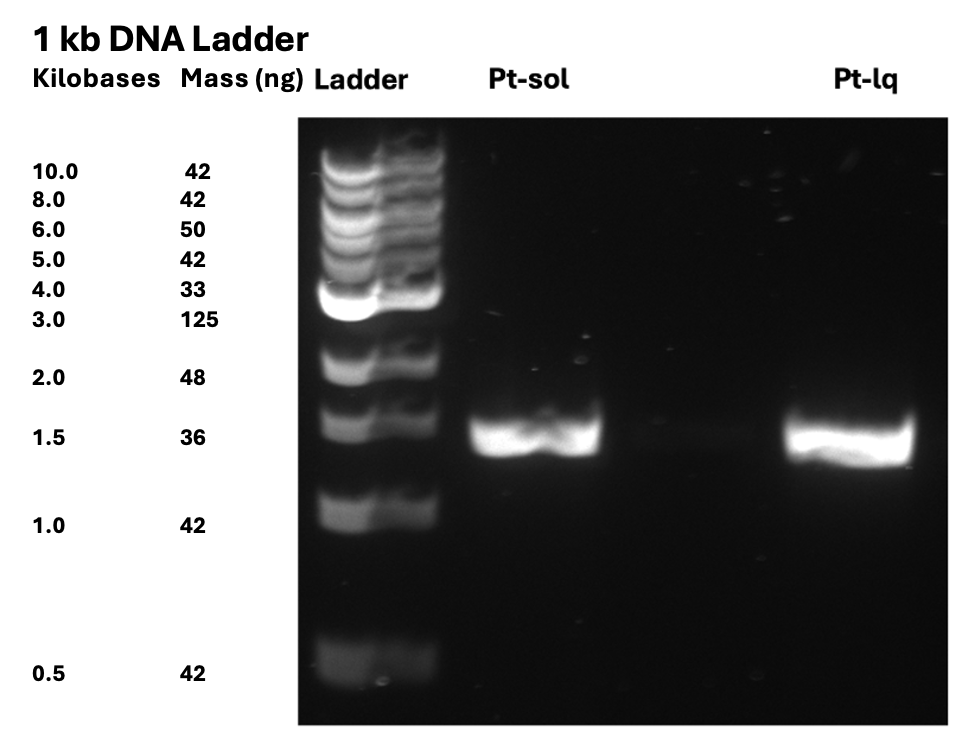

**Fig. S1**. *PCR analysis using agarose gel electrophores*is. The first lane contains a DNA size ladder serving as a molecular weight reference. **“Pt-sol”** denotes *Actinoplanes teichomyceticus* (Pt) DSM 43866 cultured on solid M65 agar medium, whereas **“Pt-lq**” represents the same strain grown in liquid M65 broth.

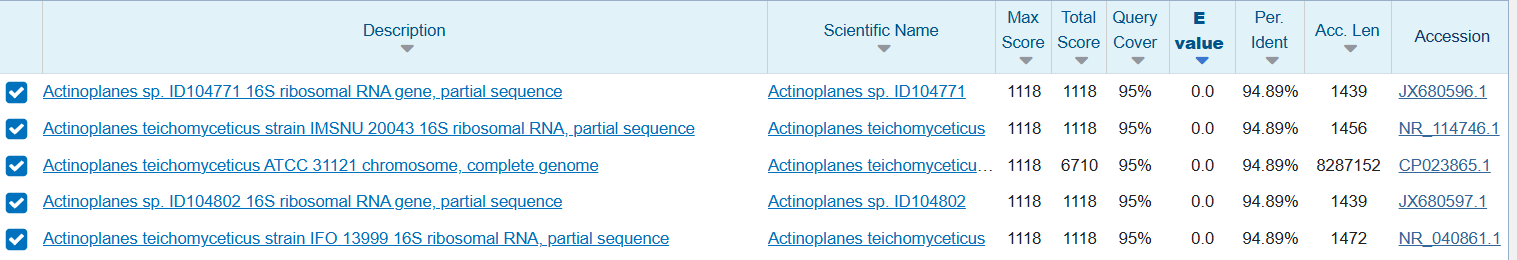
**Fig. S2**. *Gene sequencing analysis of* ***Pt-sol****.* nBLAST results indicate a strong sequence alignment with *Actinoplanes teichomyceticus* DSM 43866, showing 94.89% identity and 95% query coverage.

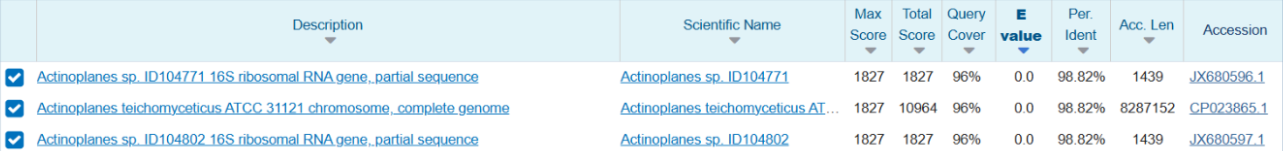

**Fig. S3**. *Gene sequencing analysis of* ***Pt-lq****.* nBLAST results indicate a strong sequence alignment with *Actinoplanes teichomyceticus* DSM 43866, showing 98.82% identity and 96% query coverage.

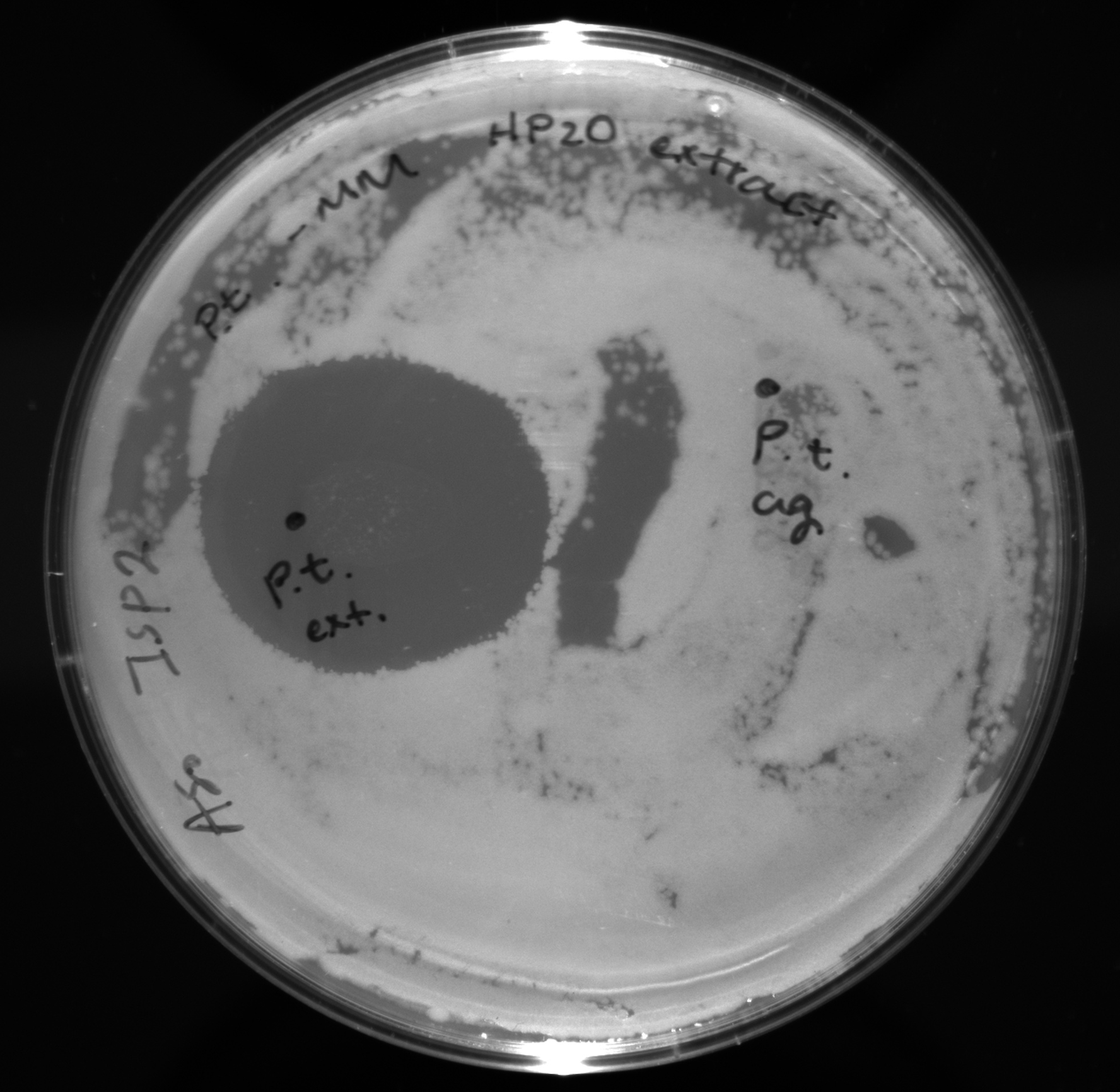

**Fig. S4.** *Agar diffusion bioassay comparing the crude extract (Pt ext) and aqueous fraction (Pt aq).* The antibacterial activity was evaluated against *Bacillus subtilis* (*Bacillus spizizenii* ATCC 6633) cultured on ISP2 agar medium.

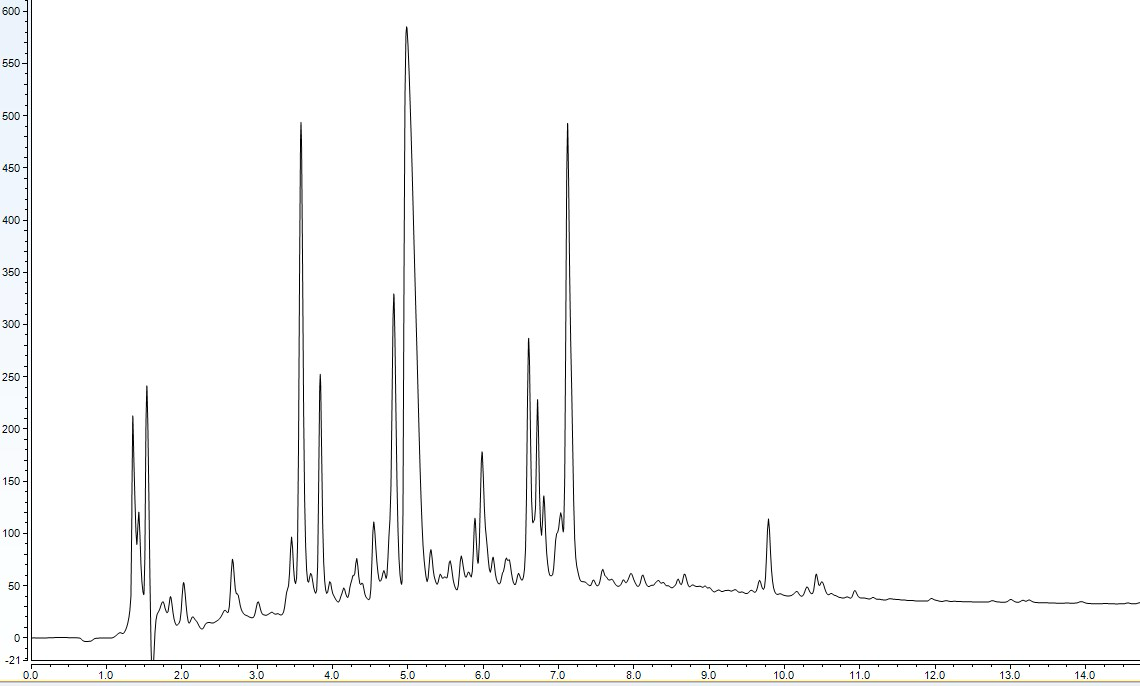

**Fig. S5.** High-performance liquid chromatography (HPLC) chromatogram of the 5 µL crude extract obtained from the matured M65 fermentation culture of *Actinoplanes teichomyceticus* DSM 43866. The analysis was performed using a reverse-phase C18 column under a gradient elution system optimized for secondary metabolite separation.

**Table S1.** Machine learning models used to predict antibacterial activity probabilities for annotated antiSMASH 5.0 biosynthetic gene clusters in *A. teichomyceticus* DSM 43866**.**

|  |  |  |  | **ML Prediction Probability Scores (Antibacterial Prediction)** | | |
| --- | --- | --- | --- | --- | --- | --- |
| **Region** | **BGC** | **BGC Type** | **Most Similar Known Cluster** | **Random Forest (Trees)** | **Logistic Regression** | **SVM** |
| Region 1.1 | VIWY01000001.1.region001 | Siderophore | Desferrioxamine E | 0.18 | 0.217 | 0.139 |
| Region 1.2 | VIWY01000001.1.region002 | T1PKS | Sceliphrolactam | 0.62 | 0.61 | 0.623 |
| Region 1.3 | VIWY01000001.1.region003 | T3PKS | Alkyl-O-dihydrogeranyl-methoxyhydroquinones | 0.7 | 0.467 | 0.653 |
| Region 1.4 | VIWY01000001.1.region004 | Nucleoside | Toyocamycin | 0.66 | 0.551 | 0.634 |
| Region 1.5 | VIWY01000001.1.region005 | Terpene | Isorenieratene | 0.18 | 0.252 | 0.142 |
| Region 2.1 | VIWY01000002.1.region001 | Indole | Erdasporine A/B/C | 0.56 | 0.515 | 0.396 |
| Region 2.2 | VIWY01000002.1.region002 | Terpene | 2-Methylisoborneol | 0.52 | 0.383 | 0.582 |
| Region 2.3 | VIWY01000002.1.region003 | NRPS, Beta-lactone | Paulomycin | 0.74 | 0.5 | 0.612 |
| Region 2.4 | VIWY01000002.1.region004 | LAP | Not Identified | 0.565 | 0.433 | 0.73 |
| Region 2.5 | VIWY01000002.1.region005 | Aminoglycoside/Cyclitol | Not Identified | 0.76 | 0.565 | 0.902 |
| Region 2.6 | VIWY01000002.1.region006 | Other | Eponemycin | 0.68 | 0.731 | 0.795 |
| Region 3.1 | VIWY01000003.1.region001 | Melanin | Not Identified | 0.58 | 0.531 | 0.784 |
| Region 3.2 | VIWY01000003.1.region002 | Lanthipeptide | Catenulipeptin | 0.52 | 0.693 | 0.709 |
| Region 3.3 | VIWY01000003.1.region003 | T1PKS, NRPS-like | Divergolide A–D | 0.7 | 0.472 | 0.609 |
| Region 3.4 | VIWY01000003.1.region004 | NRPS-like | Vazabitide A | 0.801 | 0.743 | 0.658 |
| Region 3.5 | VIWY01000003.1.region005 | Oligosaccharide, T1PKS | Maduropeptin | 0.78 | 0.836 | 0.718 |
| Region 4.1 | VIWY01000004.1.region001 | Phosphoglycolipid | Teichomycin | 0.78 | 0.621 | 0.905 |
| Region 5.1 | VIWY01000005.1.region001 | NRPS | BD-12 | 0.56 | 0.472 | 0.485 |
| Region 5.2 | VIWY01000005.1.region002 | Bacteriocin | Not Identified | 0.786 | 0.464 | 0.888 |
| Region 5.3 | VIWY01000005.1.region003 | Lanthipeptide | Microbisporicin A2 | 0.86 | 0.639 | 0.646 |
| Region 5.4 | VIWY01000005.1.region004 | Siderophore | Not Identified | 0.2 | 0.362 | 0.104 |
| Region 6.1 | VIWY01000006.1.region001 | NRPS | Teicoplanin | 0.56 | 0.525 | 0.78 |
| Region 6.2 | VIWY01000006.1.region002 | NRPS, T1PKS | Antimycin | 0.48 | 0.557 | 0.662 |
| Region 6.3 | VIWY01000006.1.region003 | NRPS, T1PKS | Aristeromycin | 0.68 | 0.723 | 0.694 |
| Region 8.1 | VIWY01000008.1.region001 | T3PKS, NRPS | Teicoplanin | 0.96 | 0.878 | 0.881 |
| Region 9.1 | VIWY01000009.1.region001 | NRPS | Ulleungmycin | 0.6 | 0.516 | 0.667 |

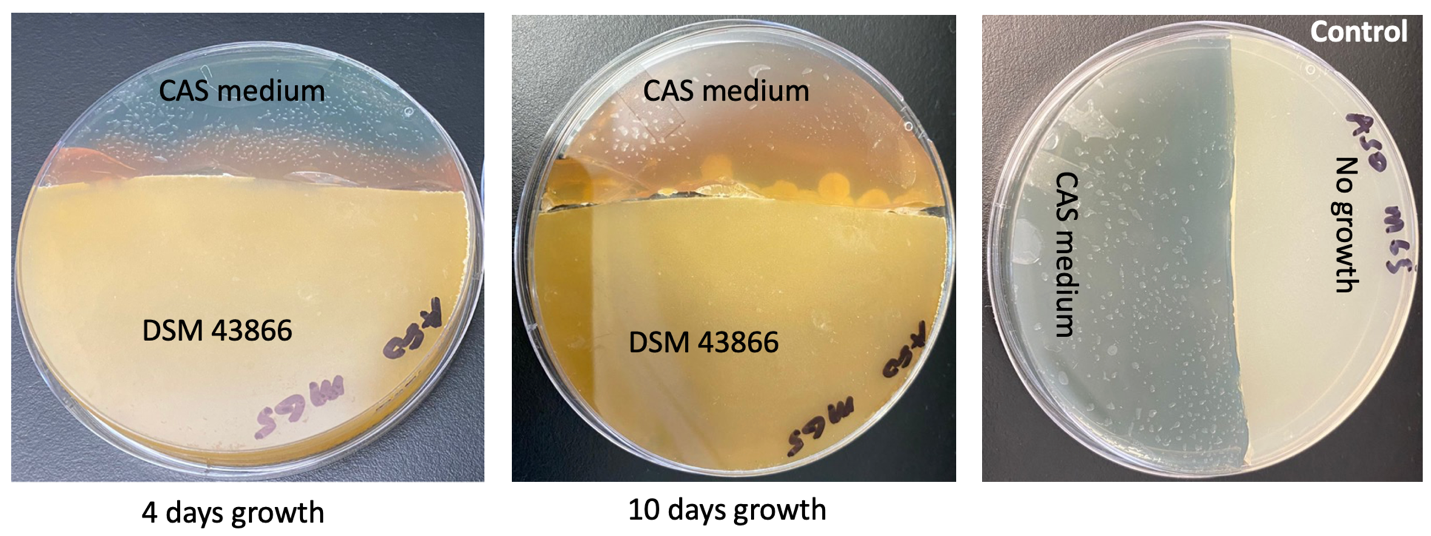

**Fig. S6***. Chrome Azurol S (CAS) agar assay for siderophore production by* *Actinoplanes teichomyceticus* DSM 43866. The CAS agar plate was incubated alongside an M65 agar plate inoculated with *A. teichomyceticus* DSM 43866. Visible color changes indicating siderophore activity were observed after 4 and 10 days of growth, compared with a control plate lacking bacterial inoculation

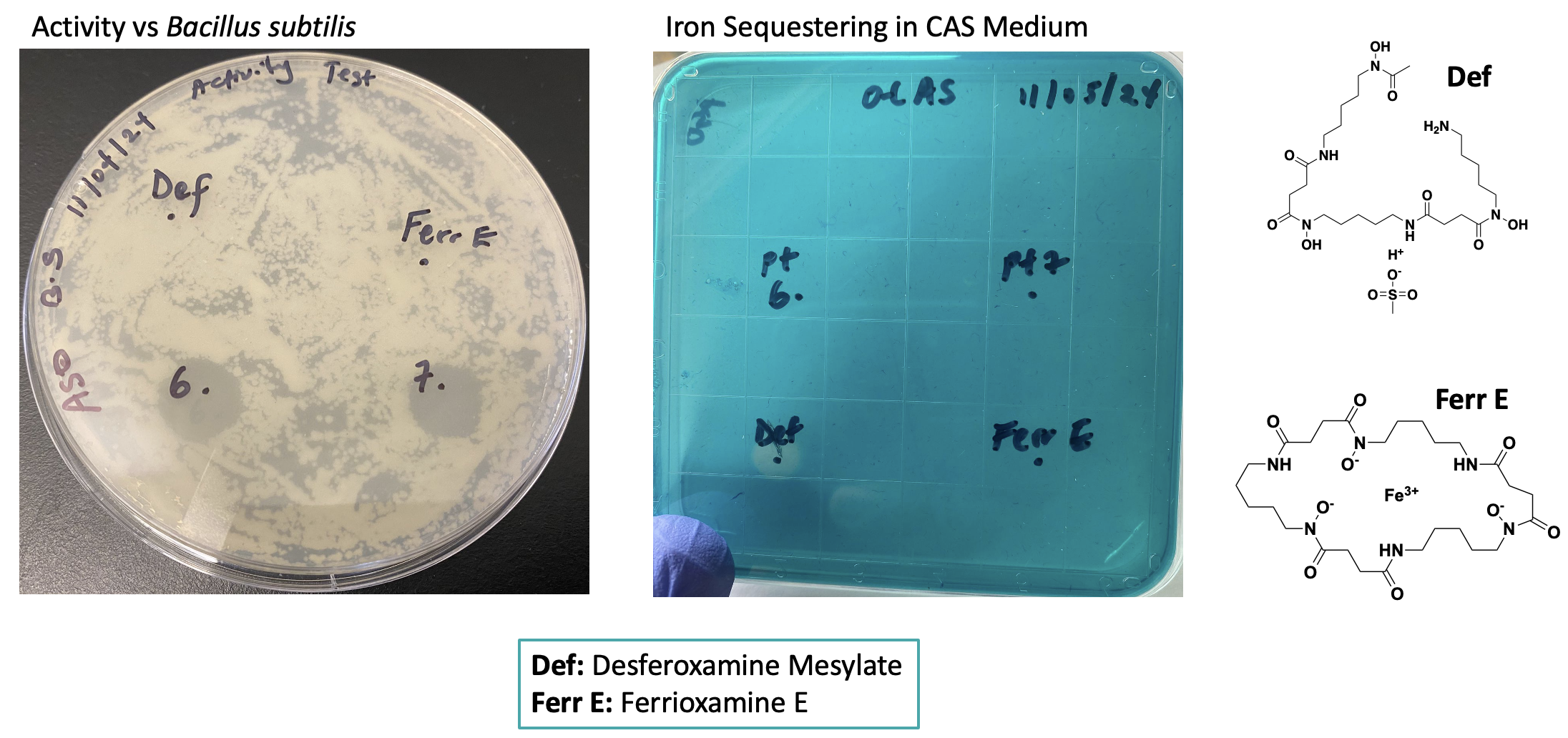

**Fig. S7.** *Comparative CAS activity assay for siderophore detection.* Known standards—desferrioxamine mesylate (**Def**) and ferrioxamine E (Ferr E)—were compared with the unknown isolated compounds (Pt-6 and Pt-7). The assay on the right evaluates the ion-chelating capacity of the isolated metabolites relative to the known siderophores.

A. First trial MIC experiment

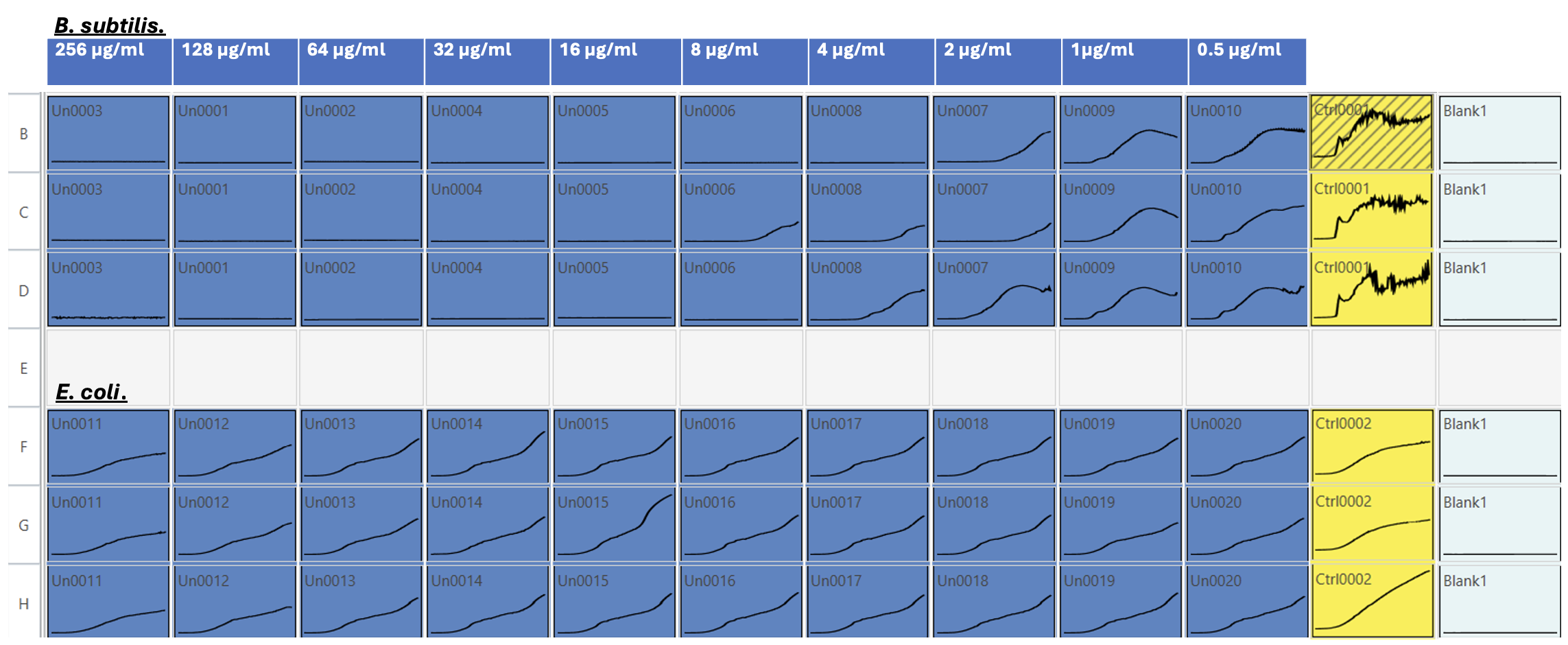

B. Second trail MIC experiment

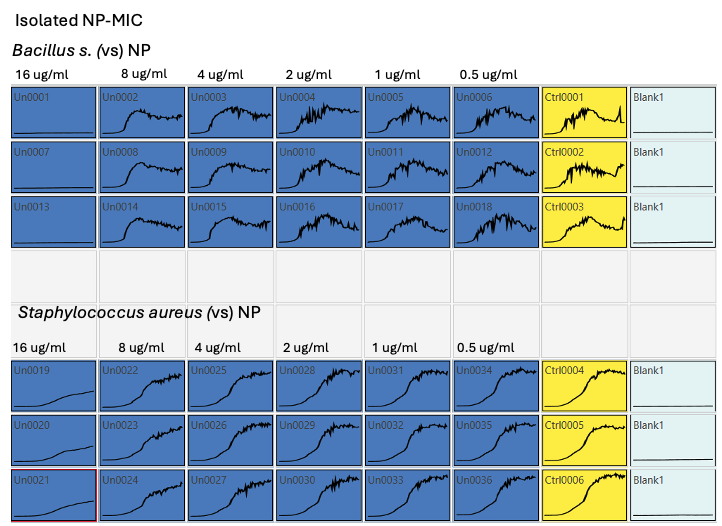

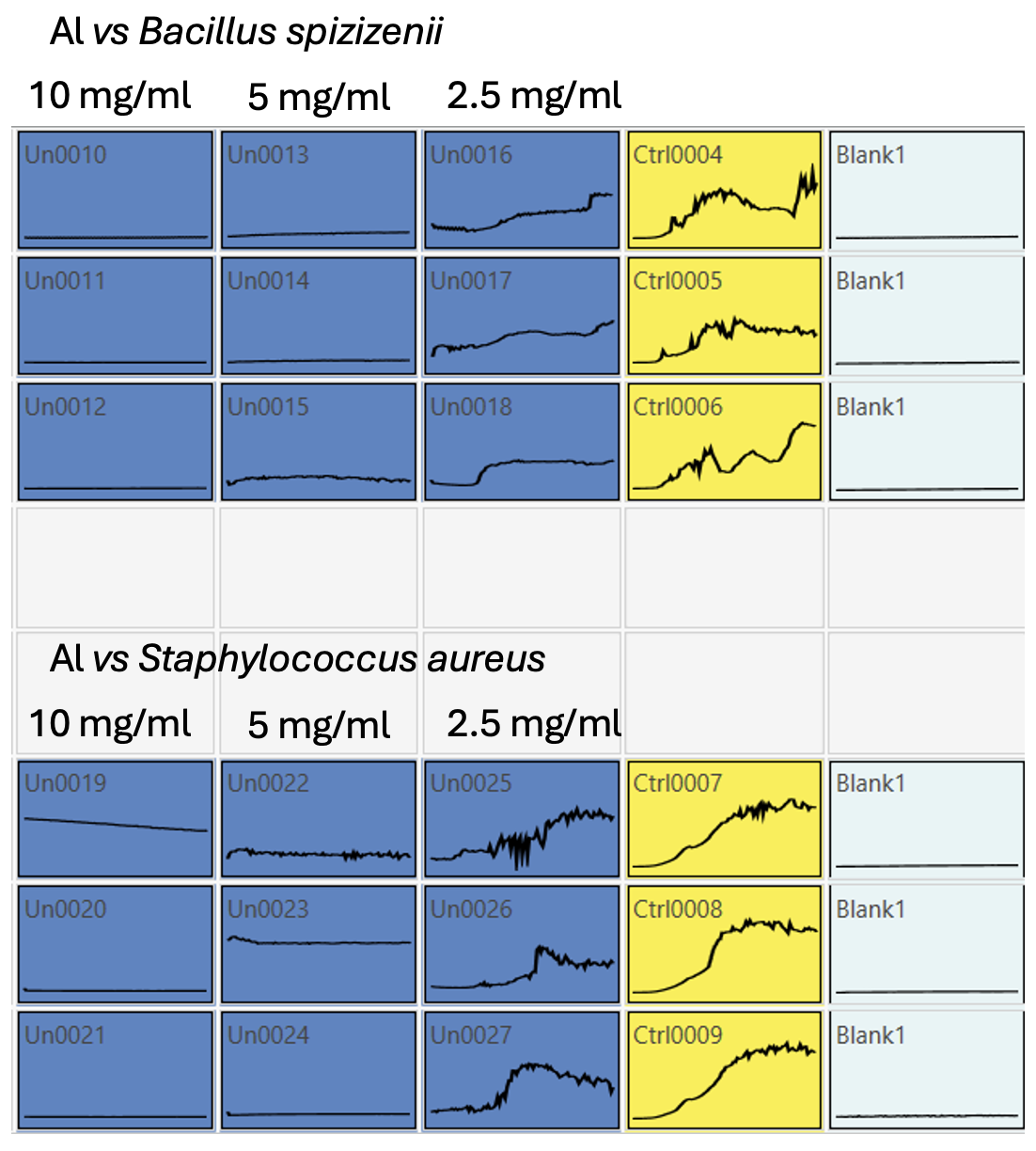

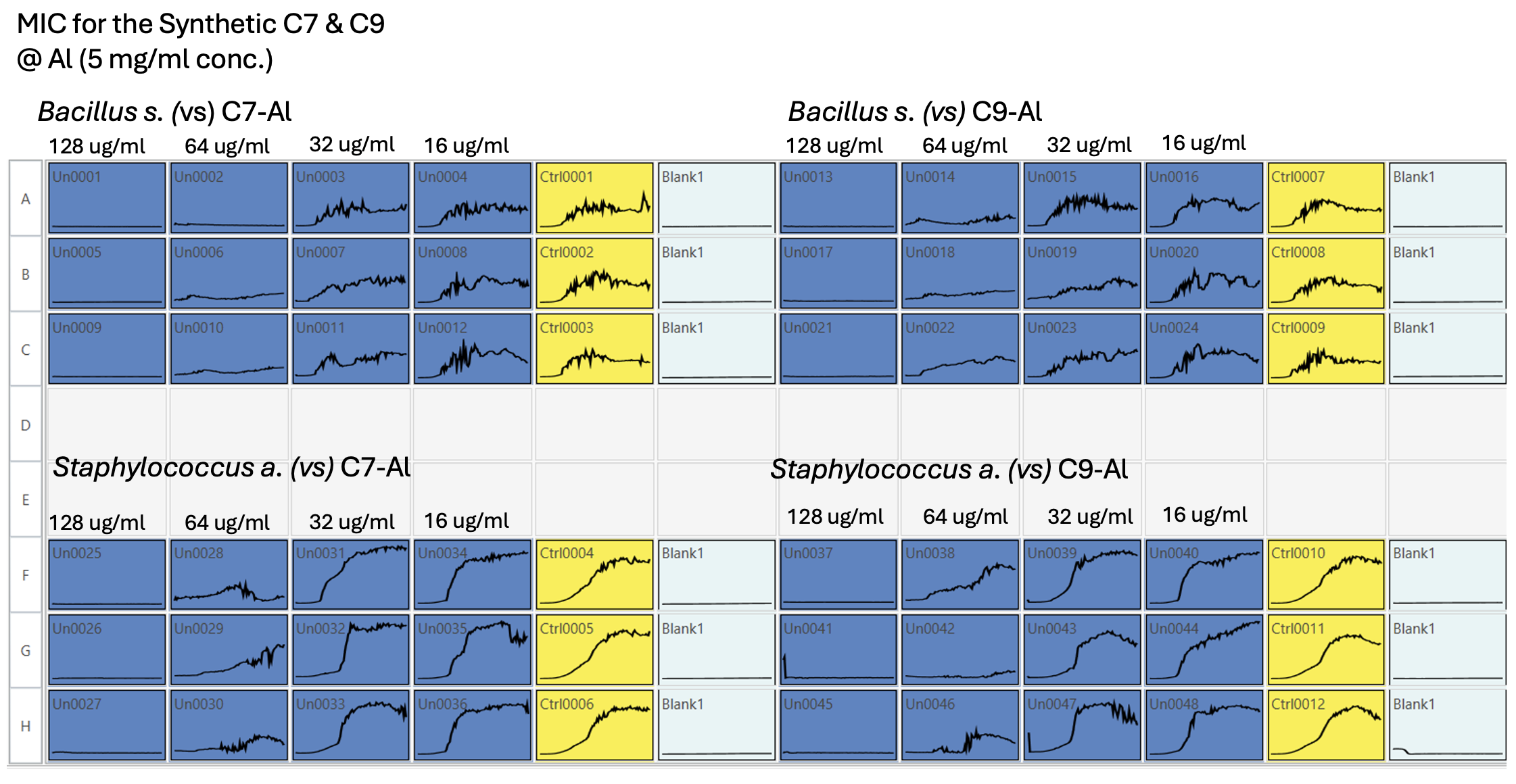

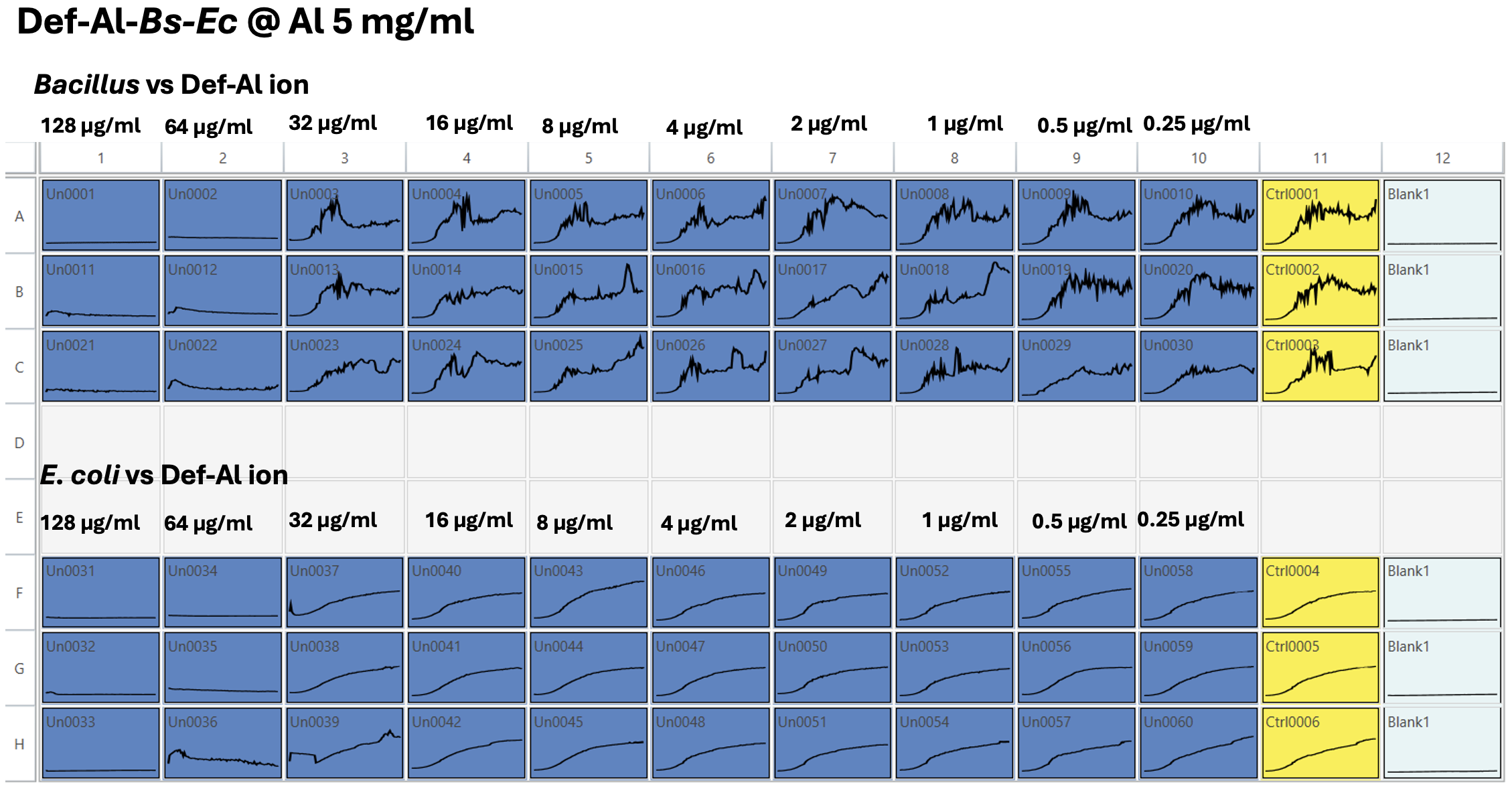

**Fig. S8. Minimum inhibitory concentration (MIC) assay results in Mueller–Hinton broth (MHB).**

(A) Preliminary plate reader data showing the antibacterial activity of the isolated hydroxamate-like siderophore against *Bacillus subtilis* (*Bacillus spizizenii* ATCC 6633) and Escherichia coli across a concentration range of 256 µg/mL to 0.5 µg/mL.

(B) Final plate reader data showing antibacterial activity of the isolated hydroxamate-like siderophore against Bacillus subtilis (*Bacillus spizizenii* ATCC 6633) and *Staphylococcus aureus* across a concentration range of 16 µg/mL to 0.5 µg/mL. Control (MHB + cells) and blank (medium only) readings are included for comparison.

(C) MIC of aluminum alone, starting at 10 mg/mL and serially diluted to 1.25 mg/mL, tested against *Bacillus spizizenii* and *Staphylococcus aureus* as a control.

(D) MIC of synthetic compounds C7 and C9 chelating aluminum ions, with an initial aluminum concentration of 5 mg/mL, serially diluted and tested against *Bacillus spizizenii* and *Staphylococcus aureus*.

(E) MIC of commercially available desferrioxamine mesylate chelating aluminum ions, with an initial aluminum concentration of 5 mg/mL, serially diluted and tested against *Bacillus spizizenii* and *E. coli*.

**Table S2.** Diagnostic MS² Fragments Observed for Aluminum- and Iron-Chelate Siderophore Analogs

| **Precursor Ion (m/z)** | **Molecular Formula** | **Diagnostic Fragment (m/z)** | **Fragment Formula** | **Notes** |
| --- | --- | --- | --- | --- |
| **697.4444 [M+H]+** | C_33_H_62_AlN_6_O_8_^+^ | 680.4177 | C₃₃H₅₉AlN₅O₈^•^ | Loss of NH group |
|  |  | 612.3538 | C₂₈H₅₁AlN₅O₈⁺ | Backbone cleavage |
|  |  | 581.3491 | C₂₈H₅₀AlN₄O₇⁺ | Loss of NO + side-chain |
|  |  | 543.3083 | C₂₃H₄₄AlN₆O₇⁺ | Multiple bond cleavage |
|  |  | 497.3277 | C₂₄H₄₆AlN₄O₅⁺ | Truncated hydroxamate-containing fragment |
| **711.4244 [M+H]+** | C_33_H_60_AlN_6_O_9_^+^ | 669.4119 | C₃₁H₅₈AlN₆O₈⁺ | Loss of C₂H₂O |
|  |  | 553.3178 | C₂₆H₄₆AlN₄O₇⁺ | Diagnostic hydroxamate cleavage |
|  |  | 469.2966 | C₂₂H₄₂AlN₄O₅⁺ | Core siderophore fragment |
|  |  | 385.2032 | C₁₆H₃₀AlN₄O₅⁺ | Terminal hydroxamate fragment |
| **740.3776 [M+H]+** | C_33_H_60_FeN_6_O_9_^+^ | 611.2507 | C₂₆H₄₅FeN₅O₈⁺ | Ferrioxamine-type diagnostic ion |
|  |  | 582.2715 | C₂₆H₄₆FeN₄O₇⁺ | Fragment consistent with hydroxamate cleavage |
|  |  | 414.1565 | C₁₆H₃₀FeN₄O₅⁺ | Core Fe–hydroxamate ion |
|  |  | 498.2503 | C₂₂H₄₂FeN₄O₅⁺ | Characteristic ferrioxamine scaffold fragment |

A.

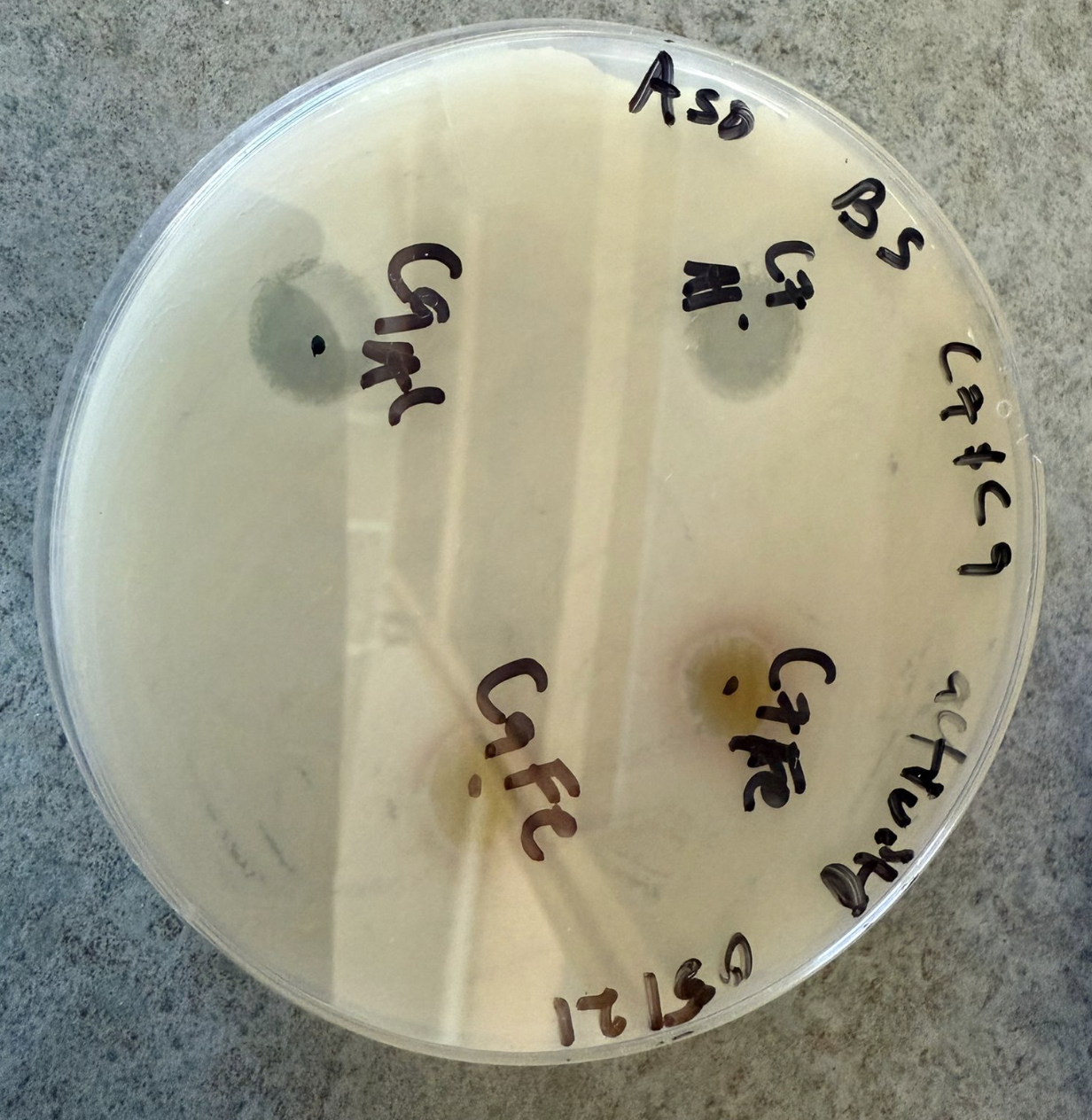

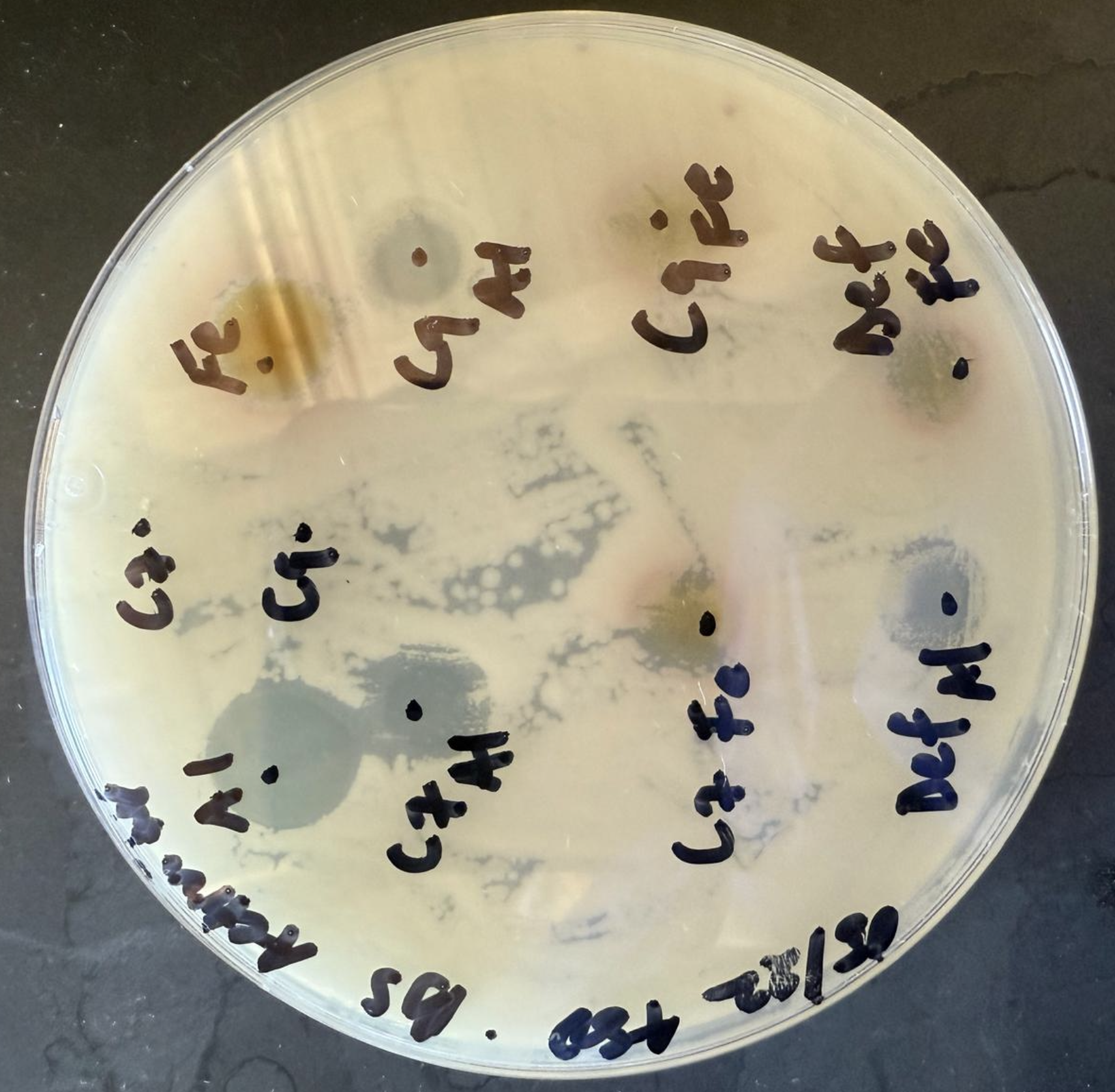

B.

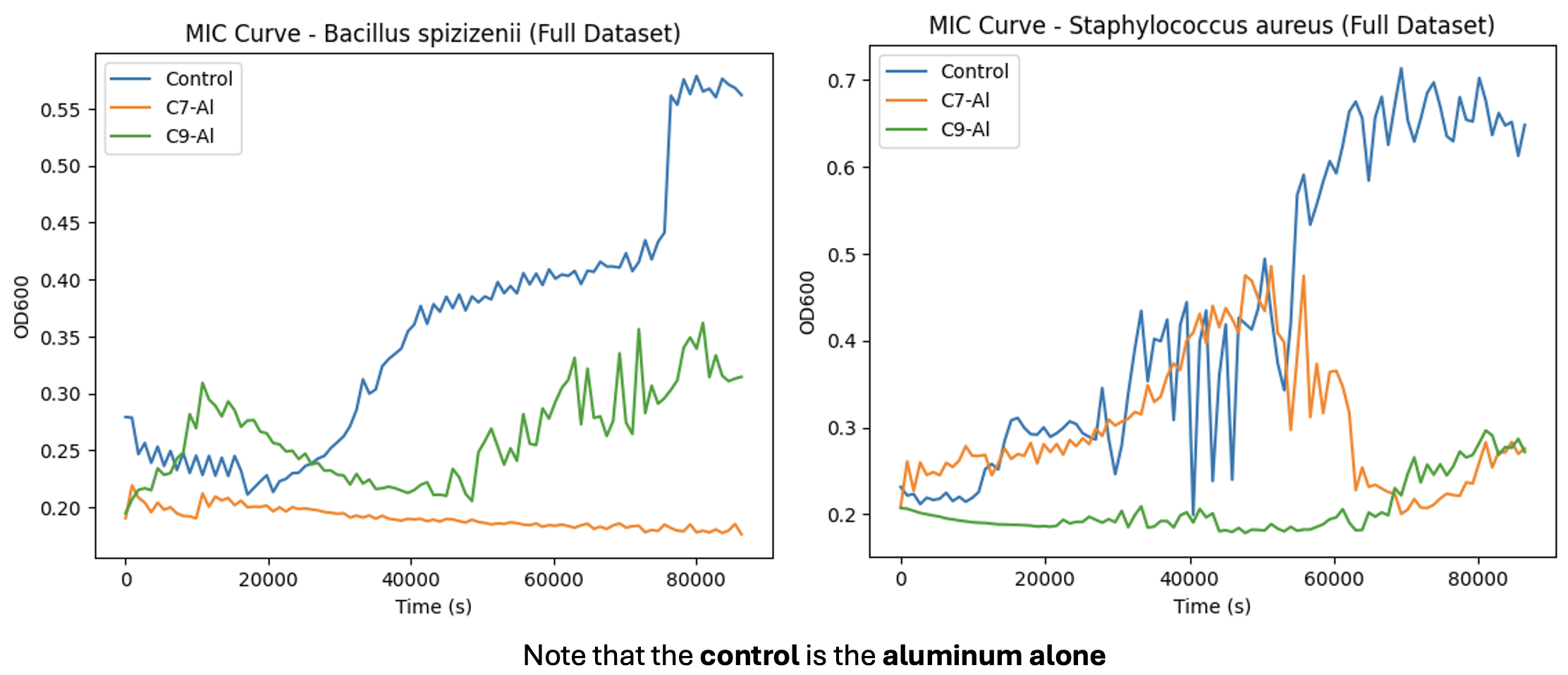

**Fig. S9*.*** **(A)** *Comparative bioactivity of synthetic and ion-chelated siderophores.* Synthetic siderophores C7 and C9 were compared with their aluminum-chelated counterparts (C7–Al and C9–Al). The Al-chelated forms exhibited enhanced antibacterial activity in the agar diffusion bioassay**. (B)** MIC growth curves of synthetic compounds C7 and C9 in complex with aluminum, showing growth inhibition at 64 µg/mL against *Bacillus spizizenii* and *Staphylococcus aureus*. The control condition (Al-alone) contains aluminum at 2.5 mg/mL.

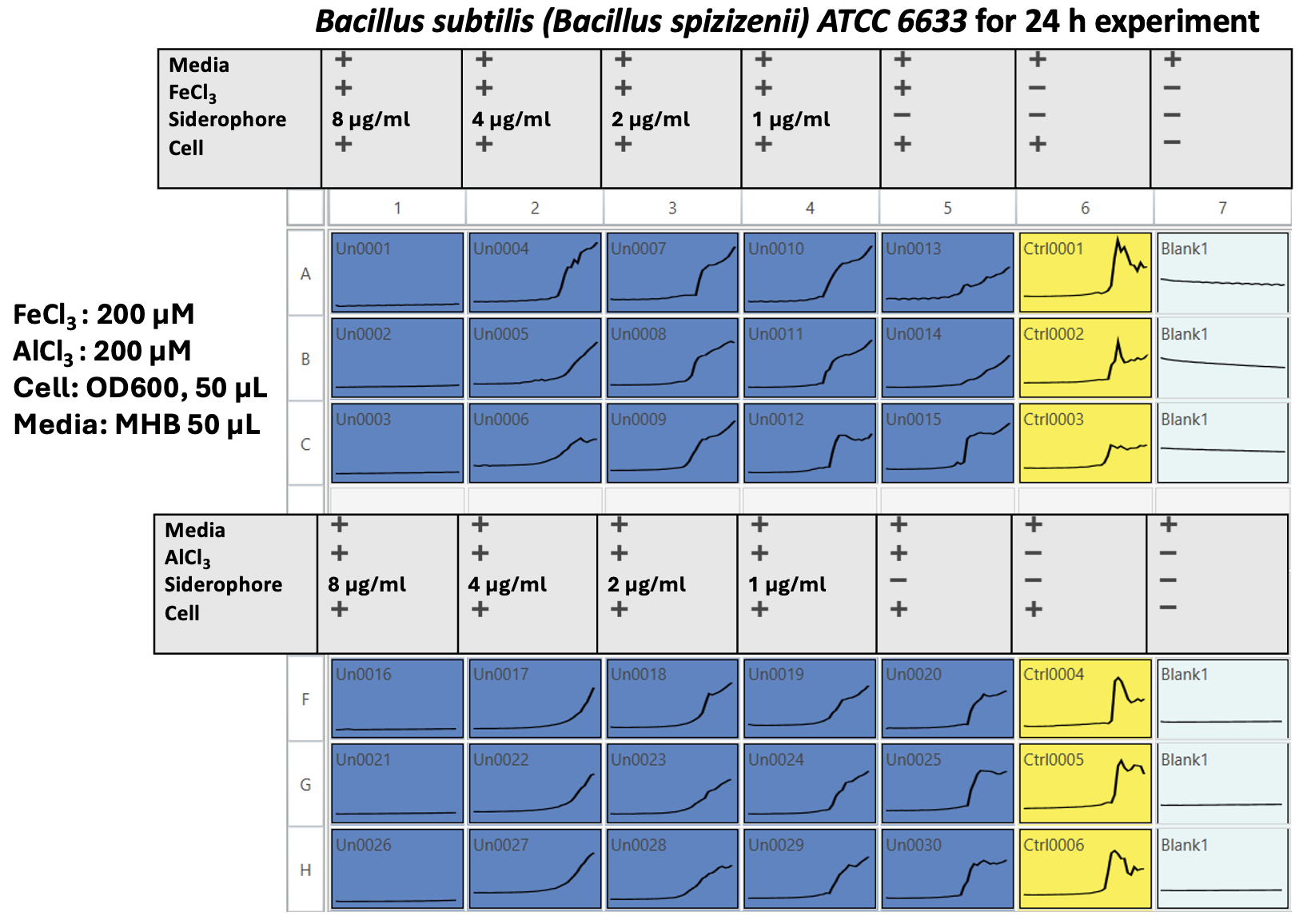

**Fig. S10.** Evaluation of the minimum inhibitory concentration (MIC) of the isolated siderophore under Fe- and Al-enriched conditions in Mueller-Hinton Broth (MHB). The siderophore was tested at an initial concentration of 8 µg/mL with serial dilutions down to 1 µg/mL against *Bacillus subtilis (Bacillus spizizenii)* ATCC 6633 over a 24-hour incubation period. The assay medium was supplemented with either FeCl₃ or AlCl₃ at a final concentration of 200 µM to assess the effect of metal enrichment on siderophore activity.

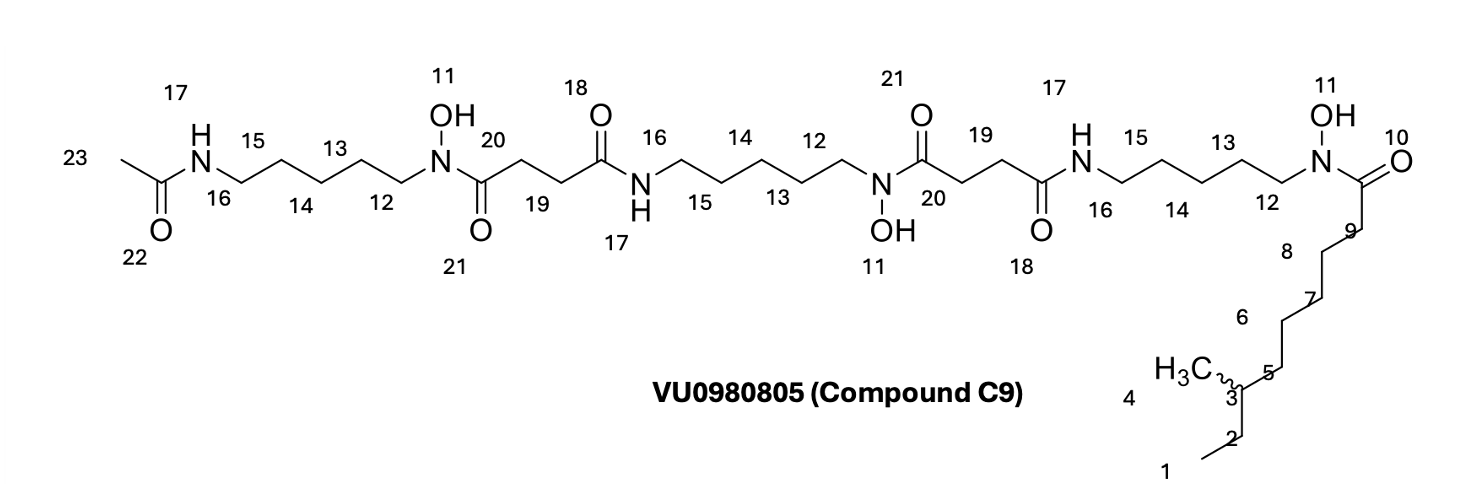

| Position | ^1^H NMR (ppm) | ^13^C NMR (ppm) |
| --- | --- | --- |
| 1 | 0.857 | 11.5 |
| 2 | 1.11 | 30.58 |
| 3 | 1.331 | 30.32 |
| 4 | 0.864 | 19.5 |
| 5 | 1.6 | 27.7 |
| 6 | 2.78 | 28.5 |
| 7 | 2.44 | 33 |
| 8 | 1.56 | 30 |
| 9 | 2.21 | 36.9 |
| 10 | N/A | 180.1 |
| 11 | N/A | N/A |
| 12 | 3.57 | 48.7 |
| 13 | 1.623 | 26.8 |
| 14 | 1.305 | 30.5 |
| 15 | 1.513 | 29.6 |
| 16 | 3.16 | 39.9 |
| 17 | N/A | N/A |
| 18 | N/A | 174.6 |
| 19 | 2.564 | 29 |
| 20 | 2.77 | 28.7 |
| 21 | N/A | 174.8 |
| 22 | N/A | 172.9 |
| 23 | 3.3 | 49.3 |

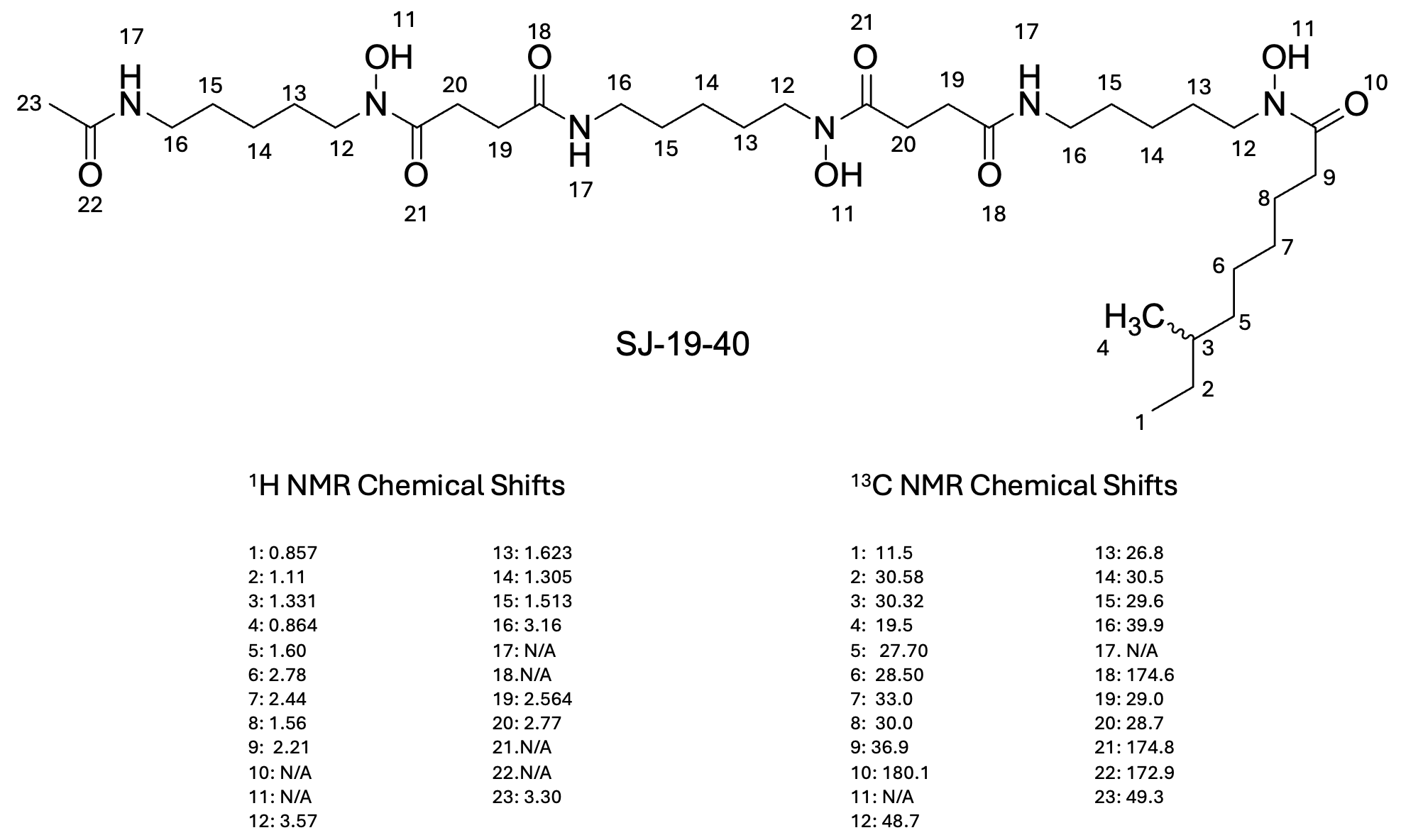

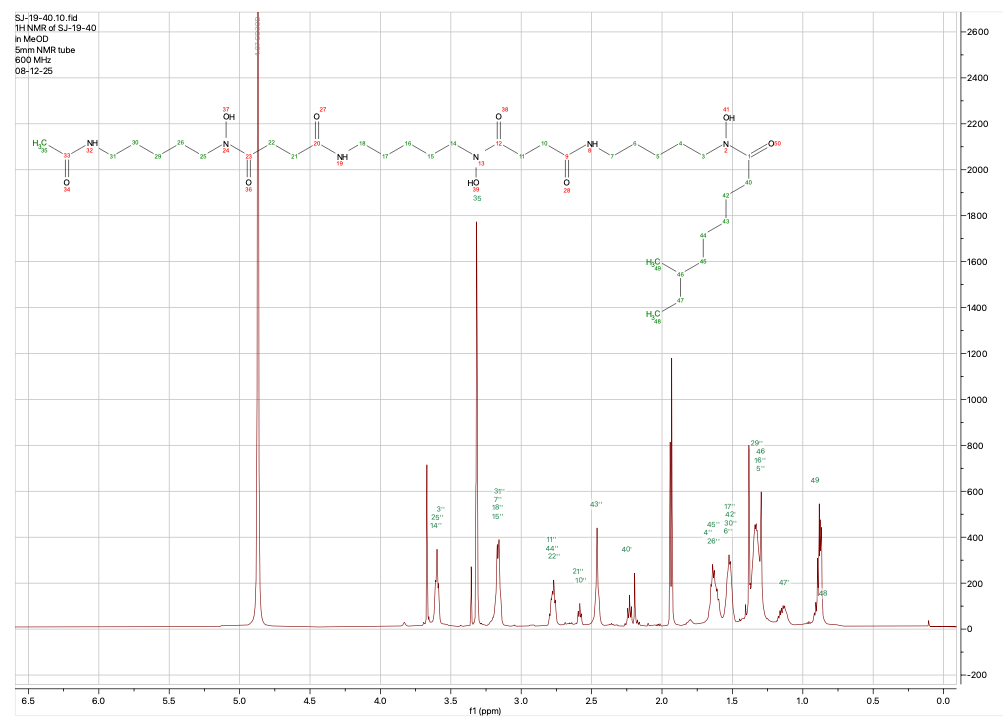

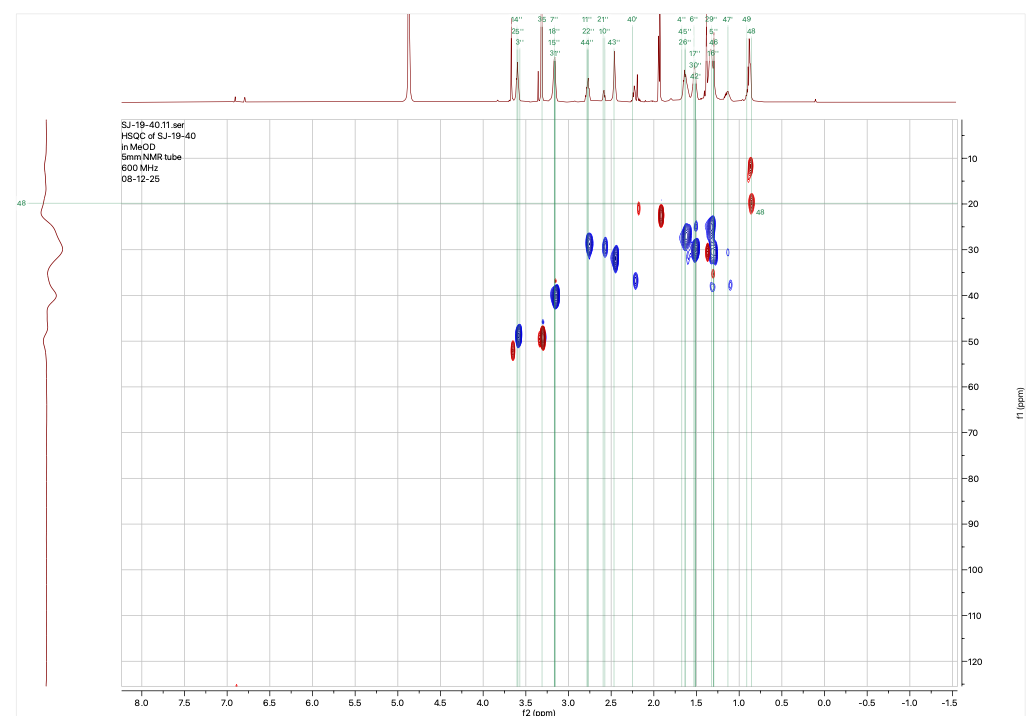

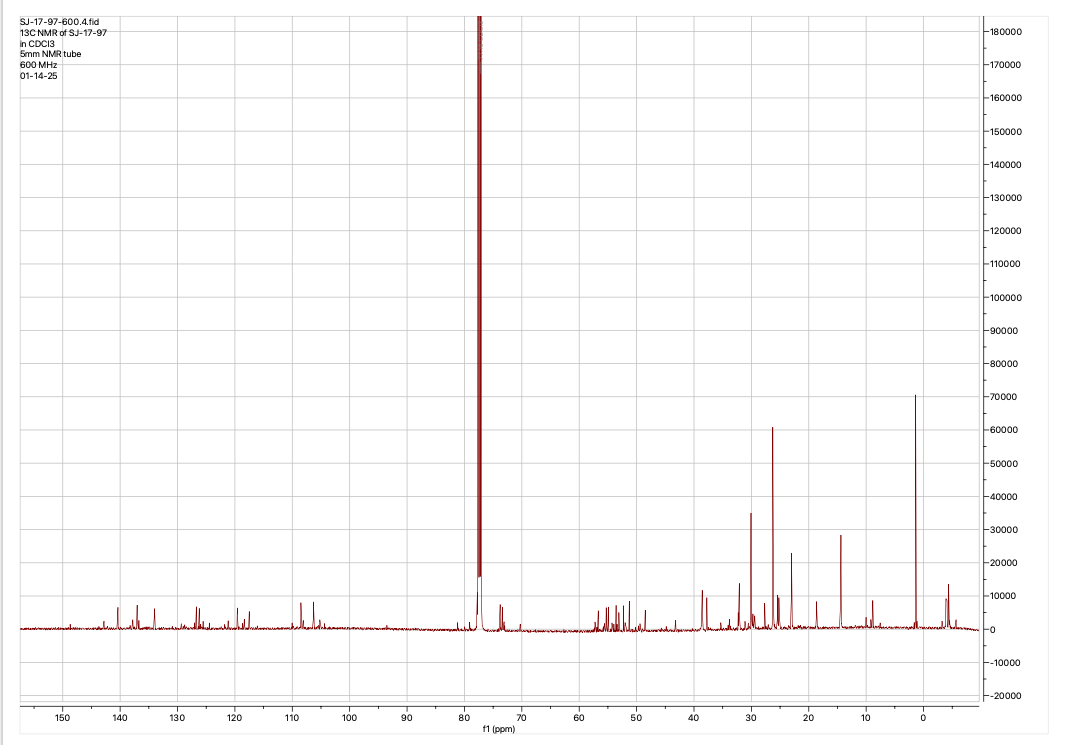

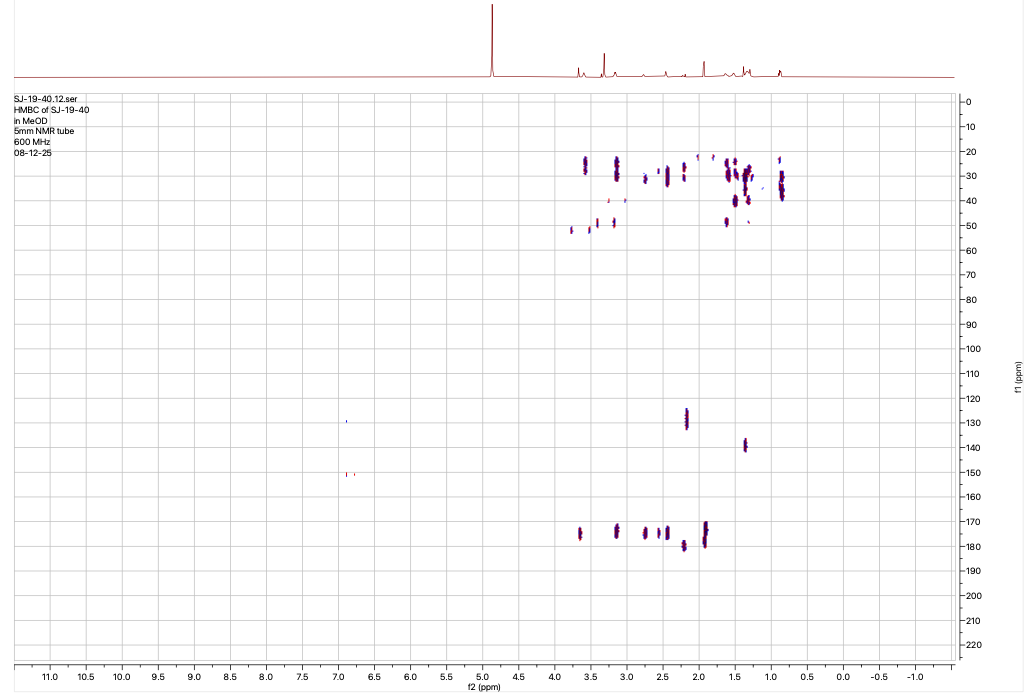

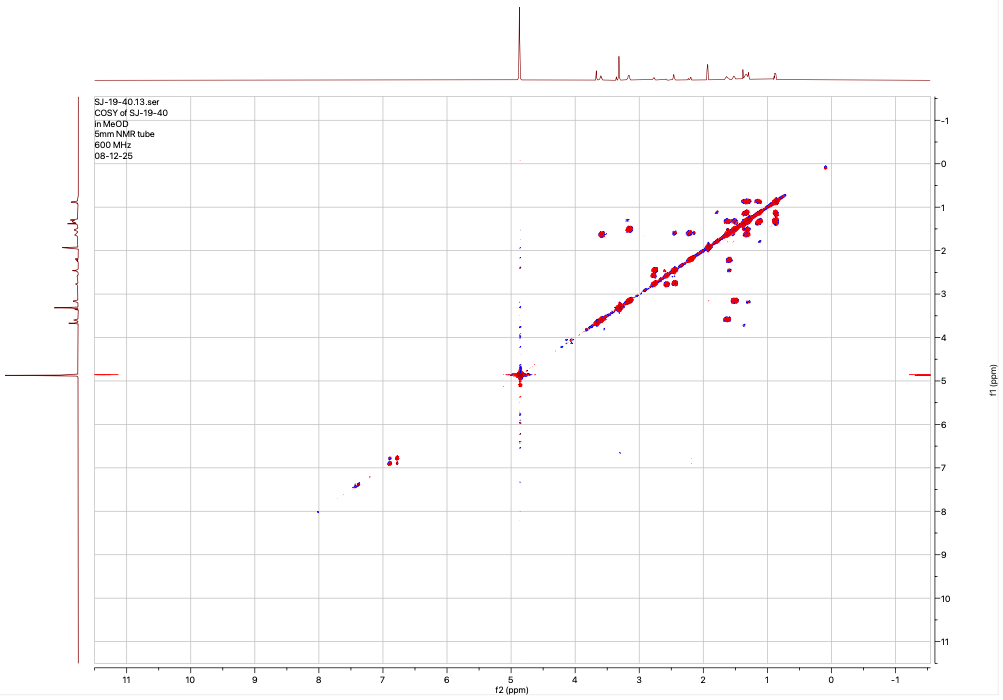

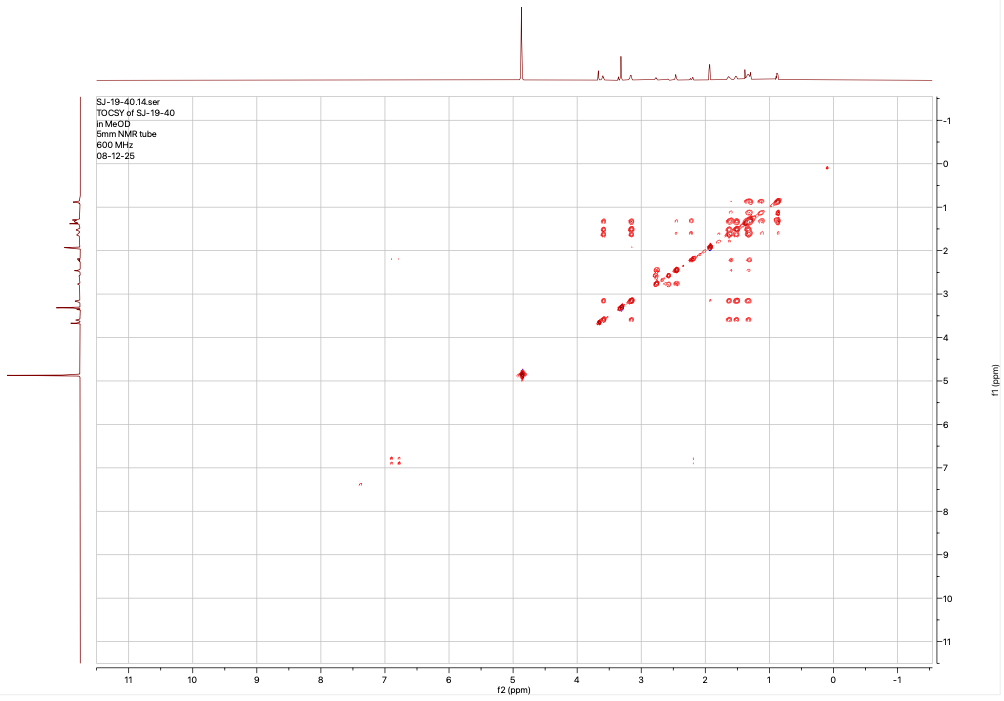

**Fig. S11.** Chemical structure along with 1D and 2D NMR characterization of the synthetic hydroxamate siderophore analog VU0980805 (Compound C9). The annotated structure includes atom numbering that corresponds to the assigned ¹H and ¹³C NMR chemical shifts listed in the table above. The dataset also includes HSQC, HMBC, COSY, and TOCSY 2D NMR spectra for synthetic compound C9

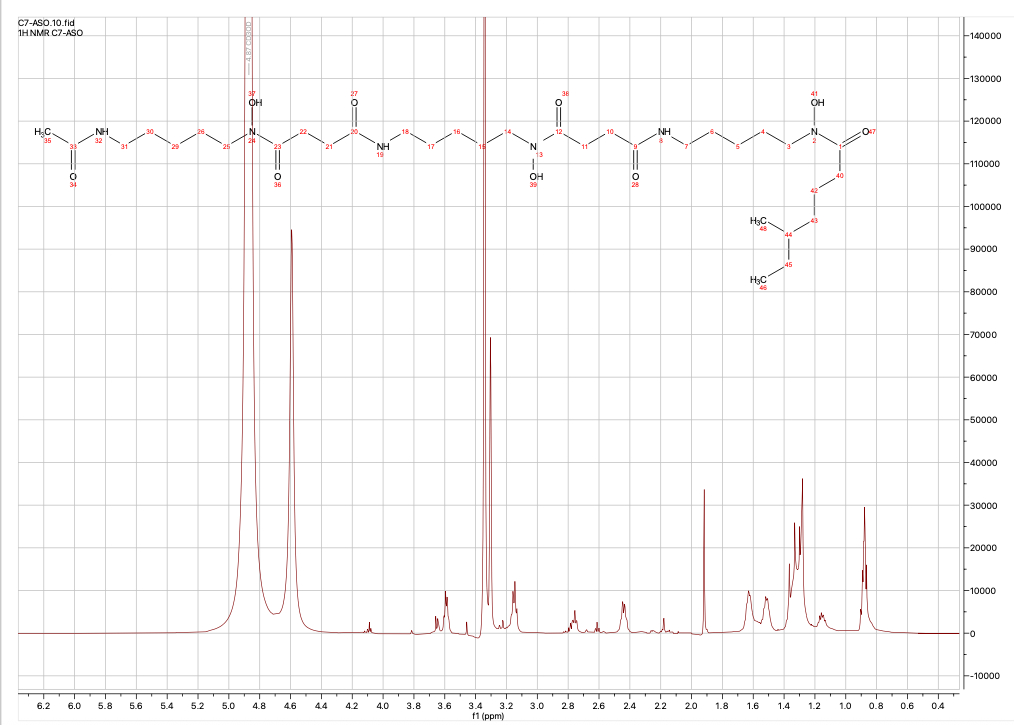

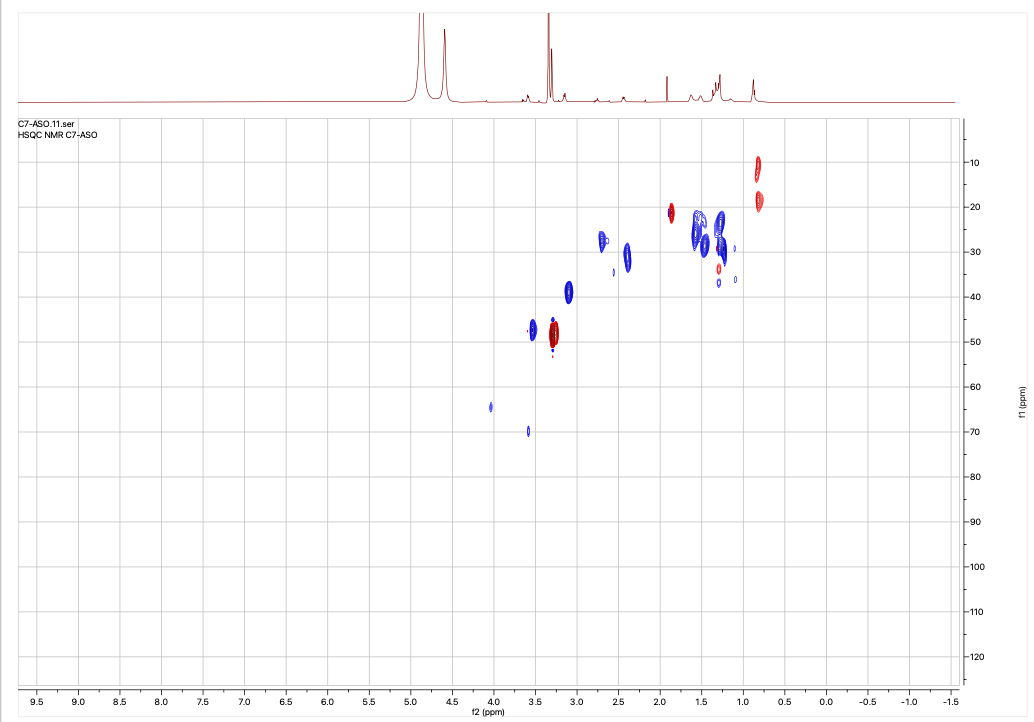
Fig. S12. illustrates the NMR data for the compound C7, both proton and HSQC 2D NMR.
